# Mechanical cues of extracellular matrix determine tumor innervation

**DOI:** 10.1101/2024.03.25.586535

**Authors:** Shu-Heng Jiang, Shan Zhang, Zhiwei Cai, Min-Hao Yu, Hui Li, Luju Jiang, Shuqi Cai, Yuheng Zhu, Hao Wang, Rui-Xue Huo, Xiang Xia, Hong-Fei Yao, Lei Zhu, Xue-Li Zhang, Li-Peng Hu, Qing Li, Jun Li, Yan-Miao Huo, Rong Hua, Junli Xue, Chongyi Jiang, Yong-Wei Sun, Jun-Feng Zhang, Zi-Zhen Zhang, De-Jun Liu, Gary Gui-Shan Xiao, Zhi-Gang Zhang

**Author notes:** These authors contributed equally to this work (Shu-Heng Jiang, Shan Zhang, Zhiwei Cai, Min-Hao Yu). **Correspondence** (S.H. Jiang); (Z.Z. Zhang); (D.J. Liu); (G.G. Xiao); (Z.G. Zhang).

## Abstract

Peripheral tumors can establish local autonomic and sensory nerve networks, termed as tumor innervation (TIN), to support tumorigenesis and metastasis. While nerve dependence in cancers is well-established, the mechanisms governing TIN remain unclear. Here, we report that extracellular matrix (ECM) stiffness, a major mechanical abnormality in the tumor microenvironment (TME), is an essential contributor of TIN. In preclinical models, reducing lysyl oxidase-mediated ECM crosslinking lowers tissue stiffness and TIN in pancreatic cancer, while inflammation-induced matrix stiffening boosts the hyperinnervation of the pancreatic precursor lesions. Mechanistically, β1-containing integrins sense the mechanical cues exerted by ECM stiffness, and the translational co-activator YAP1 acts as an essential nuclear relay to induce the expression of neurotropic genes, particularly brain-derived neurotrophic factor (BDNF) and nerve growth factor (NGF). 3D imaging of the whole cleared pancreas reveals that blockade of mechanosensor integrin β1 or pharmacological inhibition of the mechanotransducer YAP1 effectively reduces TIN. In clinical settings, tumor samples with a dense, crosslinked, and stiffened ECM exhibit significant TIN. In summary, these findings identify ECM stiffness as an important driver of TIN and suggest that targeting integrin β1/YAP1-dependent mechanotransduction may counteract TIN.

## Introduction

Cancer neuroscience is a burgeoning field that highlights the interactions between the nervous system and cancers^1^. Tumor-infiltrating nerves have become recognized as a crucial element of the tumor microenvironment (TME). Targeting tumor innervation (TIN), either autonomic nerves or sensory nerves, through genetic, chemical, or surgical methods, affects (mostly suppresses but also promotes) tumor initiation and progression and shows cancer-specific effects, suggesting that nerves are not innocent “bystander” but active “player” in the pathogenesis of cancer^2, 3, 4, 5, 6^. Analogous to the normal developmental or regeneration process, cancer is known to attract nerves to fuel tumorigenesis^7^. Recent studies support that neurotrophic factors, axon guidance molecules (AGMs), and small extracellular vesicles are mediators for nerve recruitment in cancer^8, 9, 10^.

Extracellular matrix (ECM), as the major non-cellular component of the TME, is a scaffold that supports cancer cell expansion via both biochemical and biophysical mechanisms^11^. Stiffness is a prominent property of the ECM. Excessive ECM synthesis, deposition, and crosslinking lead to matrix stiffness, which confers mechanical cues to regulate diverse malignant phenotypes, such as reprogrammed cellular metabolism^12^, epithelial-mesenchymal transition^13^, therapeutic resistance^14^, and immune escape^15^, through mechanotransduction. Reduction of ECM stiffness and targeting mechanosensors or mechanotransducers with clinically actionable drugs may help to improve cancer therapy^16, 17^. While accumulated studies have shown that matrix stiffness gives rise to cancer aggressiveness, little is appreciated about the effects of stiffening matrix on TIN.

In this work, we leveraged whether reducing or increasing ECM stiffness affects TIN in the genetically engineered mouse models of pancreatic cancer and its precancerous lesions, and found that ECM stiffness is a pronounced inducer for TIN. Then, we identified integrin-YAP1-dependent mechanotransduction that drives the ECM stiffness-induced neurotropic effects. Through a large-scale analysis of clinical specimens, we revealed a close connection between ECM stiffness and TIN in pancreatic, gastric, and colorectal cancer. Overall, this study presents a new perspective on the mechanism of TIN, highlighting that ECM mechanical properties can alter the secretome of cancer cells to enable TIN, and thus offering new avenues for understanding and targeting TIN in cancer therapy.

## RESULTS

### Inhibition of LOX-mediated crosslinking reduces tissue stiffness and TIN in a murine model of pancreatic cancer

Pancreatic innervation is characterized by significant alterations from normal pancreas to malignant tumors, particularly increased neural density (ND) and neural hypertrophy (NH) (Supplementary Fig. 1)^18^. As pancreatic ductal adenocarcinoma (PDAC) is a prime example of nerves being embedded in the desmoplastic stroma, we employed PDAC as a model to examine the potential impact of reducing matrix stiffness on TIN. The well-documented genetically engineered mouse model, KPC (*LSL-Kras^G12D/+^*; *LSL-Trp53^R172H/+^*; *Pdx1-Cre*), faithfully recapitulates the pathological alterations identified in human PDAC, especially the desmoplastic reaction. Indeed, we noticed an incremental stiffening of the pancreas as it transitioned from normal pancreas to pancreatic intraepithelial neoplasia (PanIN) and invasive PDAC (Supplementary Fig. 2A). The lysyl oxidase (LOX) family proteins encode extracellular copper-dependent amine oxidase that catalyzes the first step in the cross-linking of ECM components, including collagens and elastin^19^. LOX inhibition blocks collagen cross-linking in several preclinical models^20, 21^. Here, we generated a LOX blocking antibody that only inhibits extracellular LOX activity. KPC mice aged 10 weeks were employed since they have extensive advanced pancreatic neoplasia at this timepoint^22^. Two different strategies of LOX inhibition were performed: 1) early phase intervention, KPC mice were treated with anti-LOX for 4 weeks, followed by vehicle PBS for another 4 weeks and then sacrificed; 2) late phase intervention, KPC mice were first vehicle PBS and continued with a 4-week period anti-LOX treatment (Fig. 1A). LOX inhibition, at either early or late phase, effectively diminished serum LOX activity in KPC mice (Fig. 1B). Collagen is the most abundant ECM scaffolding protein in the stroma and the parallel fiber bundles formed by collagen fibers are essential for providing the major tensile resistance to mechanical forces^11^. Compared to control group, early LOX inhibition had mild effects on the overall collagen matrix abundance (Trichrome) but drastically reduced the width, length and straightness of collagen fibers (polarized picrosirius red) in KPC mice (Fig. 1C-E). Scanning electron microscope (SEM) showed that the frequency of tightly attached, intertwined, and thick collagen fibers was substantially reduced by early LOX inhibition (Fig. 1E, 1F). To strengthen the robustness of this finding, we performed an atomic force microscopy (AFM) analysis of the elastic modulus (stiffness) in fresh PDAC tissues and found that LOX inhibition attenuated the elastic modulus in early-phase disease but not late-stage disease, which already presents significant levels of cross-linked collagen (Fig. 1G).

**Figure 1.**
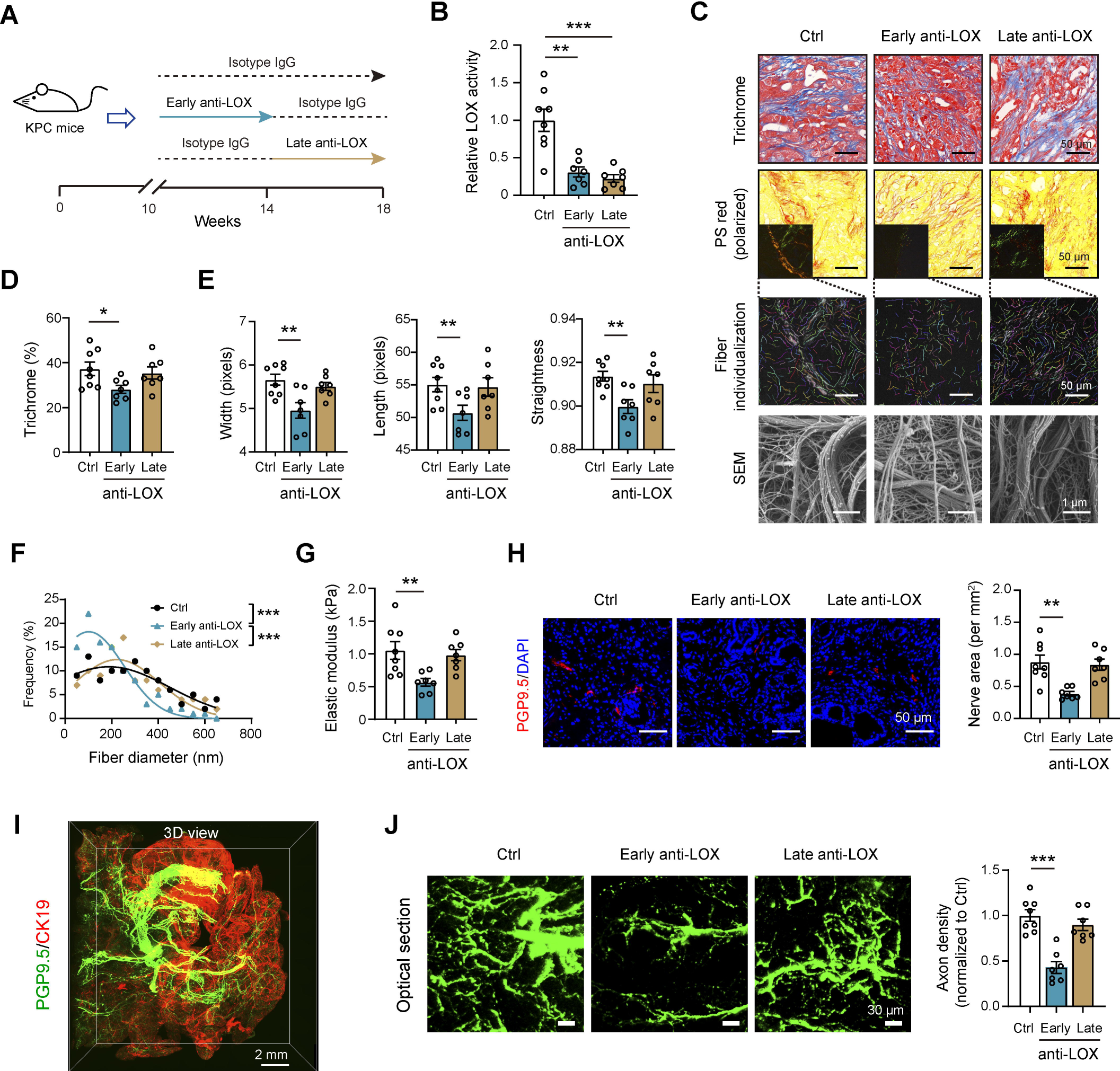
Reducing ECM stiffness inhibits TIN in a murine model of PDAC. (**A**) Experimental design of anti-LOX treatment in KPC mice. (**B**) LOX activity after early anti-LOX and late anti-LOX treatment. (**C**) Representative histology images, obtained by Trichrome stain, picrosirius red (including polarized images and collagen fiber individualization analysis), and scanning electron microscope (SEM). Scale bar, 1 and 50 μm as indicated. (**D**) Quantification of Trichrome (collagen content). (**E**) Quantification of collagen fiber width, length, and straightness. (**F**) Frequency distribution of collagen fiber diameter (selected from 5 fields of SEM images). (**G**) Young’s modulus of fresh tumor samples, measured by atomic force microscopy. (**H**) Representative images of PGP9.5^+^ nerve fibers in tumor tissues (left panel; scale bar, 50 μm). Quantification of PGP9.5^+^ nerve area (right panel). (**I**) 3D images of a pancreas from KPC mice labeled with anti-PGP9.5 and anti-CK19. (**J**) Axon projections in PDAC and PanIN lesions used for quantification of nerve densities. Scale bar, 30 μm. In **A**-**G**, n = 8 for control, n = 7 for early anti-LOX, and n = 7 for late anti-LOX. \**P* < 0.05, \*\**P* < 0.01 and \*\*\**P* < 0.001. Values as mean ± SD and compared by the one-way ANOVA multiple comparisons with Tukey’s method among groups.

In alignment with a reduction in stroma stiffness, inhibition of LOX activity early but not late resulted in less nerve area in KPC tumors, as demonstrated by PGP9.5^+^ nerves (Fig. 1H). Because pancreatic innervation is very heterogeneous and intra-visceral axons less than 5 μm are not easily detected, there are likely deviations in TIN measured in 2D tissue sections. To assess TIN more accurately, we employed a three-dimensional (3D) imaging of optically cleared tissues to visualize TIN (Fig. 1I). Compared with control or late LOX inhibition, nerve fibers at the PDAC and PanIN-afflicted regions upon early LOX inhibition had a grossly retarded branching pattern (Fig. 1J). Specifically, denervated mice display a significant reduction in the PanIN-related fibroinflammatory stroma^23^. As collagen deposition is decreased by LOX inhibition, it is likely a positive feedback loop between TIN and ECM stiffness. Taken together, these results show inhibition of LOX-mediated ECM crosslinking is effective in weakening ECM stiffness and attenuating TIN.

By analyzing the tumor and stromal components, we found that inhibition of LOX activity did not substantially affect histological differentiation and the proliferative capacity or apoptosis of tumor epithelial cells (Supplementary Fig. 2B-D). Despite a moderate increase in the number of CD31^+^ vessels and the populations of overall intratumoral infiltration of CD45^+^ cells as well as CD8^+^ T cells in KPC mice, neither early-phrase nor late-phrase LOX inhibition delayed PDAC malignant transformation (Supplementary Fig. 2E-H), suggesting that stromal or tumor alterations induced by LOX inhibition, either pro-tumorigenic or anti-tumorigenic, might be counterbalanced to affect PDAC biology^24^.

### Inflammation-induced matrix stiffness promotes hyperinnervation of pancreatic precursor lesions

Axonal sprouting responses are present in chronic inflammatory diseases of the pancreas, skin, joints, and mucosa^25, 26, 27^. Chronic pancreatitis (CP) is characterized by significant tissue stiffness, and increased neural density and enlarged nerve trunks are frequently observed in human CP^28^, especially in the extensive fibrotic areas (Supplementary Fig. 3A, 3B). Importantly, tissue extracts of CP can stimulate neurite outgrowth of dorsal root ganglion (DRG) and enteric neurons^29^. Therefore, we surveyed whether CP cooperates with the Kras mutation to boost innervation of the pancreas. To achieve this, we generated a CP model in *LSL-Kras^G12D/+^*; *Pdx1-Cre* (KC) mice and their *LSL-Kras^G12D/+^* littermates (Control), aged 24 weeks (Fig. 2A). Treatment of cerulein for 4 weeks led to a mild pancreatitis that lacks significant ECM deposition and fibrillar collagen in control mice (Fig. 2B-D). In contrast, cerulein exacerbated pancreatitis phenotypes in KC mice, producing exuberant desmoplatic reactions (Supplementary Fig. 3C, 3D). Secretion of matrix crosslinking enzymes, like LOX and transglutaminase 2 (TGM2), by the tumor cells and the cancer-associated fibroblasts (CAFs) significantly reinforces tissue stiffness^30^. Indeed, LOX and TGM2 levels were upregulated by cerulein treatment in KC mice (Supplementary Fig. 3C, 3E, 3F). In alignment with this, AFM showed that KC mice but not the control mice had incremental tissue stiffness upon cerulein treatment (Fig. 2F). Different from the observation in human CP samples, no significant fibrosis and hyperinnervation of the pancreas were noticed in CP mice (Fig. 2B, 2G). However, cerulein-mediated CP substantially amplified pancreatic innervation in the KC mice (Fig. 2G). 3D imaging of the whole pancreas also revealed that cerulein increased hyperinnervation of the PanIN lesions in the KC mice (Fig. 2H). Taken together, these findings suggest that inflammation-induced ECM stiffness induce the innervation of premalignant disease.

**Figure 2.**
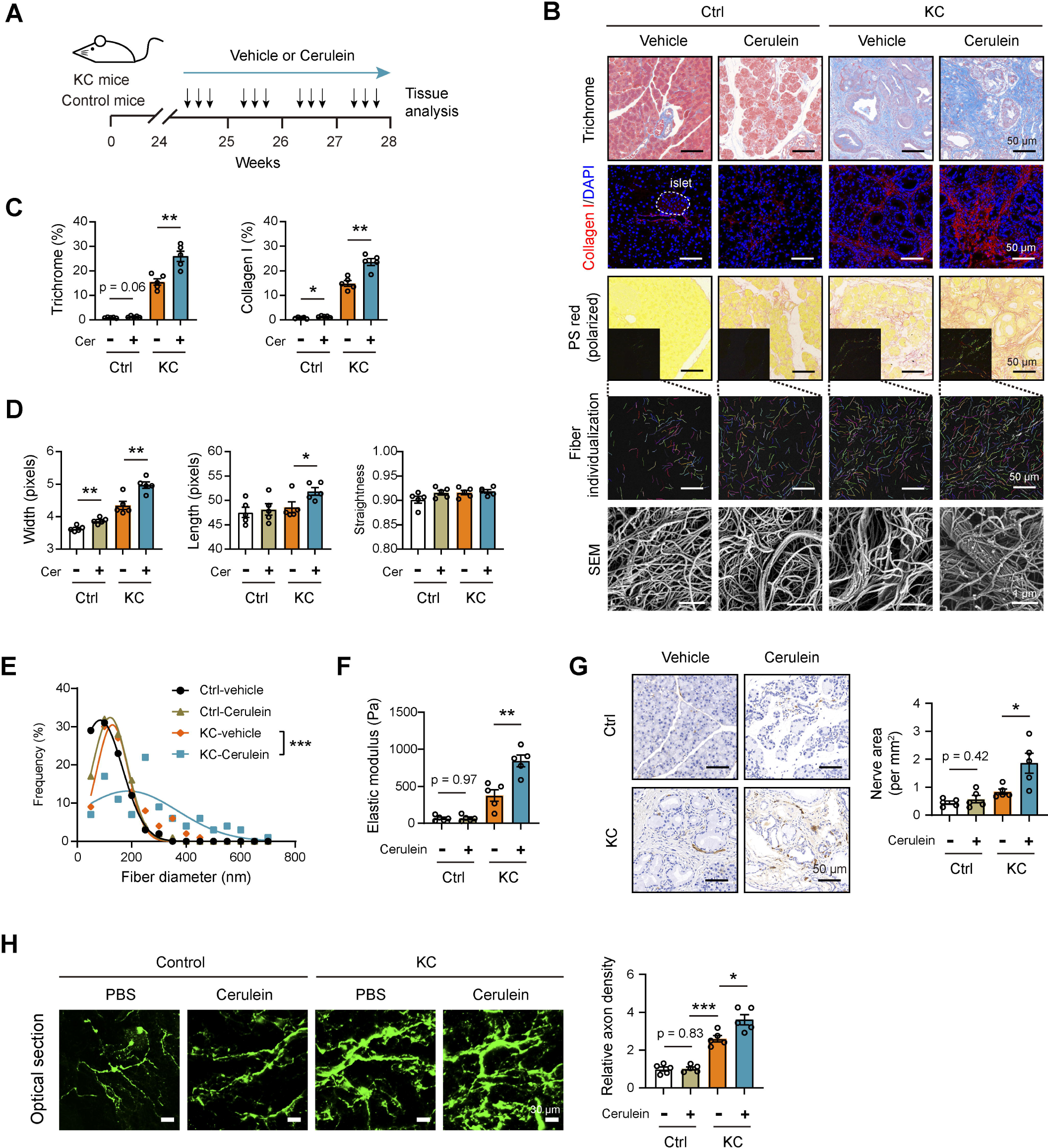
Increasing ECM stiffness promotes hyperinnervation of pancreatic precursor lesions. (**A**) Experimental design of cerulein treatment in KC and littermate control mice. (**B**) Representative histology images, obtained by Trichrome stain, collagen I IF stain, picrosirius red (including polarized images and collagen fiber individualization analysis), and scanning electron microscope (SEM). Scale bar, 1 and 50 μm as indicated. (**C**) Quantification of Trichrome (collagen content) and collagen I stain in tumor samples from indicated groups. (**D**) Quantification of collagen fiber width, length, and straightness in tumor samples from indicated groups. (**E**) Frequency distribution of collagen fiber diameter in tumor samples from indicated groups, selected from 5 fields of SEM images. (**F**) Young’s modulus of fresh tumor samples from indicated groups, measured by atomic force microscopy. (**G**) PGP9.5^+^ nerve fibers in tumor tissues from indicated groups. Scale bar, 50 μm. (**H**) Axon projections in PanIN lesions used for quantification of nerve densities in indicated groups. Scale bar, 30 μm. In **A**-**G**, n = 5 per group. \**P* < 0.05, \*\**P* < 0.01 and \*\*\**P* < 0.001. Values as mean ± SD and compared by the one-way ANOVA multiple comparisons with Tukey’s method among groups.

### Increased ECM stiffness induces neurotrophic effects

To decode the mechanism by which ECM stiffness induces TIN, we compared the bulk transcriptome in PDAC samples with higher elastic modulus (> 10 kPa) and lower elastic modulus (< 3 kPa) (Fig. 3A). Gene Ontology (GO) analyses of the differentially expressed genes (DEGs) yielded many ECM-related biological processes, including extracellular matrix organization, fibrinolysis, and cell-matrix interaction, suggesting that the analysis was built on the meaning context of matrix stiffness (Fig. 3B). Specifically, many neural categories were also significantly enriched, such as peripheral nervous system neuron development, positive regulation of neuron differentiation, innervation (Fig. 3B). The neurotrophins (NTs) and AGMs are the major drivers of TIN in human cancers^31^. Tumor cells are shown to recruit nerves, presumably by secreting NTs/AGMs encompassing nerve growth factor (NGF)^32, 33^, brain-derived neurotrophic factor (BDNF)^34^, SEMA3D^9^, and EphrinB1^10^. Indeed, PDAC samples with higher stiffness had marked changes in the expression of NTs/AGMs, either upregulated attractive guidance factors (BDNF, SEMA3C, and SEMA3D) or decreased repulsive cues (SEMA3G and Slit1) (Fig. 3C)^35, 36, 37^. Interestingly, two neural signatures, neuroactive ligand receptor interaction and axon guidance, were found in the annotation of TIN-associated DEGs, which we identified in the analysis of the TCGA cohort (Supplementary Fig. 4A). Therefore, we hypothesized that ECM stiffness induces the expression of NFs/AGMs to facilitate TIN. To test this, we seeded patient-derived PDAC cells (PDC0034) on stiff (12 kPa) and soft (0.5 kPa) type I collagen-coated matrices, which match the natural elasticities of PDAC tissues and softer normal tissues. RNA sequencing analysis revealed that PDC0034 cells under stiff cultures presented robust proneural transcriptional changes (Fig. 3D). Of the major NTs/AGMs induced by matrix stiffness, *BDNF* and *SEMA3C* were two of the most upregulated genes, and this finding was validated in PDC0034 cells and other cellular systems (Supplementary Fig. 4B, 4C). BDNF was hereafter selected for investigation as it is abundantly expressed in human PDAC tissues (Supplementary Fig. 4D).

**Figure 3.**
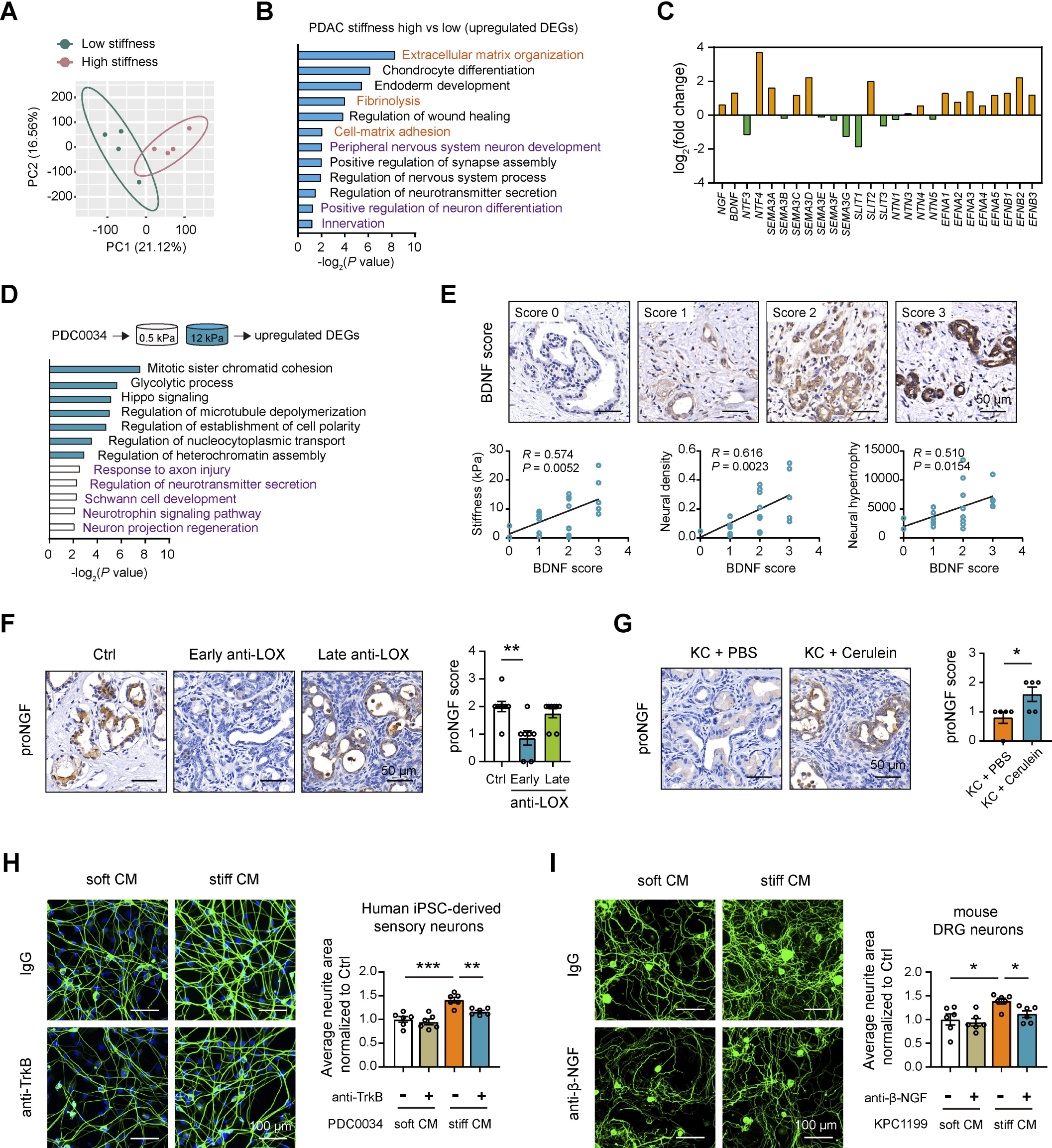
ECM stiffness induces neurotrophic effects in PDAC. (**A**) PCA plot of the RNAseq data of bulk PDAC samples with higher and lower stiffness (n = 4 per group). (**B**) KEGG annotation of upregulated DEGs as retrieved from (**A**). (**C**) Fold change of neurotrophins and axon guidance molecules in PDAC samples with higher and lower stiffness. (**D**) RNA sequencing analysis of PDC0034 cells, cultured under soft and stiff matrices (n = 3 per group). Upregulated DEGs were enriched with KEGG terms. (**E**) Correlation analysis of BDNF expression with tissue stiffness, ND, and NH in a PDAC cohort (n = 22). Scale bar, 50 μm. (**F**) proNGF expression in tumor samples from control (n = 8), early anti-LOX (n = 7) and late anti-LOX (n = 7) groups. Scale bar, 50 μm. (**G**) proNGF expression in pancreas samples from KC + PBS and KC + cerulein groups (n = 5 per group). Scale bar, 50 μm. (**H**) Neurite outgrowth of human iPSC-derived sensory neurons upon treatment with anti-TrkB and indicated CM from PDC0034 cells, which cultured under soft and stiff matrices. Scale bar, 100 μm. (**I**) Neurite outgrowth of mouse DRG neurons upon treatment with anti-β-NGF and indicated CM from KPC1199 cells, which cultured under soft and stiff matrices. Scale bar, 100 μm. \**P* < 0.05, \*\**P* < 0.01, \*\*\**P* < 0.001. Values were compared by the one-way ANOVA multiple comparisons with Tukey’s method among groups (**F**, **H**, **I**), Spearman’s rank correlation methods (**E**), and Student’s t test (**G**). Experiments were independently repeated three (**H**, **I**) times with similar results. RNA sequencing experiments (**A**, **D**) were not repeated.

To test the association of BDNF with tissue stiffness and TIN, we conducted AFM analysis and PGP9.5 staining in a cohort of PDAC samples (n = 22). Previous findings support that both localized sprouting of new branches and axonal hyperbranching of preexisting nerve bundles are the origin of nerves in pancreatic cancer^38^. Therefore, we considered both ND and NH for TIN analysis. ND was calculated by determining the total number of nerves within a unit area (1 mm^2^), and NH was expressed as the ratio of the total nerve area to the total number of nerves in the analyzed section. As a result, BDNF expression was closely associated with the tissue stiffness, ND, and NH in PDAC (Fig. 3E). Different from the expression pattern of NTs/AGMs in human PDAC, NGF has the highest level in murine PDAC^8^. Indeed, *Ngf* mRNA level was markedly increased under stiff cultures in murine PDAC cells (Supplementary Fig. 4E). Moreover, early LOX inhibition was capable of reducing proNGF expression in tumor tissues from KPC mice (Fig. 3F). In contrast, cerulein treatment augmented proNGF expression in PanIN lesions (Fig. 3G). To determine the link between stiffness-induced neurotrophic effects and TIN, we stimulated human induced pluripotent stem cell (iPSC)-derived sensory neurons with conditioned medium (CM) from PDC0034 cells. Compared to the CM from soft culture condition (soft CM), stiff CM efficiently promoted neurite outgrowth of iPSC-derived sensory neurons; moreover, this effect was massively curbed by neutralizing TrkB, the functional receptor for BDNF (Fig. 3H). Likewise, stiff CM from KPC-derived PDAC cells (KPC1199) increased the neurite outgrowth of murine DRG neurons compared with soft CM, and anti-β-NGF treatment abrogated this effect by 70% (Fig. 3I). Altogether, these findings bolster that ECM stiffness promotes the secretion of NTs/AGMs to induce TIN.

Outside of the source of cancer cells, neurotrophic factors might be derived from stromal cells within the TME, such as immune cells and CAFs. This notion is especially motivated by the hyperinnervation in CP, as the substantial expansion of stromal fibroblasts during CP development and inflammatory score is closely associated with increased neural density in CP tissues^39^. Moreover, hyperinnervation appears as early as the PanIN stage in mice^23^, when the pancreas stiffness is not significant (Supplementary Fig. 2A). Indeed, data from single-cell RNA sequencing analysis showed the diverse sources of NFs/AGMs in the TME of human PDAC and PanIN-afflicted donor pancreas, particularly ductal cancer cells, fibroblasts, and endothelial cells (Supplementary Fig. 4F, 4G). Therefore, ECM stiffness might not be the inducer of TIN during early tumorigenesis but a driver of TIN with the progress of the disease.

### YAP1 is responsible for ECM stiffness-induced neurotrophic effects

The matrix signals are eventually transmitted to the nucleus, which induces functional output through transcriptional mechanisms^40^. Mass spectrometry analysis of the nuclear lysates of PDC0034 cells, which were harvested under stiff and soft culture conditions, revealed 492 upregulated and 221 downregulated proteins (Table. S1). These differentially expressed proteins (DEPs) were functionally involved in peptidyl-proline hydroxylation, aerobic electron transport chain, maintenance of DNA methylation, mitotic spindle elongation, and peripheral nervous system development (Fig. 4A). Among the DEPs (Fig. 4B), we were particularly intrigued by YAP1, which is known as a nuclear convey of mechanical signals exerted by ECM stiffness^41^. The Hippo pathway responds to extracellular environments and regulates the subcellular localization of the transcriptional co-activator YAP1^42^. When PDC0034 cells were cultured under stiff matrices, immunoblot showed prominent nuclear YAP1 localization (Fig. 4C), and YAP1 transcriptional activity was enhanced as evidenced by the expression of the YAP1 target genes (*CTGF* and *CYR61*) (Fig. 4D) and a YAP/TAZ-responsive luciferase reporter (8×GTIIC-luc) (Fig. 4E). In preclinical animal models, early LOX inhibition reduced nuclear YAP1 levels in tumors of KPC mice, while opposite results were found in PanIN lesions of KC mice upon cerulein treatment (Supplementary Fig. 5A, 5B). In human PDAC samples, nuclear expression of YAP1 was closely associated with tissue stiffness, ND, and NH (Supplementary Fig. 5C). Pharmacological inhibition of YAP1 with verteporfin, which mitigates the YAP1-TEAD1-4 interaction, prevented PDC0034 cells from acquiring a neurotropic signature under stiff culture condition (Fig. 4F) and strongly lowered the secretion of BDNF (Fig. 4G). In contrast, XMU-MP-1, an agonist of YAP1, increased BDNF secretion toward high stiffness values under soft culture conditions (Fig. 4H). Using chromatin immunoprecipitation (ChIP), we discovered that YAP1 occupancy at the promoters of *BDNF* genes was elevated under stiff culture conditions compared to soft culture conditions, and this effect was reversed by the addition of verteporfin (Supplementary Fig. 5D), reflecting YAP1-dependent control of BDNF expression.

**Figure 4.**
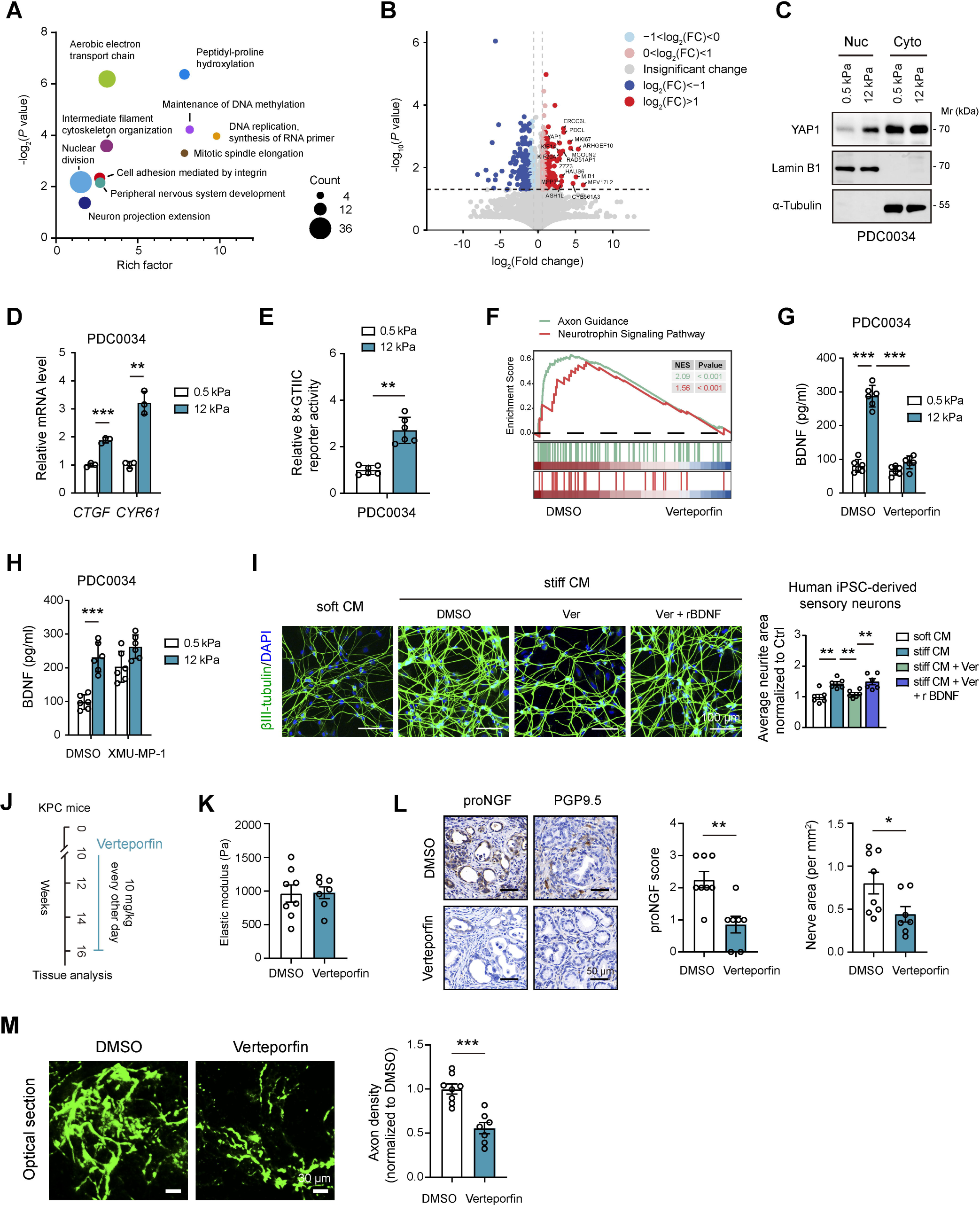
YAP1-dependent mechanotransduction induces the neurotrophic effects. (**A**) Mass spectrometry analysis of the nuclear lysates of PDC0034 cells, cultured under soft and stiff matrices (n = 3 per group). Differentially expressed proteins (DEPs) were enriched with GO biological process terms. (**B**) Volcano plot of DEPs as retrieved from (**A**). (**C**) Western blot of YAP1 in the nuclear and cytoplasmic lysates. Lamin B1 and α-Tubulin were loaded as the nuclear and cytoplasmic control, respectively. (**D**) mRNA level of *CTGF* and *CYR61* in PDC0034 cells, cultured under soft and stiff matrices (n = 3). (**E**) 8×GTIIC reporter activity in PDC0034 cells, cultured under soft and stiff matrices (n = 6). (**F**) Gene set enrichment analysis (GSEA) plot of axon guidance and neurotrophin signaling pathway between DMSO and verteporfin group. RNA sequencing was performed with PDC0034 cells cultured under stiff conditions. (**G**) Enzyme-linked immunosorbent assay (ELISA) of BDNF level in the supernatants of PDC0034 cells upon verteporfin treatment (n = 6). (**H**) ELISA of BDNF level in the supernatants of PDC0034 cells upon XMU-MP-1 treatment (n = 6). (**I**) Neurite outgrowth of human iPSC-derived sensory neurons upon treatment with rBDNF and indicated CM from PDC0034 cells, which cultured under soft and stiff matrices (n = 6). Scale bar, 100 μm. (**J**) Experimental design of verteporfin treatment in KPC mice. (**K**) Young’s modulus of fresh tumor samples from control and verteporfin groups, measured by atomic force microscopy. (**L**) proNGF expression and PGP9.5^+^ nerve fibers in tumor tissues from control and verteporfin groups. Scale bar, 50 μm. (**M**) Axon projections in PDAC and PanIN lesions used for quantification of nerve densities. Scale bar, 30 μm. In **J**-**L**, n = 8 for vehicle, n = 7 for verteporfin treatment. \**P* < 0.05, \*\**P* < 0.01, \*\*\**P* < 0.001. Values were compared by the one-way ANOVA multiple comparisons with Tukey’s method among groups (**D**, **G**, **H**, **I**) and the Student’s t test (**E**, **K**, **L, M**). Experiments were independently repeated two (**I**) or three (**C**-**E**, **G**, **H**) times with similar results. RNA sequencing experiment in **F** was not repeated.

Functionally, the neurite outgrowth of iPSC-derived sensory neurons induced by stiff CM was not seen in stiff CM from PDC0034 cells preincubated with verteporfin, but it could be rescued significantly (though not completely) by replenishment of recombinant BDNF (Fig. 4I). The close link between matrix stiffness and YAP1-dependent neurotropic effects was also found in murine PDAC cells (Supplementary Fig. 5E-I). To extend this link to an *in vivo* setting, we assessed YAP1 inhibition in KPC mice (Fig. 4J). Verteporfin dampened the expression of proNGF, while without significant influences on the tissue stiffness and YAP1 levels in cancer cells (Fig. 4K). Importantly, verteporfin drastically reduced the infiltration of PGP9.5^+^ nerve fibers compared to the control group, as revealed by 2D tissue sections and 3D imaging analysis (Fig. 4L, 4M). Collectively, these findings suggest that YAP1 is a required mediator of the neurotrophic effects controlled by matrix stiffness.

### Integrins relay mechanical signal from ECM stiffness to YAP1 in TIN

Next, we probed the cell surface mechanosensors that engage in outside-in signaling. Integrins, especially β1-containing integrins, transduce cues from the ECM by assembling adhesion plaque complexes that initiate biochemical signaling and stimulate cytoskeletal remodeling to regulate cell behavior^43^. The force increases integrin expression, activity, and focal adhesions. Tumor tissues, particularly PDAC, are often fibrotic and stiff, and PDAC cells that exhibit high tension have elevated focal adhesions and increased integrin signaling^44^. In line with this notion, integrin activation and focal adhesion assembly were enriched in the analysis of PDAC samples with high and low tissue stiffness (Supplementary Fig. 6A). Motivated by the highly expressed integrin β1 in PDAC tissues (Supplementary Fig. 6B)^45^, we determined whether integrin β1 senses substrate stiffness. Inactivation of integrin β1 with a blocking antibody reduced YAP1 nuclear translocation (Fig. 5A) and robustly inhibited YAP1-dependent transcriptional activity (Fig. 5B). Consistently, the DEGs induced by anti-ITGB1 treatment, either downregulated or upregulated, had higher similarity in stiffness-responsive transcriptome with that in cells upon YAP1 inhibition (Fig. 5C). Likewise, anti-ITGB1 largely reversed the neurotropic gene signatures, resembling changes found in verteporfin treatment (Fig. 5D). Moreover, integrin β1 blocking attenuated stiffness-induced neurotropic effects in YAP WT but not YAP1 knockdown or dominant-negative (YAP1 S94A) cells, as assayed by BDNF secretion and neurite outgrowth (Fig. 5E-G and Supplementary Fig. 6C, 6D). Inhibition or knockdown of focal adhesion kinase (FAK) or p130Cas, two integrin adhesion plaque proteins^46^, consensually impaired YAP1 activation and the neurotropic effects in cells cultured under conditions of high stiffness (Supplementary Fig. 6E-G). Conversely, PDC0034 cells expressing autoclustered integrin β1 (β1 V737N) phenocopied high stiffness responses on soft gels (Fig. 5H-J), emphasizing integrin β1-mediated signal transduction. Then, we asked whether integrin β1 could sense ECM stiffness to regulate YAP1 and TIN *in vivo*, by using the KPC model (Fig. 5K). As a result, blocking integrin β1 had no impact on the elastic modulus of PDAC tissues (Fig. 5L), but effectively lowered the nuclear abundance of YAP1 and proNGF expression in PDAC cells (Fig. 5M). Moreover, 3D imaging of the whole pancreas revealed a significant reduction of TIN in KPC mice with integrin β1 blocking (Fig. 5N).

**Figure 5.**
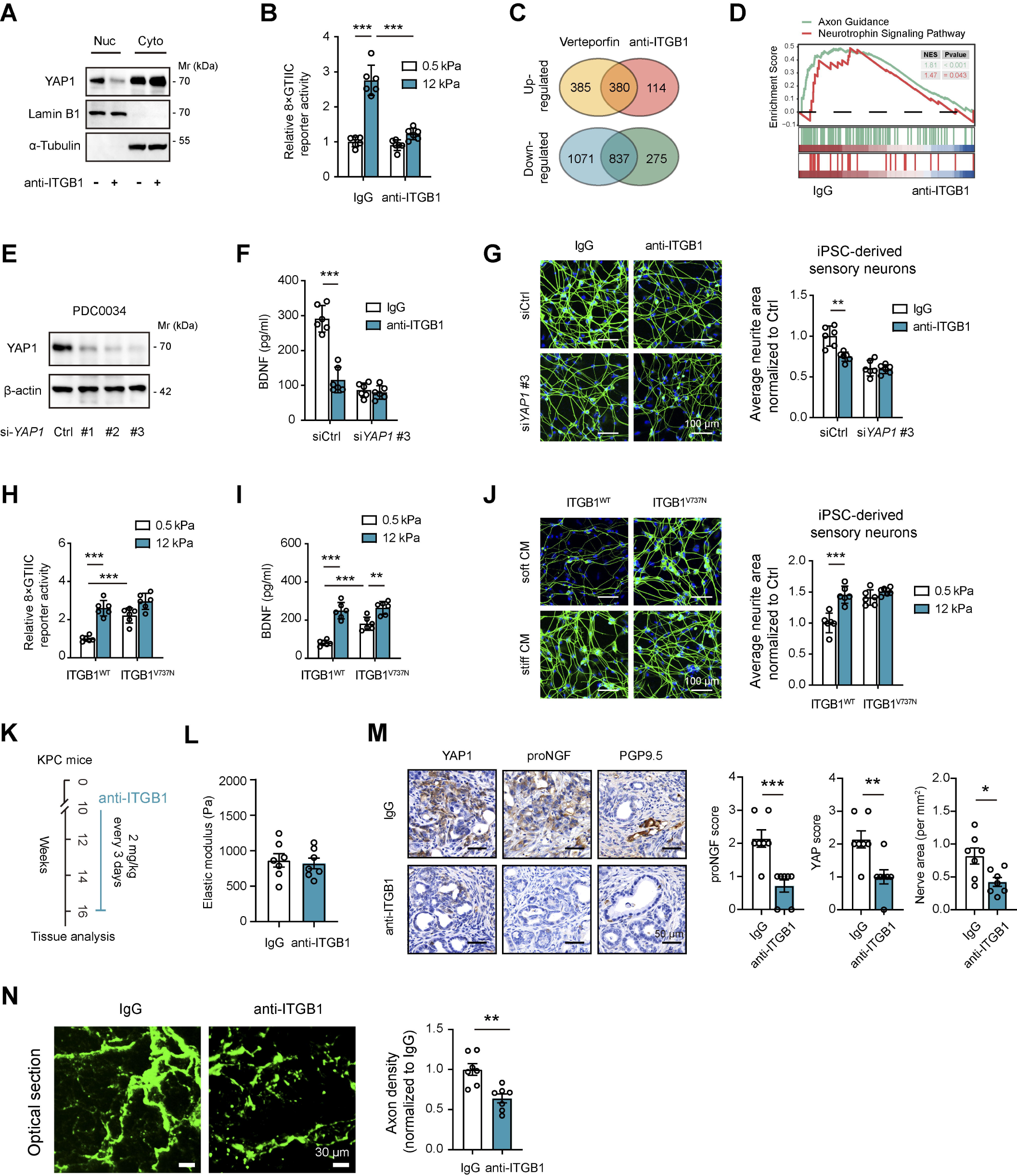
Integrins sense matrix stiffness and transduce signal cascades. (**A**) Western blot of YAP1 in the nuclear and cytoplasmic lysates of PDC0034 cells upon anti-ITGB1 treatment. Lamin B1 and α-Tubulin were loaded as the nuclear and cytoplasmic control, respectively. (**B**) Effects of anti-ITGB1 on the 8×GTIIC reporter activity in PDC0034 cells, cultured under soft and stiff matrices (n = 6). (**C**) Comparison of up-regulated and down-regulated genes in PDC0034 cells upon verteporfin and anti-ITGB1 treatment. (**D**) GSEA plot of axon guidance and neurotrophin signaling pathway between IgG and anti-ITGB1 group. (**E**) siRNA-mediated *YAP1* knockdown in PDC0034 cells. (**F**) Effects of anti-ITGB1 on the secretion of BDNF by siCtrl and si*YAP1*-#3 PDC0034 cells (n = 6). (**G**) Neurite outgrowth of human iPSC-derived sensory neurons upon treatment with CM from siCtrl and si*YAP1*-#3 PDC0034 cells, which were cultured under stiff conditions and treated with or without anti-ITGB1. Scale bar, 100 μm. (**H**) 8×GTIIC reporter activity in ITGB1^WT^ and ITGB1^V737N^ PDC0034 cells, cultured under soft and stiff conditions (n = 6). (**I**) BDNF secretion by ITGB1^WT^ and ITGB1^V737N^ PDC0034 cells, cultured under soft and stiff matrices (n = 6). (**J**) Neurite outgrowth of human iPSC-derived sensory neurons upon treatment with CM from ITGB1^WT^ and ITGB1^V737N^ PDC0034 cells, which were cultured under soft and stiff conditions. Scale bar, 100 μm. (**K**) Experimental design of anti-ITGB1 treatment in KPC mice. (**L**) Young’s modulus of fresh tumor samples from IgG and anti-ITGB1 groups, measured by atomic force microscopy. (**M**) YAP1, proNGF expression, and PGP9.5^+^ nerve fibers in tumor tissues from IgG and anti-ITGB1 groups. Scale bar, 50 μm. (**N**) Axon projections in PDAC and PanIN lesions used for quantification of nerve densities. Scale bar, 30 μm. In **K**-**M**, n = 7 per group. \**P* < 0.05, \*\**P* < 0.01, \*\*\**P* < 0.001. Values were compared by the one-way ANOVA multiple comparisons with Tukey’s method among groups (**B**, **F**-**J**, **H**, **I**) and the Student’s t test (**L**, **M**, **N**). Experiments were independently repeated two (**G**, **J**) or three (**A**, **B**, **F**, **H**, **I**) times with similar results. RNA sequencing experiment in **C** was not repeated.

Besides integrins, stiffness information can be conveyed by other putative receptors or channels, particularly Piezo1^47, 48^, DDR1^49^, and CD44^50^. Molecular silencing of these receptors by siRNAs minimally or partially blunted YAP1 activation, the expression of neurotropic genes, and neurite outgrowth of iPSC-derived sensory neurons. In contrast, the knockdown of integrin β1 abolished most of the changes in the stiffness-induced signaling and neurotropic effects (Supplementary Fig. 6H-K). Collectively, integrins may couple the matrix stiffness to intracellular YAP1 signaling pathways, which enable the development of TIN.

### ECM stiffness is linked to TIN in clinical settings

To aid the clinical relevance, we examined the connection between ECM stiffness and TIN in two independent cohorts. First, in the cohort mentioned in Fig. 3E (n = 22), we found that the average elastic modulus of PDAC tissues had a close correlation with both ND and NH (Fig. 6A, 6B). As second line of evidence, TIN was assayed by co-immunofluorescence staining of S100β (Schwann cell marker) and NF-L (a nerve marker) in another PDAC cohort (n = 74) (Fig. 6C). Quantitative analysis revealed that samples with high neural density (ND-H) had overall collagen matrix abundance compared to samples with low neural density (ND-L), as evidenced by Trichrome staining and collagen I expression (Fig. 6C, 6D). Compared with ND-L cases, ND-H cases showed higher expression of matrix crosslinking enzymes, LOX and TGM2 (Fig. 6C, 6D). Moreover, a major cause of ECM stiffening is the increased contractile forces generated by α-SMA stress fibers, which are produced by myofibroblasts^30^. Analogously, higher α-SMA activity was noticed in ND-L samples when compared to the ND-H samples (Fig. 6C, 6D). Picrosirius red staining and polarized microscopy found that ND-H samples were associated with wide, long, and aligned collagen fibers (Fig. 6C, 6E). Furthermore, we compared the connection between matrix stiffness and NH and found similar results (Supplementary Fig. 7A-C). Because pancreatic innervation is heterogeneous with anatomical regions^51^ and nerve distribution tends to decrease toward tumor core^52^, we further analyzed ND nor NH categorized by tumor location (pancreas head versus body/tail) and tumor size. Notably, neither ND nor NH correlated with tumor size, but ND-H was more frequently found in tumors located in the head of the pancreas (Fig. 6F, 6G and Supplementary Fig. 7D, 7E).

**Figure 6.**
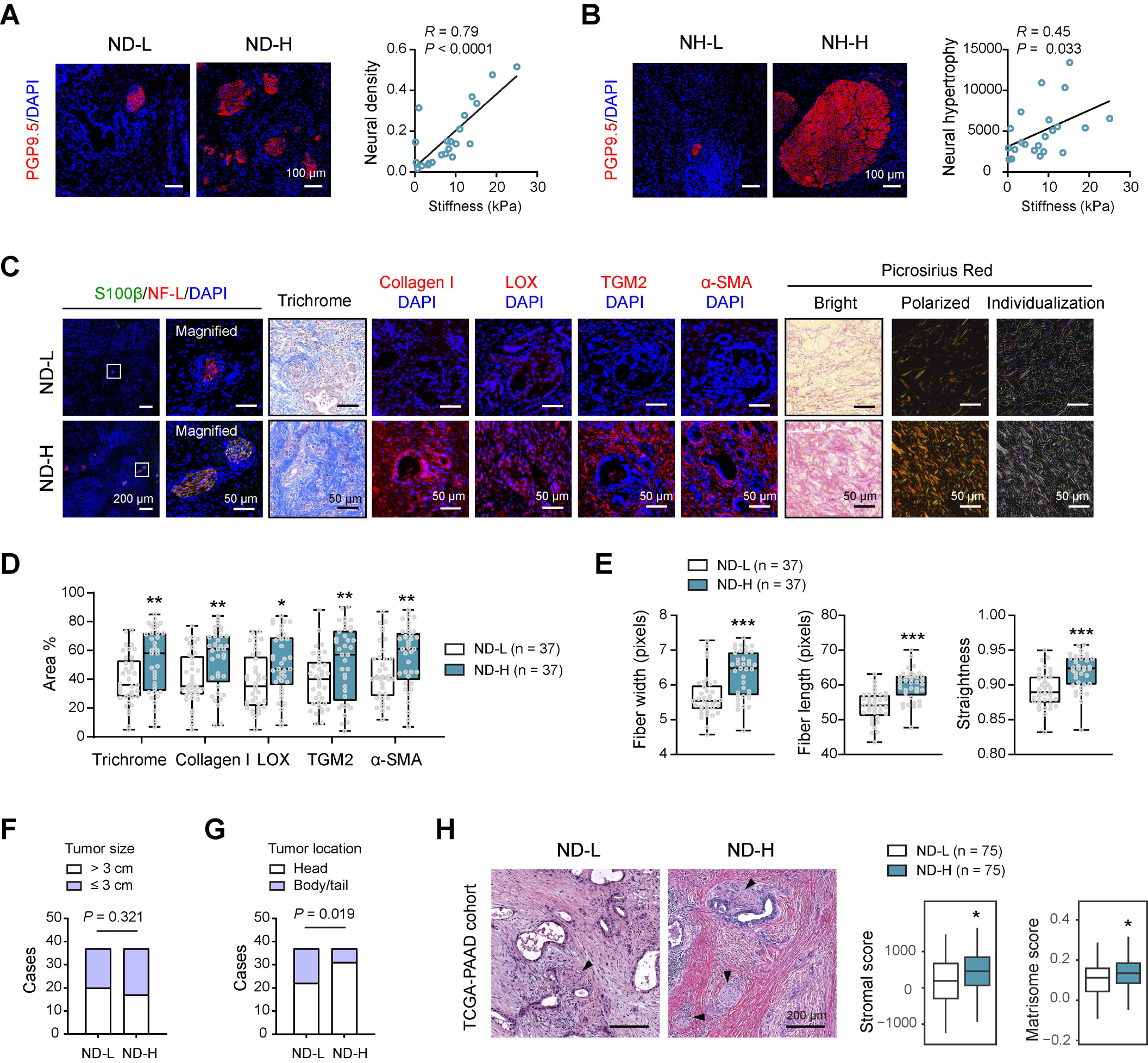
ECM stiffness is linked to TIN in PDAC. (**A**) Representative images of ND-L and ND-H samples, as displayed by PGP9.5 IF stain. Correlation analysis of tissue stiffness with ND (n = 22, right panel). Scale bar, 100 μm. (**B**) Representative images of NH-L and NH-H samples, as displayed by PGP9.5 IF stain. Correlation analysis of tissue stiffness with NH (n = 22, right panel). Scale bar, 100 μm. (**C**) Representative histology images of ND-L and ND-H samples, obtained by co-immunofluorescence (IF) analysis of S100β and NF-L, Trichrome stain, collagen I IF stain, LOX IF stain, TGM2 IF stain, α-SMA IF stain, and picrosirius red (including polarized images and collagen fiber individualization analysis). Scale bar, 50 and 200 μm as indicated. (**D**) Quantification of Trichrome (collagen content), collagen I area, LOX area, TGM2 area, and α-SMA area between ND-L and ND-H samples (n = 37 per group). (**E**) Collagen fiber width, length, and straightness in ND-L and ND-H samples (n = 37 per group). (**F**) The association between ND and tumor size in PDAC (n = 37 per group). (**G**) The association between ND and tumor location in PDAC (n = 37 per group). (**H**) Stromal score and matrisome score of ND-L and ND-H cases in the TCGA cohort (n = 150). The arrow heads indicate nerves. Scale bar, 200 μm. \**P* < 0.05, \*\**P* < 0.01 and \*\*\**P* < 0.001. Values were compared by the Spearman’s rank correlation methods (**A**, **B**), Student’s t test (**D**, **E**, **H**), and the chi-square test (**F**, **G**). ND, neural density; NH, neural hypertrophy.

In The Cancer Genome Atlas (TCGA) cohort, we quantified the ND status of PDAC samples using the whole-slide H&E diagnostic images and combined corresponding bulk RNAseq data of each sample for differential analysis (n = 150). Strikingly, samples with ND-H had higher matrix activity compared with samples with ND-L, as demonstrated by higher stromal score^53^ and the expression of matrisome genes (glycoproteins, collagens, and proteoglycans) (Fig. 6H).

Such a connection between stiff matrix and neural density is not specific to PDAC but also found in tumors of the highly innervated gastrointestinal tract, as seen in stomach adenocarcinoma (STAD, Renji cohort, n = 158; TCGA cohort, n = 369) and colorectal cancer (CRC, Renji cohort, n = 144; TCGA cohort, n = 375) (Supplementary Fig. 8 and 9). Collectively, these findings indicate that differences in a dense, crosslinked, and stiffened ECM may translate into differences in TIN across cancer types.

## Discussion

Infiltrating nerves are emerged as a conspicuous component in the TME, but the underlying mechanisms of innervation remain largely unclear. This study identifies a previously unprecedented link between ECM stiffness and TIN and shows that integrins-YAP1-mediated mechanotransduction is required for the expression of neurotropic genes, which enable TIN (Supplementary Fig. 10).

Our findings have several important implications. First, we provide important insights into the mechanism of TIN and illuminate that TIN is largely driven by the mechanic property of the TME, stiffness. This further broadens a previously unrecognized function of ECM within the TME. Our cross-cancer analysis suggests that TIN is especially evident in the cancer types with rich and stiffened stroma, particularly PDAC. Consistently, significant TIN is present in highly desmoplastic prostate cancer and cholangiocarcinoma^54, 55^. Moreover, the normal pancreas and colorectal tissue are abundant in innervation, but the presence of tumor-infiltrating nerves in colorectal cancer is notably lower compared to that in PDAC. Certainly, neurotrophic factors (NGF, Artemin, and GAP-43) are significantly less potent in colon cancer than in PDAC^56^. Considering the higher stiffness of PDAC compared to colorectal cancer, this variance may be attributed to ECM stiffness. One may wonder about deficiency of TIN in liver cancer, where fibrosis is recognized as a key player in cancer biology. A possible explanation is the low elastic modulus of primary liver tumors (less than 1 kPa)^57^. Conflicting findings are also encountered in breast cancer, which presents higher tissue elastic modulus (10.4-42.52 kPa)^58^ but lesser frequency of TIN (18%-36.93%)^59, 60^, raising that TIN may emerge via common and distinct signaling mechanisms in different types of cancers.

Second, our study proposes that a stiffened matrix stimulates integrin-mediated mechanotransduction to drive the expression of neurotropic factors, which further induces neurite outgrowth and innervation. We delineated integrin β1/YAP1 as a central hub in response to ECM stiffness and showed that inactivation of either integrin β1 or YAP1, stiff matrices are ineffective in inducing TIN in PDAC, suggesting the fundamental integrin sensing as a mechanism for TIN. Neurotrophic growth factors are profoundly implicated in the initiation of TIN. We highlighted the source of neurotrophic factors from cancer cells but did not exclude the contributions from other stromal cells (e.g., fibroblasts and immune cells). Indeed, emerging roles of fibroblasts and macrophages have been reported in neurite outgrowth and neural remodeling^61, 62^.

Third, our study emphasizes the importance of considering cancer-specific TIN patterns for nerve-targeting therapy. Accumulated studies have demonstrated that TIN is associated with malignant phenotypes and correlates with poor clinical outcomes. Given the tumor-promoting role of sensory nerves^63^, tumor-suppressive parasympathetic nerves^64^, and contradictory roles of sympathetic nerves^38, 65^ in PDAC, the remodeling changes of intratumoral nerves may dictate their actual contribution. As ECM is highly dynamic and undergoes remodeling during the establishment of a mature TME^66^, TIN status may also fluctuate in a context-dependent manner. Therefore, whether cutting off TIN can achieve clinical therapeutic effect depends on their function within the TME and may be related to different cancer types and disease stages. From the therapeutic point of view, our results open up avenues for indirect targeting TIN, such as stiffness, integrin signaling, YAP1, and neurotropic factors. Moreover, drug repurposing of clinical neurologic drugs might represent a way to treat tumors. Whether targeting these components alone or in combination with chemotherapy or radiotherapy provided long-term control of tumors warrants further investigation.

To distill, we describe the mechanistic understanding of ECM stiffness-induced TIN. This model applies not only to cancer but perhaps also to other inflammatory diseases, where significant demoplasia is present. Considering that targeting TIN has a limited therapeutic index in cancer, targeting innervation may be applicable to diseases with abundant nerve infiltration, such as CP and Crohn’s disease.

### Limitations of the study

We studied matrix stiffness with cells cultured in 2D, on top of soft and stiff elastic substrates. However, cancer cells interact with ECM in a 3D environment, and the mechanism underlying mechanotransduction in 3D is different from that in 2D^67^. Thus, whether stiffness-mediated integrin clustering and YAP1 activation converge on the nucleus, to 3D context remains to be investigated. Our model implies that cancer cells are the critical source of BDNF and NGF in the TIN of PDAC. Although cutting-edge studies support this model, unequivocal evidence to clarify the cell-specific loss of these neurotrophic factors will be more convincing.

## Methods

### Patient population

Two cohorts, Renji cohort and the TCGA cohort, were included for analysis. Clinical specimens for immunohistochemical staining, including samples of PDAC (n = 74), STAD (n = 158), and CRC (n = 144), were obtained from Ren Ji Hospital, School of Medicine, Shanghai Jiao Tong University. For atomic force microscope analysis, fresh tumor samples from 22 PDAC patients, 21 STAD patients, and 20 CRC patients were studied. Additionally, normal pancreas (NP, n = 20) and chronic pancreatitis (n = 8) were included. NP tissues were all pancreatic body tissues obtained from patients with pancreatic tail tumors. The patients included in the study were provided with detailed information regarding their age, gender, diagnosis, tumor location, disease stage, and histological differentiation. All protocols involving the use of human materials were approved by the ethics committee of Ren Ji Hospital, School of Medicine, Shanghai Jiao Tong University (approval number RA-2021-095, KY2021-120-B). Prior to participation, written informed consent was obtained from all patients. In the TCGA cohort, whole-slide tissue hematoxylin and eosin (H&E) diagnostic images, clinical information, and molecular data of PDAC (n = 150), STAD (n = 369), and CRC (n = 375) were obtained from the Genomic Data Commons (GDC) Legacy Archive (https://portal.gdc.cancer.gov/).

### Histological evaluation of tumor innervation

Intratumor nerves in tissue sections of PDAC, STAD, and CRC were immunolabeled using the neuronal marker NF-L and Schwann cell marker S100β. In NP and CP, the nerves were immunolabeled with PGP9.5. To quantify neural density and neural hypertrophy, the entire tissue sections were scanned at high magnification and reconstructed into a mosaic image using the 3DHISTECH CaseViewer2.3 (RRID: SCR_017654). In cases with endoneural invasion of cancer cells, only the nerve trunk without cancer cell invasion was included in the analysis. Nerve fibers were counted regardless of the nerve size, nerve shape and cut directions of the nerves. Contiguous nerve segments were counted as one nerve trunk. All nerves larger than approximately 10 μm in diameter were identified in the tumor bulk, while tiny nerve axons were not counted. The number of nerves and the nerve area in the entire tissue sections were marked with the CaseViewer program. For the TCGA cohort, the H&E diagnostic images were downloaded in the original image format (Aperio SVS files) and analyzed with Aperio ImageScope 12.3.3 (RRID: SCR_020993). Neural density was calculated as the total number of nerves within a unit area (1 mm^2^) of the investigated specimen. Accordingly, neural hypertrophy was expressed as the ratio of the total measured area of the nerves to the total number of nerves in the analyzed section. A logistic regression model then was constructed using neural density and neural hypertrophy as the dichotomized-dependent variables, and fitting model variables of age, sex, tumor location, and tumor size.

### H&E, picrosirius red, Masson’s trichrome, and alcian blue staining

For all histological staining, 5 μm paraffin-embedded tissue sections were deparaffinized and rehydrated using a series of xylene and alcohol washes and then subjected to H&E staining (C0105M, Beyotime), picrosirius red staining (G1470, Solarbio), Masson’s trichrome staining (ab150686, Abcam), and alcian blue staining (G1027, Servicebio) according to the manufacturer’s instructions. For picrosirius red staining, the stained sections were serially imaged using the Olympus fluorescence microscope with the analyzer and polarizer oriented parallel and orthogonal to each other. Under polarized light microscopy, the stained collagen fibers appear in different colors depending on their thickness and organization. Thick collagen type I fibers appear red or orange, while thin collagen type III fibers appear green or yellow against a black background. Collagen fiber properties were analyzed with the CT-FIRE fiber analysis software (http://loci.wisc.edu/software/ctfire). For fiber metrics of width, length, and straightness, the mean values were averaged across two to three ROIs for each sample. For evaluation of collagen with Masson’s trichrome staining, nuclei and other basophilic structures appear blue-black, cytoplasm and muscle fibers appear red, and collagen fibers appear blue. Alcian blue staining was used to detect acidic mucins in murine PDAC tissues. The alcian blue-stained acidic polysaccharides appear as blue or bluish-green coloration.

### Immunohistochemical (IHC) staining

IHC analysis was routinely performed as reported previously^68^. The following primary antibodies was used: PGP9.5 (14730-1-AP, Proteintech, 1:200), Ki67 (28074-1-AP, Proteintech, 1:1,000), cleaved caspase 3 (#9661, Cell Signaling Technology, 1:400), CD45 (ab40763, Abcam, 1:250), CD8A (ab217344, Abcam, 1:1,000), CD31 (28083-1-AP, Proteintech, 1:2,000), BDNF (ab108319, Abcam, 1:500), proNGF (ab52918, Abcam, 1:250), Sema3C (ab135842, Abcam, 1:100), YAP1 (#14074, Cell Signaling Technology, 1:300). Immunoreactivity was generated with diaminobenzidine under a phase-contrast microscope. The sections were then dehydrated, mounted, and observed under the microscope. For IHC scoring, if the staining intensity is equal to the normal adjacent tissue, the specimen is considered negative and scored as 0; staining intensity greater than the control normal adjacent pancreatic tissue but limited to less than 25% of the cells in a tissue section, it is scored as weak and assigned a score of 1; staining intensity greater than the control and present in less than 50% of the cells, it is scored as moderate and assigned a score of 2; and staining intensity is greater than the control and present in more than 50% of the cells, it is scored as strong and assigned a score of 3.

### Immunofluorescence (IF) analysis

Paraffin-embedded tissue sections were deparaffinized, rehydrated with graded ethanol, and subjected to antigen retrieval. After blocking with 10% (m/v) BSA (A500023-0100, Sangon Biotech) for 1 h at room temperature, the tissue sections were incubated with primary antibodies at 4 °C overnight. Primary antibodies were S100β (66616-1-Ig, Proteintech, 1:200), NF-L (ab223343, Abcam, 1:100), Collagen I (#72026, Cell Signaling Technology, 1:200), LOX (17958-1-AP, Proteintech, 1:200), TGM2 (15100-1-AP, Proteintech, 1:200), α-SMA (A5228, Sigma-Aldrich, 1:400), PGP9.5 (14730-1-AP, Proteintech, 1:200), PGP9.5 (66230-1-Ig, Proteintech, 1:500; for co-IF analysis), CK19 (1:200, Proteintech, 10712-1-AP), and AMY2A (15845-1-AP, Proteintech, 1:100). The next day, the tissue sections were incubated with species-specific secondary antibodies (ThermoFisher Scientific, USA) for 45 min at room temperature. Finally, nuclei were counterstained with 4’,6-diamidino-2-phenylindole (DAPI; AppliChem, Cat. A4099) and the stained sections were imaged with confocal microscopes (Nikon, Japan).

### Atomic force microscopy (AFM)

Tumor tissues were dissected followed by incubation using phosphate-buffered saline (PBS) and immobilized to the surface of a 35 mm dish with an adhesive glue. The samples were then imaged using an atomic force microscope (FastScan Bio, Bruker, USA) mounted on an inverted microscope (Nikon Eclipse-Ti). Silicon nitride cantilevers with a spring constant of 0.15 N/m were used, which were attached to a borosilicate glass spherical tip with a diameter of 5 µm. The cantilevers were tapped on the tumor tissues and five 15 µm × 15 µm AFM stiffness maps were acquired for each sample. To calculate the elastic modulus, the Hertz model for a spherical tip was applied to fit the experimental indentation curves.

### Scanning electron microscopy

Harvested pancreas tissues were sliced to a thickness of 1 mm and fixed with 2.5% glutaraldehyde (Servicebio, G1102) for 2 h. All samples were post-fixed 1% osmium tetroxide (Polysciences, Cat. 23311-10, USA) for 2 h and dehydrated in graded concentration of alcohols. After drying with a critical point dryer (Quorum Technologies Ltd, K850, UK), the samples were attached to the double-sided conductive carbon tape and put on the sample table of ion sputtering instrument (IXRF, MSP-2S, USA) for gold spraying for about 30 sec. Finally, the samples were viewed under an Emission Scanning Electron Microscope (SEM; hitachi, SU8010, Japan).

### Mice

LSL-Kras^G12D/+^; LSL-Trp53^R172H/+^; Pdx1-Cre (referred to as KPC) mice and LSL-Kras^G12D^; Pdx1-Cre (referred to as KC) were described previously^69^. For the KPC study, LOX function-blocking antibody (3 mg/kg; purified; twice a week; SinoBiological, China), anti-ITGB1 (1 μg/g; twice a week; eBioscience, Cat. 16-0291-85), IgG isotype antibody (1 μg/g; twice a week; eBioscience, Cat. 14-4888-85) or Verteporfin (20 mg/kg; daily; Selleck, Cat. S1786), was intraperitoneally (i.p.) administered. LOX function-blocking antibody treatment was continued for 4 weeks and started at two different time points: the early disease stage (around 10 weeks of age) and the late disease stage (around 14 weeks of age). The anti-ITGB1 treatment was started at 10 weeks of age and continued for 6 weeks. In the chronic pancreatitis model, KC mice and their littermates were given six hourly i.p. injections of 50 μg/kg body weight cerulein (Selleck, Cat. S9690) three times per week for a total of 4 weeks. Experimental mice with desired genotypes were used, and both female and male mice were included. There was no randomization or blinding during the monitoring and analysis of the mice. Mice were housed in specific pathogen-free conditions and maintained on a 12-hour light-dark cycle at a temperature of 23 ± 1 °C and a relative humidity range of 40-75%. All mice were fed standard chow ad libitum and provided with water throughout the experiment. All animal procedures were reviewed and approved reviewed and approved by the Research Ethics Committee of Shanghai Jiao Tong University (approval number 202201427).

### 3D imaging of the whole cleared pancreas

Intact pancreas tissues were subjected to immunostaining and clearing using the FDISCO protocol^70^, which enables superior fluorescence preservation. Briefly, mice were anesthetized by intraperitoneal injection of a mixture of 10% urethane (Selleck, Cat. S4544), 2% chloral hydrate (Merck, Cat. C8383) along with 0.3% xylazine (MedChemExpress, Cat. HY-B0443A). Thereafter, mice were intracardiacally perfused with 20 ml of PBS followed by 20 ml of 4% paraformaldehyde (PFA) in PBS. Harvested pancreas tissues were postfixed overnight at 4 °C in 4% PFA, rinsed several times with PBS, placed in 50 and 80% methanol (in PBS) for 1 h at each step and twice in 100% methanol for 1 h, and bleached with 5% H_2_O_2_ in 20% DMSO/methanol at 4°C overnight. Before immunostaining, samples were incubated with 0.2% Triton X-100/20% DMSO/0.3 M glycine at 37 °C overnight and blocked in 6% goat serum/0.2% Triton X-100/10% DMSO at 37°C for 2 days. Samples were then incubated with primary antibodies at 37 °C for 4 days, washed in PBS, and incubated with secondary antibodies for 4 days at 37 °C. The following antibodies were used: primary antibody, PGP9.5 (Proteintech, Cat. 14730-1-AP) and CK19 (Abcam, Cat. ab7754); secondary antibodies, donkey anti-rabbit Alexa Fluor 594 (Jackson ImmunoResearch, Cat. 711-585-152) and donkey anti-mouse Alexa Fluor 488 (Jackson ImmunoResearch, Cat. 715-545-150). DAPI (AppliChem, Cat. A4099) was used for nuclear counterstaining after incubation with the secondary antibody. After immunostaining, the tissues were cleared using a commercial tissue clearing reagent kit (Jarvis, Cat. JA11012, Wuhan, China), which includes 6 reagents labeled A-F. The labeled tissues were sequentially incubated in A, B, C, D, and E reagents for 12 h each, followed by another 12 h incubation in E reagent. Then, the sample was incubated in F reagent until it became transparent. All the clearing steps were performed at 6-8°C with slight shaking. To clear the tissue after dehydration, pure dibenzyl ether (Sigma-Aldrich, Cat. 108014) was used as a refractive index matching solution. Finally, the tissues were imaged with LiTone XL (Light Innovation Technology, Hong Kong, China). To process the imaging data, the TIFF raw data images were first stitched and converted using LitScan software. The resulting images were analyzed using Imaris software (Version 7.2.3, Bitplane, Switzerland).

### RNA sequencing analysis

Human PDAC tissues and patient-derived cancer cells were employed for RNA sequencing analysis. The total RNA was extracted from PDAC tissues with varying stiffness and PDC0034 cells that underwent different manipulations, such as soft culture, stiff culture, YAP1 inhibition, and ITGB1 blocking, using the Trizol method. RNA quality and cDNA library quality were checked before paired indexing sequencing (Illumina NovaSeq 6000). The sequencing process was entirely controlled by data collection software provided by Illumina, and the sequencing result data was analyzed in real-time. The bulk RNA-sequencing data can be accessed from the Sequence Read Archive (SRA) repository under accession numbers PRJNA1064588 and PRJNA1064589. Gene sets were all obtained from the MSigDB database (https://www.gsea-msigdb.org/gsea/index.jsp). GSEA analysis was performed in GSEA software following instructions of the software.

### Cell culture and reagents

Human PDAC cell lines (AsPC-1 and Capan-2) were obtained from the American Type Culture Collection (ATCC, Manassas, VA, USA). Two mouse PDAC cell lines, KPC1199 and Panc02, and PDAC patient-derived PDC0034 cells were generously provided by Professor Jing Xue from Ren Ji Hospital, School of Medicine, Shanghai Jiao Tong University. Cells were cultured in RPMI-1640 medium or DMEM according to ATCC protocols and supplemented with 10% (v/v) fetal bovine serum (FBS, Cat. 30067185, Thermo Fisher Scientific, USA) and 1% antibiotics (100 μg/mL streptomycin and 100 units/mL penicillin, Cat. SB-CR011, Share-bio, China) at 37 °C in a humidified incubator under 5% CO_2_ condition. All cell lines were routinely tested for mycoplasma contamination and were found to be negative. The collagen I-coated soft (0.5 kPa) and stiff (12 kPa) culture plates, including 6-well plate, 12-well plate, and 6 cm plate, were all sourced from Matrigen (Irvine, CA, USA).

### Cell transfection and plasmids

To initiate the transfection process, exponentially growing untreated PDC0034 cells were first plated in 60-mm dishes for 24 h. Upon reaching 70% confluence, the cells were transfected with 50 ng of specific siRNAs targeting the indicated genes or a scrambled siRNA that did not target any known gene sequence. The Lipofectamine® RNAiMAX reagent (ThermoFisher Scientific, Cat. 13778030) was used according to the manufacturer’s instructions. The siRNA oligonucleotides were synthesized by GenePharma (Shanghai, China), and the detailed sequences of the siRNA are available in Supplementary Table 2. After 48 h of transfection, the knockdown efficiency of indicated genes was determined by immunoblot analysis or real-time qPCR, and the cells were either harvested or processed for further experiments. The plasmids encoding wild-type YAP1 (YAP1^WT^), dominant-negative YAP1 (YAP1^S94A^), wild-type ITGB1 (ITGB1^WT^), and autoclustered ITGB1 (ITGB1^V737N^) were generated by GENEray Biotechnology (Shanghai, China).

### Mouse DRG neurons and iPSC-derived sensory neurons

For isolation of mouse DRG neurons, mice were euthanized by CO_2_ asphyxiation followed by cervical dislocation. The DRGs from the thoracic areas were dissected under sterile conditions using fine forceps and scissors and transferred to a 15 ml tube containing Dulbecco’s Phosphate Buffered Saline (DPBS) solution (Hyclone, Cat. SH30028). To digest the DRG tissue, collagenase D (Sigma-Aldrich, Cat. 11088866001) and 2.5% trypsin-EDTA solution (R&D systems, Cat. B81710) were added to the tubes and incubated for 30 min at 37°C. To stop the digestion process, a complete neurobasal medium (Invitrogen, Cat. 21103049) with 2% horse serum was added to the tubes and centrifuged for 2 min at 800× g. To enrich DRG neurons, 1 ml of 35% Percoll in 0.9 M NaCl was added to the bottom of a 15 ml tube and covered with 1 ml of cell suspension. The tube was then centrifuged for 15 min at 285 × g, resulting in a reddish pellet composed of sensory neurons at the bottom of the tube. Finally, the pellet was resuspended with pre-warmed complete medium containing 50 ng/mL nerve growth factor (Sigma-Aldirch, Cat. N6009) and 20 μM 5-FU (Sigma-Aldrich, Cat. F6627). The isolated DRG neurons were seeded in 6-well culture plates at a density of 1 × 10^5^ cells per well. To minimize the nonneuronal contamination, the culture medium was changed to serum-free medium for 3-4 h after the DRG cells had adhered to the culture plate. In the process of generating iPSC-derived sensory neurons, human iPSC-derived neural crest cells are replated onto a poly-L-ornithine (PLO) solution and a laminin-coated dish. Over the initial 6 days, the sensory neuron differentiation media (Amplicongene, Cat. SN-D0001, China) is replaced daily, after which it is substituted with sensory neuron growth media (Amplicongene, Cat. SN-C0001, China). Subsequently, the sensory neuron growth media is changed every 2 days.

### Conditioned medium collection

PDC0034 and KPC1199 cells, including wild type and genetically manipulated cells (siCtrl, si*YAP1*, si*ITGB1*, si*DDR1*, si*PIEZO1*, si*CD44*, si*p130Cas*, YAP1^S94A^, and ITGB1^V737N^), were cultured in a 6-well plate under stiff and soft ECM conditions. After 24 h, the cells were washed three times with PBS and then cultured with 400 μl of fresh medium. Following treatment with verteporfin (Selleck, Cat. S1786), anti-TrkB (R&D systems, Cat. AF1494), anti-ITGB1 (R&D systems, Cat. MAB17781), or defactinib (Selleck, Cat. S7654) for 12 h, the CM was collected, centrifuged at 15,000× g for 5 minutes to remove cellular debris, and filtered through a 0.22-μm filter to ensure sterility. Mouse DRG neurons or human iPSC-derived sensory neurons were exposed to fresh standard culture medium mixed with CM at a 1:1 ratio (v/v) during their growth period.

### Neurite outgrowth assay

Mouse DRG neurons were seeded on a glass-bottom cell culture dish and cultured in neurobasal medium (Gibco, Cat. 21103049) supplemented with B-27 supplement (Gibco, Cat. 17504044), 2 mM L-glutamine (Gibco, Cat. 25030149), 10% FBS, and 1% antibiotic-antimycotic (Gibco, Cat. 15240062). Human iPSC-derived sensory neurons were cultured in dishes coated with Matrigel matrix (Corning, Cat. 356253), poly-d-lysine (Sigma-Aldrich, Cat. P4957), and laminin (Sigma-Aldrich, Cat. L2020). CM stimulation was performed daily and continued for 4 days. After the stimulation period, neurons were washed with PBS three times and fixed with 4% PFA for 30 min, followed by immunofluorescence analysis with anti-βIII tubulin antibody (Abcam, Cat. ab18207). Images were acquired using a confocal microscopy system (Leica, TSC SP8), and fields were randomly selected and imaged for subsequent analysis. The average neurite areas were quantified using Image J (version 1.54b) and normalized to the total cell numbers.

### RNA isolation, cDNA generation, and qPCR

PDAC cells were homogenized using TRIzol reagent (ThermoFisher Scientific, #15596018) to extract RNA. The isolated RNA was quantified and assessed for purity using a Nanodrop spectrophotometer. cDNA synthesis was performed using a one-step PrimeScript RT-PCR kit (Takara Bio, Cat. RR057B, Japan) as per the manufacturer’s protocols. qPCR analysis was carried out using SYBR-Green Premix Ex Taq (Bimake, Cat. B21203, China) on a ViiA7 Real-time PCR system (Applied Biosystems, USA). The detailed sequences for primers used in this study are available in Supplementary Table 3. Relative gene expression was calculated based on the 2^−ΔΔCt^ method and normalized to β-actin mRNA levels.

### Isolation of cytoplasm and nuclear protein

The cytoplasmic and nuclear proteins were isolated using a commercial kit (Share-Bio, Cat. SB-PR013HZ, China) following the manufacturer’s instructions. Briefly, cultured cells were harvested by scraping and transferred to a tube, followed by centrifugation at 500 × g for 5 min at 4°C. The cell pellet was then resuspended with ice-cold cytoplasmic extraction buffer and incubated on ice for 15 min, followed by centrifugation at 14,000 × g for 10 minutes at 4°C. The resulting supernatant constituted the cytoplasmic fraction. Subsequently, the nuclear pellet was resuspended with ice-cold nuclear extraction buffer and incubated on ice for 30 min with intermittent mixing, followed by centrifugation at 14,000 × g for 15 min at 4°C to collect the nuclear protein-containing supernatant. The protein concentrations in the cytoplasmic and nuclear fractions were determined using a Pierce BCA Protein Assay Kit (ThermoFisher Scientific, Cat. 23228). The isolated cytoplasmic and nuclear proteins were aliquoted and stored at -80°C for subsequent analysis.

### Mass spectrometric analysis

The cytoplasmic and nuclear protein samples from PDC0034 cells, cultured under both soft and stiff conditions, underwent a series of meticulous procedures for comprehensive analysis. After concentration and freeze-drying, SDT lysis buffer was added to the samples, which were then transferred to an EP tube and homogenized using a homogenizer. Subsequently, the samples were bathed in boiling water for 3 min and ultrasonically crushed for 2 min, followed by centrifugation at 16,000 × g for 20 min at 4°C. The resulting supernatant was collected, and the protein content was determined using the BCA method. DTT was added to a concentration of 100 mM in each sample, and samples were bathed in boiling water for 5 min. Subsequently, 200 µl of UA buffer (8 M urea, 150 mM Tris-HCl, pH 8.0) was added to the sample and mixed evenly. The mixture was then transferred to a 10 KD ultrafiltration centrifuge tube and centrifuged at 12,000× g for 15 min. After this step, 100 µl of IAA (50 mM IAA in UA) was added, and the mixture was shaken at 600 rpm for 1 min. It was then kept out of light for 30 min and centrifuged at 12,000 g for 10 min. Next, 100 µl of UA buffer was added to the sample, and the sample was centrifuged at 12,000× g for 10 min, which was repeated twice. Following this, 100 µl of NH_4_HCO_3_ buffer was added to the sample, and it was centrifuged at 14,000× g for 10 min, which was repeated twice. Finally, 40 µl of trypsin buffer (6 g trypsin in 40 µl NH_4_HCO_3_ buffer) was added to the sample, and the sample was shaken at 600 rpm for 1 min at 37°C for 16-18 h. The resulting hydrolyzed peptides were desalted using a C18 Cartridge and subsequently freeze-dried under vacuum. After enzymatic hydrolysis, the peptides were dried and then redissolved with 0.1% formic acid (FA), and the peptide concentration was determined for LC-MS analysis. For DIA mass spectrometry data acquisition, 9 μl of peptide fragments were extracted from each sample, and 1μl of 10× iRT (Biognosys AG) peptide fragments were added. After mixing, 2 μg samples were taken, and chromatographic separation was performed using the Easy-nLC1200 chromatographic system (ThermoFisher Scientific). Subsequently, the separated peptide was analyzed using DIA (data independent acquisition) mass spectrometry with a Q-Exactive HF-X mass spectrometer (ThermoFisher Scientific).

### Immunoblotting

Total protein from cultured cells was extracted using an IP lysis buffer (Servicebio, Cat. G2038, Wuhan, China) with protease and phosphatase inhibitors. The protein concentration was determined using a Pierce BCA Protein Assay Kit (ThermoFisher Scientific, Cat. 23228). Proteins were separated by SDS–PAGE electrophoresis and transferred onto polyvinylidene fluoride (PVDF) membranes (Beyotime, Cat. FFP33, China). The membranes were blocked with 5% (m/v) nonfat milk (Sangon, Cat. A600669-0250, China) in Tris-buffered saline (TBS)/Tween 20 for 1 hour and then incubated with primary antibodies, including YAP1 (Cell Signaling Technology, Cat. 14074, diluted at 1:1,000), Lamin B1 (Proteintech, Cat. 12987-1-AP, 1: 2,000), and α-Tubulin (Proteintech, Cat. 11224-1-AP, 1: 2,000). After washing with TBS/Tween 20 three times, the membranes were incubated with horseradish peroxidase (HRP)-linked secondary antibodies. The signals were detected using an enhanced chemiluminescent (ECL) plus reagent kit (Share-Bio, Cat. SB-WB012L, China) and viewed with a ChemiDoc MP Imaging System (Bio-Rad Laboratories).

### 8×GTIIC-luciferase activity assay

The YAP-dependent transcriptional activity was assessed using an 8xGTIIC-luciferase construct (Addgene plasmid # 34615). Briefly, cells were co-transfected with either the GTIIC plasmid or a control empty plasmid, along with 10 ng/well of phRLSV40 (Renilla expression plasmid serving as an internal transfection control). After 24 h post-transfection, the firefly and renilla luciferase activities were measured using a dual luciferase reporter assay (Promega, Cat. E1960) according to the manufacturer’s protocols. Relative firefly luciferase activity was normalized to Renilla activity.

### Chromatin immunoprecipitation

PDC0034 cells, cultured under soft or stiff conditions in the presence or absence of verteporfin, were subjected to chromatin immunoprecipitation (ChIP) analysis. The ChIP assay was performed using control IgG (Cell Signaling Technology, Cat. 3900) or anti-YAP1 antibody (Cell Signaling Technology, Cat. 14074) with the MAGnify Chromatin Immunoprecipitation System (ThermoFisher Scientific, Cat. 492024) following the standard protocol. Following the antibody pulling down, the target DNA was resuspended in water for qPCR analysis. Primers were designed within YAP1 binding sites at the BDNF promoter region. The known YAP1 target, CTGF, was used as a positive control, and Hemoglobin beta (HBB) served as a negative control. The primers used for ChIP-PCR are available in Supplementary Table 3.

### Enzyme-linked immunosorbent assay

The levels of secreted BDNF in cell culture supernates were measured using enzyme-linked immunosorbent assay (ELISA). ELISA kit for human BDNF (Cat. DBD00) was sourced from R&D Systems, and the experiments were conducted according to the manufacturer’s instructions. Absorbance at 450 nm was detected using a Multi-Mode Microplate Reader (BioTEK, USA). BDNF levels were calculated using a standard curve, and the results were expressed in pg/ml.

### Lysyl oxidase activity assay

The serum LOX activity in KPC mice upon LOX blocking antibody treatment was tested to verify its effectiveness. The activity of LOX protein was measured using an LOX activity assay kit (Abcam, Cat. ab112139) according to the manufacturer’s instructions. Briefly, 50 µl of serum sample or 50 µl of assay buffer were dispensed in triplicate into a 96-well plate. Subsequently, 50 µL of reaction mix was added to each well. The plate was then incubated for 1 h at 37 °C in the absence of light. Following incubation, fluorescence was measured using a microplate reader at Ex/Em = 560/590 nm.

### Bioinformatics analysis

For analysis of stromal score and matrisome score, RNA-seq data of PAAD, STAD, and CRC (COAD + READ) was acquired from the TCGA database (https://portal.gdc.cancer.gov/). The stromal score was calculated based on the ESTIMATE method^71^. The matrisome score was estimated according to a 29-gene tumor matrisome signature (*COL11A1, COL10A1, COL6A6, SPP1, CTHRC1, TNNC1, ABI3BP, PCOLCE2, OGN, MMP12, MMP1, ADAMTS5, GREM1, SFTPC, SFTPA2, SFTPD, FCN3, S100A2, CXCL13, WIF1, CHRDL1, CXCL2, IL6, HHIP, S100A12, LPL, CPB2, MAMDC2,* and *CD36*)^72^. To identify TIN-related molecular events, we stratified TCGA samples into ND-high and ND-low groups. The differentially expressed genes were analyzed by Limma package in R. The DEGs were set as log_2_|Fold Change| >_J1 and adjusted *P*_J<_J0.05. The molecular functions of DEGs were annotated by KEGG. For single-cell RNA-seq (scRNAseq) analysis of expression of NFs/AGMs, a human pancreatic tumor dataset deposited to GSA under the accession number CRA001160^73^ and a PanIN-afflicted donor pancreas dataset^74^ deposited to GEO under the accession number GSE229413 were included. The metadata containing cell type annotation was also released in the above datasets. “DotPlot”function in Seurat (version 4.3.0) R package was used for visualizing the expression of NFs/AGMs among cell clusters.

### Statistical analysis

Statistical analyses of quantifications were performed with unpaired, two-tailed t test, one-way ANOVA with Tukey’s multiple comparison test, or Fisher’s exact test using GraphPad Prism V7 (GraphPad Softward, Inc., La Jolla, CA). Cumulative survival time was calculated by the Kaplan-Meier method and analyzed by the log-rank Mantel-Cox test. Correlation analysis was determined using Spearman’s test. Detailed information regarding the mice, tumor, and cell numbers per condition, as well as statistical specifics for each experiment, can be found in the figure legends. P < 0.05 was considered statistically significant.

## ACKNOWLEDGMENTS

The research was supported by grants from National Natural Science Foundation of China (82173153, 83272879 and 82230087), Shanghai Pilot Program for Basic Research-Shanghai Jiao Tong University (21TQ1400225), and Innovative research team of high-level local universities in Shanghai (SHSMU-ZDCX20210802).

## AUTHOR CONTRIBUTIONS

Conceptualization: S.H.J., G.G.X. and Z.G.Z; Methodology: S.H.J., S.Z., Z.W.C. and M.H.Y.; Formal Analysis: S.H.J., S.Z., H.L. and D.J.L.; Resources: D.J.L., Z.Z.Z., M.H.Y., Y.W.S. and S.H.J.; Investigation: all authors; Supervision: S.H.J. and Z.G.Z.; Funding Acquisition: S.H.J. and Z.G.Z.; Writing-Original Draft: S.H.J, S.Z. and Z.G.Z.; Writing-Review & Editing: all authors.

## INCLUSION AND DIVERSITY

We support inclusive, diverse, and equitable conduct of research.

