## Supplemental Figures and Tables for "Mechanical cues of extracellular matrix determine tumor innervation"

**Supplementary Materials**

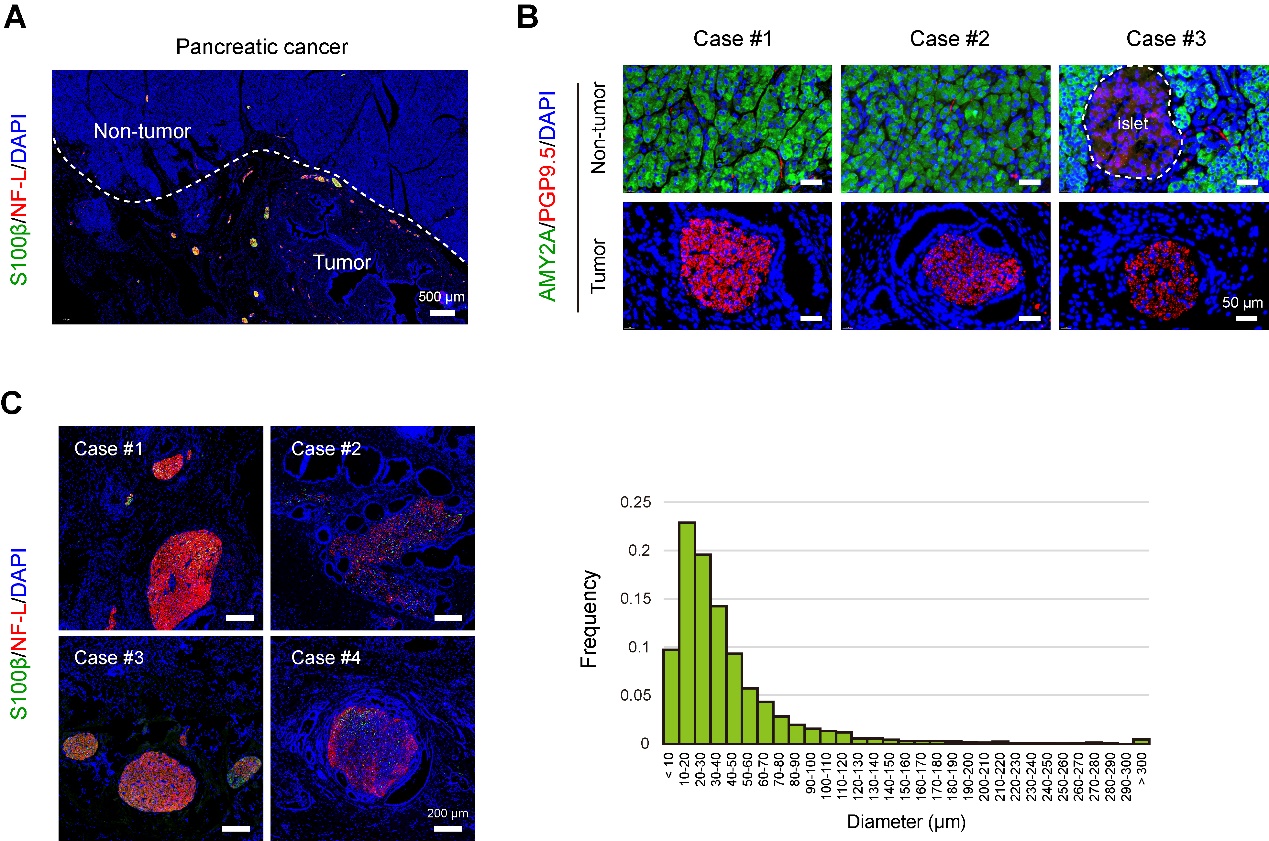

**Figure S1. Tumor innervation in PDAC**. (**A**) Representative histology images of tumor innervation in PDAC and the adjacent non-tumor tissues, obtained by co-immunofluorescence (IF) analysis of S100β and NF-L. Scale bar, 500 μm. (**B**) Representative images of PGP9.5^+^ nerves in PDAC and non-tumor pancreas tissues. Positive signal of PGP9.5 was also present in the islet, as marked by the dotted line. Acinar cells were marked by AMY2A. Scale bar, 50 μm. (**C**) Representative images of neural hypertrophy in PDAC, obtained by co-IF analysis of S100β and NF-L. Scale bar, 200 μm. The frequency of nerve diameter was shown on the right panel.

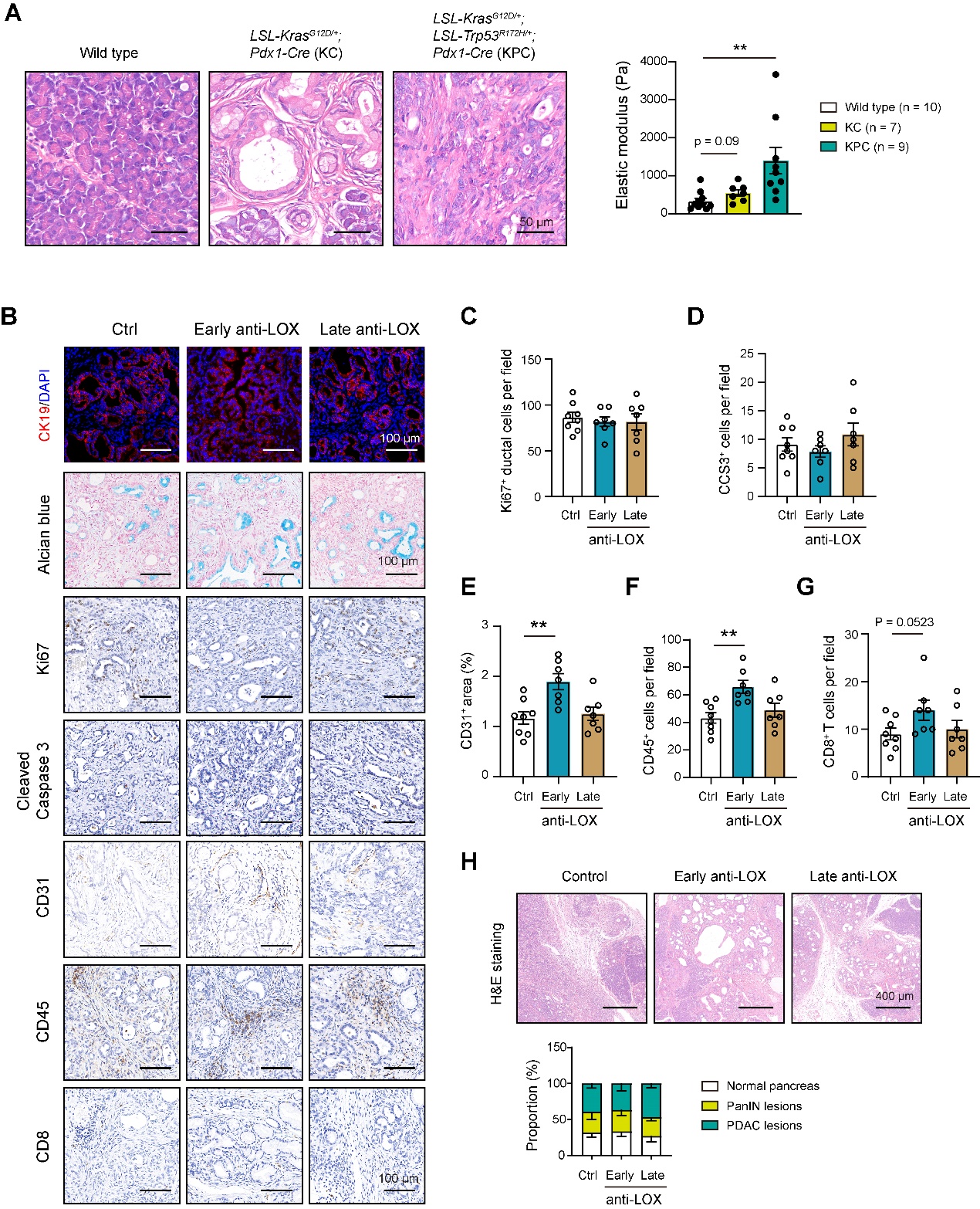

**Figure S2. The effects of anti-LOX treatment on the development of tumors bearing from KPC mice.** (**A**) Representative histology images of pancreas tissues from wide type (n = 10), KC (n = 7), and KPC mice (n = 9). The Young’s modulus of fresh pancreas samples was shown on the right panel. Scale bar, 50 μm. (**B**) Representative histology images, obtained by CK19 IF stain, alcian blue stain, Ki67 IHC stain, cleaved caspase 3 (CCS3) IHC stain, CD31 IHC stain, CD45 IHC stain, and CD8 IHC stain. Scale bar, 100 μm. (**C**) Quantification of Ki67^+^ ductal cancer cells. (**D**) Quantification of CCS3^+^ ductal cancer cells. (**E**) Quantification of CD31^+^ area. (**F**) Quantification of CD45^+^ cells. (**G**) Quantification of CD8^+^ T cells. (**H**) Representative H&E images of pancreatic sections. The bar graph shows the quantification of PanIN lesions and PDAC areas; scale bar, 400 μm. **P* < 0.05, ***P* < 0.01 and ****P* < 0.001. In **B**-**H**, n = 8 for control, n = 7 for early anti-LOX, and n = 7 for late anti-LOX. Values were compared by the one-way ANOVA multiple comparisons with Tukey’s method among groups (**A**, **C**-**G**) and the chi-square test (**H**).

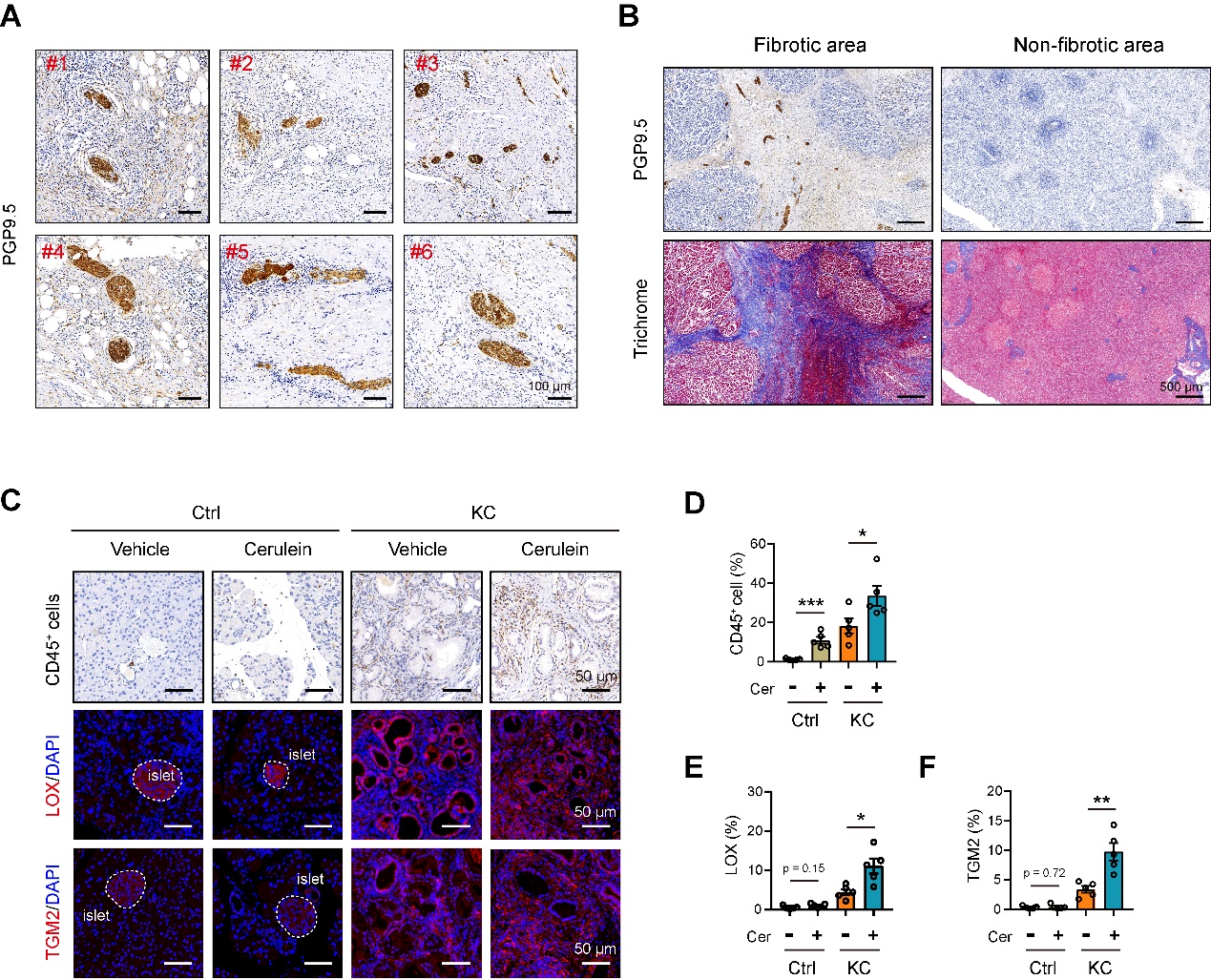

**Figure S5. Pancreatic innervation and expression of crosslinking enzymes in chronic pancreatitis.** (**A**) Representative histology images of nerves in human chronic pancreatitis (CP), obtained by PGP9.5 IHC stain. Scale bar, 100 μm. (**B**) Representative images of PGP9.5^+^ nerves in human CP tissues. The fibrotic and non-fibrotic areas were reflected by Trichrome stain. Scale bar, 500 μm. (**C**) Representative histology images, obtained by CD45 IHC stain, LOX IF stain, and TGM2 IF stain. Scale bar, 50 μm. Positive signals in LOX and TGM2 were marked by a dotted line. (**D**) Quantification of CD45^+^ cells. (**E**) Quantification of LOX^+^ areas. (**F**) Quantification of TGM2^+^ areas. In **C**-**F**, n = 5 per group. **P* < 0.05, ***P* < 0.01 and ****P* < 0.001. Values as mean ± SD and compared by the one-way ANOVA multiple comparisons with Tukey’s method among groups.

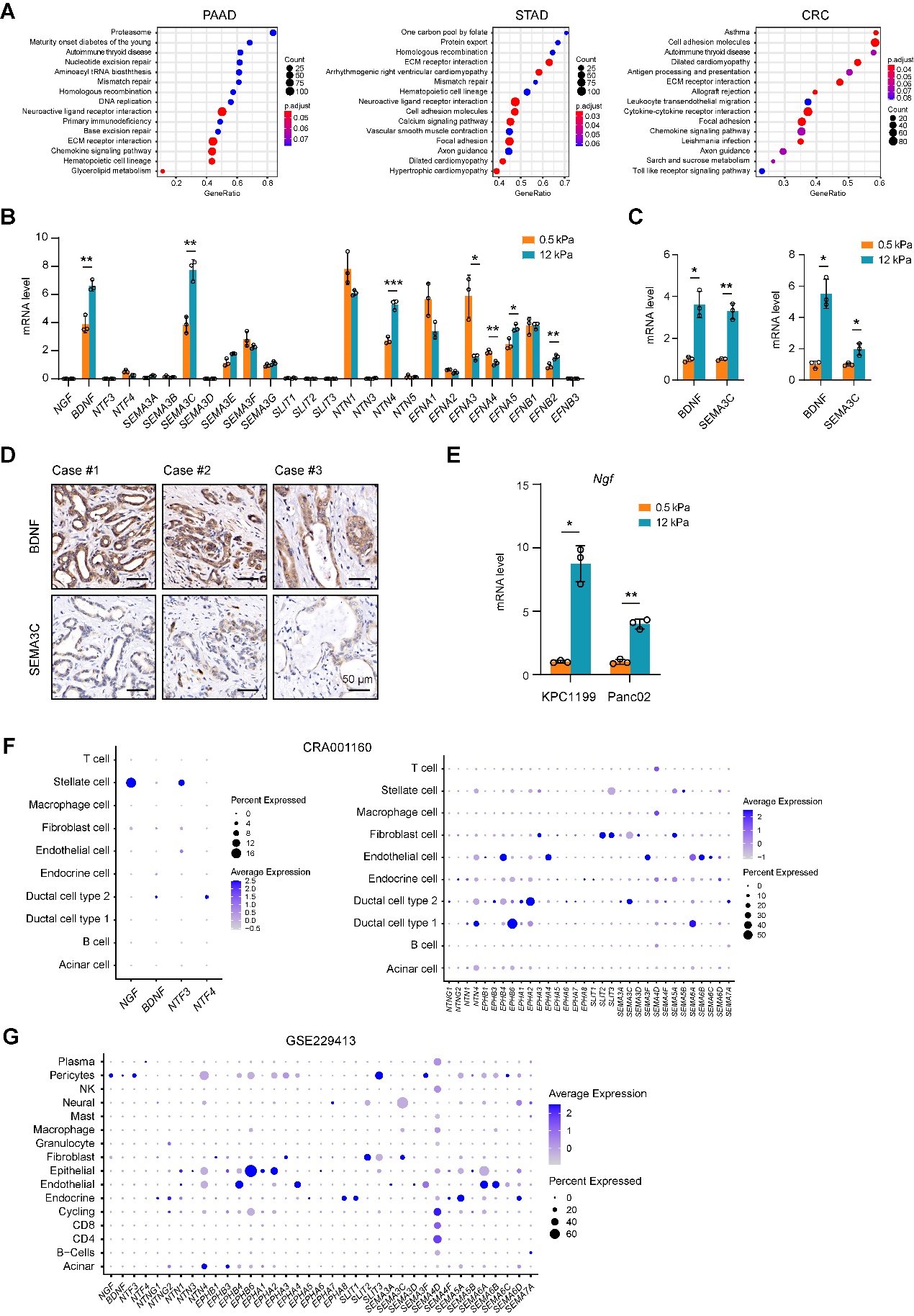

**Figure S4. The neurotrophic factors in PDAC and their links with ECM stiffness.** (**A**) KEGG annotation of the TIN-associated DEGs. Data from the TCGA cohort. (**B**) mRNA level of neurotrophins and axon guidance molecules in PDC0034 cells, cultured under soft and stiff matrices (n = 3). (**C**) mRNA level of *BDNF* and *SEMA3C* in AsPC-1 and Capan-2 cells, cultured under soft and stiff matrices (n = 3). (**D**) Representative IHC stains of BDNF and SEMA3C in PDAC tumor samples. Scale bar, 50 μm. (**E**) mRNA level of *Ngf* in mouse KPC1199 and Panc02 cells, cultured under soft and stiff matrices (n = 3). (**F**, **G**) Expression profile of neurotrophins and axon guidance molecules among various cell clusters from the single-cell RNA-seq datasets of human PDAC (**F**, CRA001160) and PanIN-afflicted donor pancreas (**G**, GSE229413). **P* < 0.05, ***P* < 0.01, ****P* < 0.001. Values were compared by the Student’s t test. Experiments were independently repeated three (**B**, **C**, **E**) times with similar results.

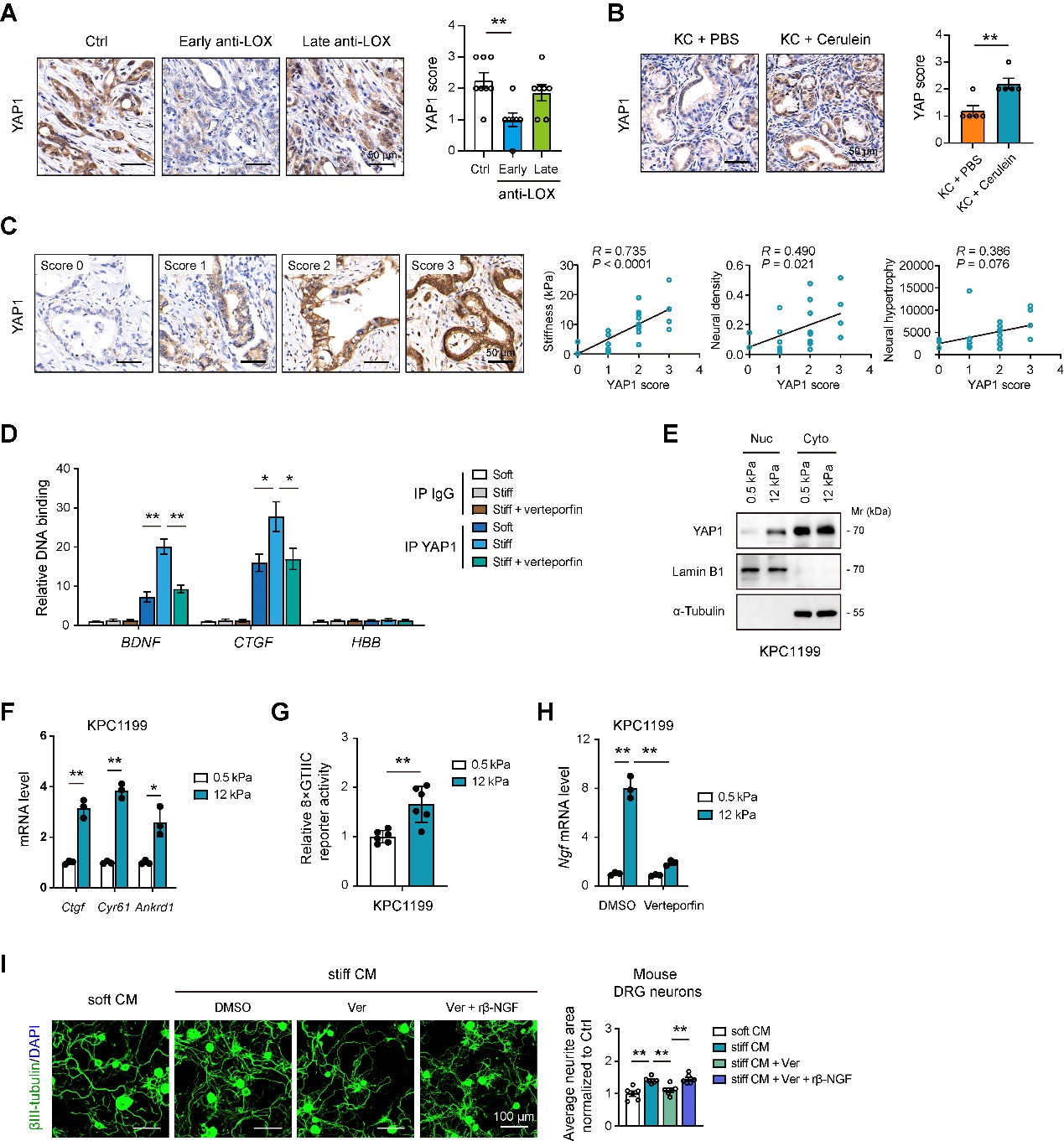

**Figure S5. YAP1-mediated neurotrophic effects in PDAC.** (**A**) YAP1 expression in tumor samples from control (n = 8), early anti-LOX (n = 7), and late anti-LOX (n = 7) groups. Scale bar, 50 μm. (**B**) YAP1 expression in pancreas samples from KC + PBS and KC + cerulein groups (n = 5 per group). Scale bar, 50 μm. (**C**) Correlation analysis of YAP1 expression with tissue stiffness, ND, and NH in a PDAC cohort (n = 22). Scale bar, 50 μm. (**D**) Chromatin immunoprecipitation of PDC0034 cells with control IgG or anti-YAP antibodies and subjected to detect the TEAD-binding regions present in the *BDNF* and *CTGF* promoters. Hemoglobin beta (HBB) was amplified as a negative control. (**E**) Western blot of YAP1 in the nuclear and cytoplasmic lysates in KPC1199 cells, cultured under soft and stiff matrices. Lamin B1 and α-Tubulin were loaded as the nuclear and cytoplasmic control, respectively. (**F**) mRNA level of *Ctgf*, *Cyr61*, and *Ankrd1* in KPC1199 cells, cultured under soft and stiff matrices (n = 3). (**G**) 8×GTIIC reporter activity in KPC1199 cells, cultured under soft and stiff matrices (n = 6). (**H**) *Ngf* mRNA level in KPC1199 cells, which cultured under soft and stiff matrices upon verteporfin treatment (n = 3). (**I**) Neurite outgrowth of mouse DRG neurons upon treatment with rBDNF and indicated CM from PDC0034 cells, which were cultured under stiff conditions and treated with or without verteporfin (n = 6). Scale bar, 100 μm. **P* < 0.05, ***P* < 0.01, ****P* < 0.001. Values were compared by the one-way ANOVA multiple comparisons with Tukey’s method among groups (**A**, **D**, **H**, **I**), Spearman's rank correlation methods (**C**), and Student’s t test (**B**, **F**, **G**). Experiments were independently repeated two (**D**, **I**) or three (**F**-**H**) times with similar results.

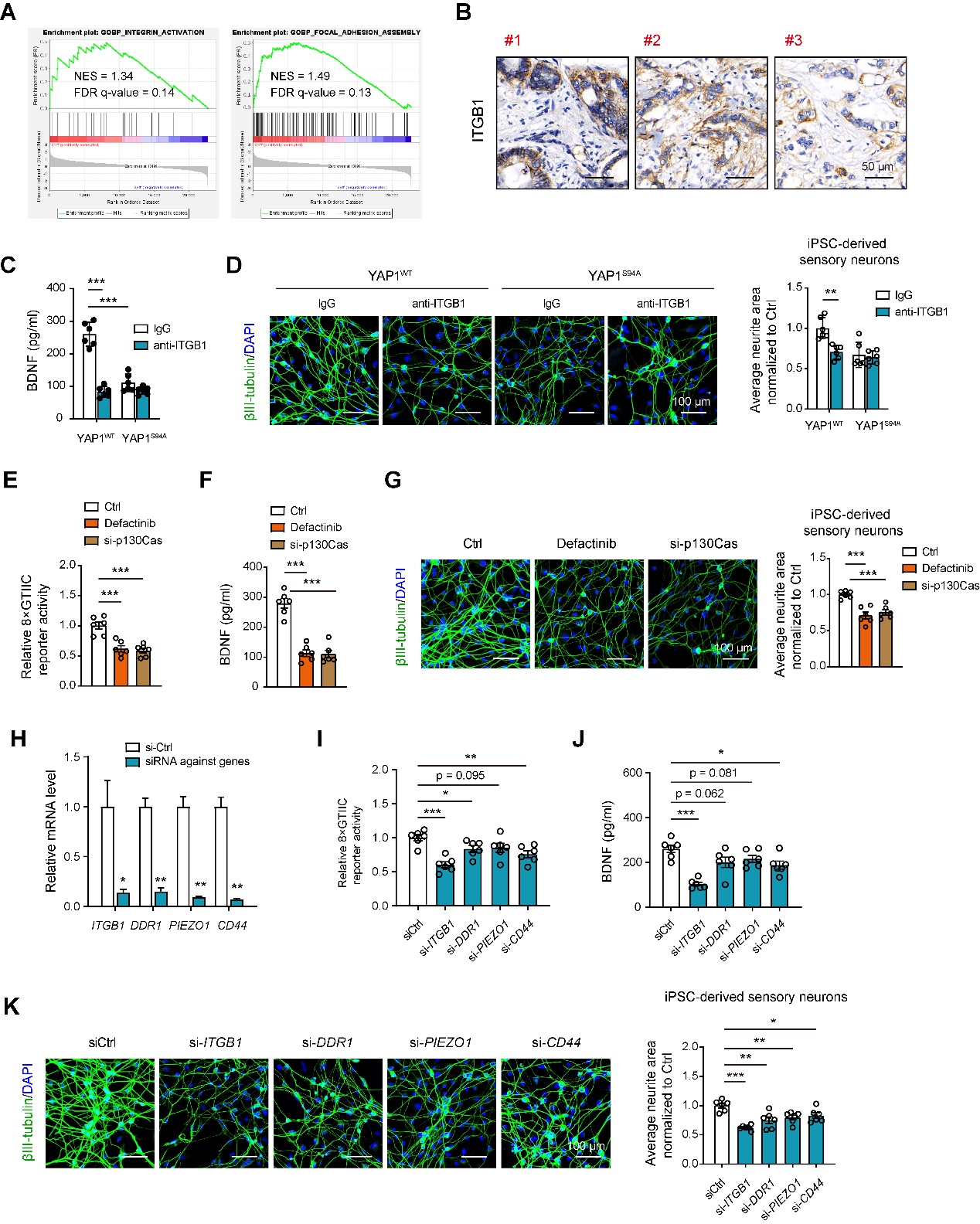

**Figure S6. ITGB1-containing integrins respond to matrix stiffness and contribute to YAP1-mediated neurotrophic effects.** (**A**) GSEA plot of integrin activation and focal adhesion assembly pathway among PDAC samples with high and low stiffness. (**B**) Representative IHC stains of ITGB1 in PDAC tumor samples. Scale bar, 50 μm. (**C**) BDNF secretion by YAP1^WT^ and YAP1^S94A^ PDC0034 cells, cultured under stiff matrices, in the presence or absence of anti-ITGB1 treatment (n = 6). (**D**) Neurite outgrowth of human iPSC-derived sensory neurons upon treatment with CM from YAP1^WT^ and YAP1^S94A^ PDC0034 cells, which were cultured under stiff conditions and treated with or without anti-ITGB1. Scale bar, 100 μm. (**E**) 8×GTIIC reporter activity in Ctrl, defactinib-treated, and si-p130Cas PDC0034 cells, cultured under stiff matrices (n = 6). (**F**) BDNF secretion by Ctrl, defactinib-treated, and si-p130Cas PDC0034 cells, cultured under stiff matrices (n = 6). (**G**) Neurite outgrowth of human iPSC-derived sensory neurons upon treatment with CM from Ctrl, defactinib-treated, and si-p130Cas PDC0034 cells, which were cultured under stiff conditions. Scale bar, 100 μm. (**H**) qPCR analysis of the knockdown efficiency of *ITGB1*, *DDR1*, *PIEZO1*, and *CD44* in PDC0034 cells (n = 3). (**I**) 8×GTIIC reporter activity in si-*ITGB1*, si-*DDR1*, si-*PIEZO1*, and si-*CD44* PDC0034 cells, cultured under stiff conditions (n = 6). (**J**) BDNF secretion by si-*ITGB1*, si-*DDR1*, si-*PIEZO1*, and si-*CD44* PDC0034 cells, cultured under stiff matrices (n = 6). (**K**) Neurite outgrowth of human iPSC-derived sensory neurons upon treatment with CM from si-*ITGB1*, si-*DDR1*, si-*PIEZO1*, and si-*CD44* PDC0034 cells, cultured under stiff conditions (n = 6). Scale bar, 50 μm. **P* < 0.05, ***P* < 0.01, ****P* < 0.001. Values were compared by the one-way ANOVA multiple comparisons with Tukey’s method among groups (**C**-**G**, **I**-**K**) and the Student’s t test (**H**). Experiments were independently repeated two (**D**, **G**, **K**) or three (**C**, **E**, **F**, **I**, **J**) times with similar results.

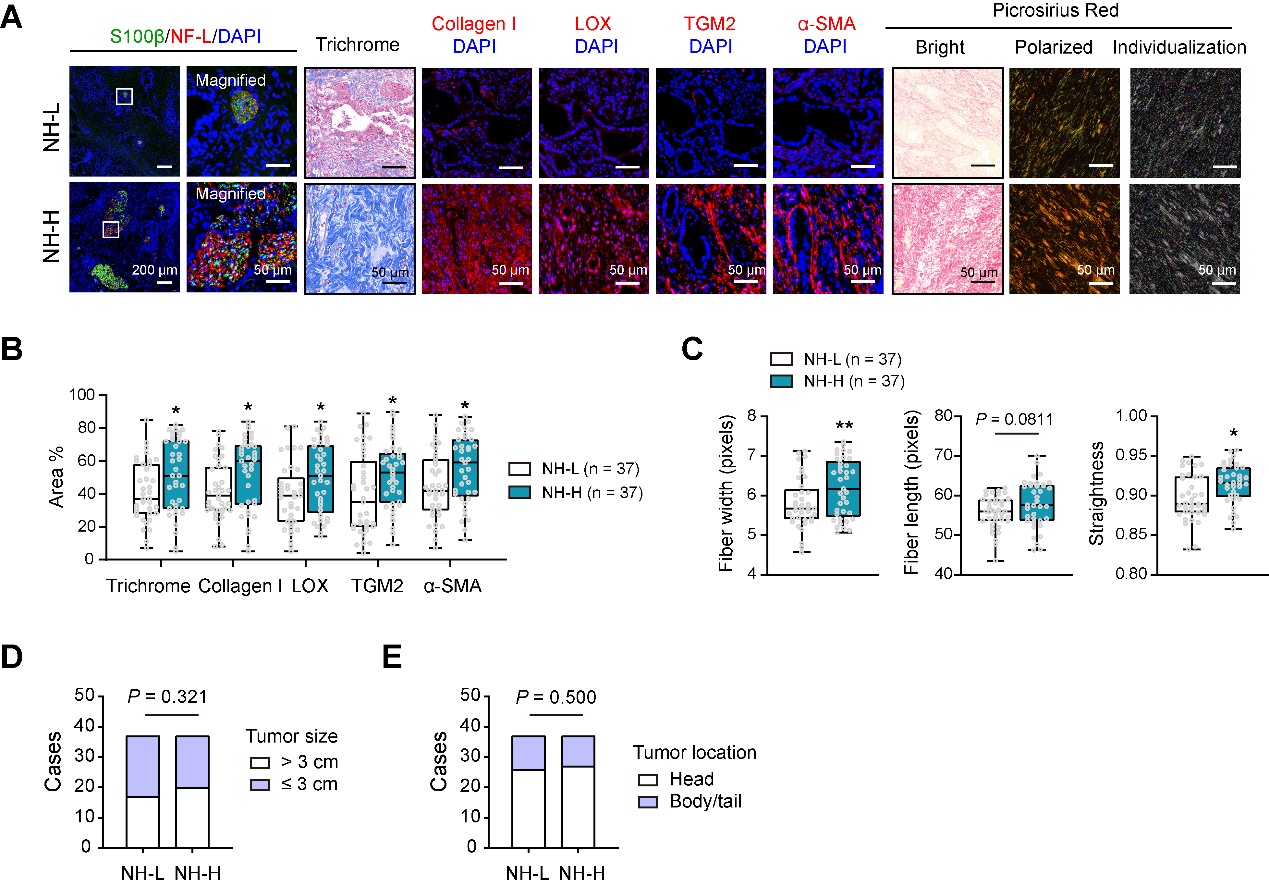

**Figure S7 ECM stiffness is linked to neural hypertrophy in PDAC**

(**A**) Representative histology images of NH-L and NH-H samples, obtained by co-immunofluorescence (IF) analysis of S100β and NF-L, Trichrome stain, collagen I IF stain, LOX IF stain, TGM2 IF stain, α-SMA IF stain, and picrosirius red (including polarized images and collagen fiber individualization analysis). Scale bar, 50 and 200 μm as indicated. (**B**) Quantification of Trichrome (collagen content), collagen I area, LOX area, TGM2 area, and α-SMA area between NH-L and NH-H samples (n = 37 per group). (**C**) Collagen fiber width, length, and straightness in NH-L and NH-H samples (n = 37 per group). (**D**) The association between NH and tumor size in PDAC (n = 37 per group). (**E**) The association between NH and tumor location in PDAC (n = 37 per group). **P* < 0.05 and ***P* < 0.01. Values were compared by the Student’s t test (**B**, **C**) and the chi-square test (**D**, **E**). NH, neural hypertrophy.

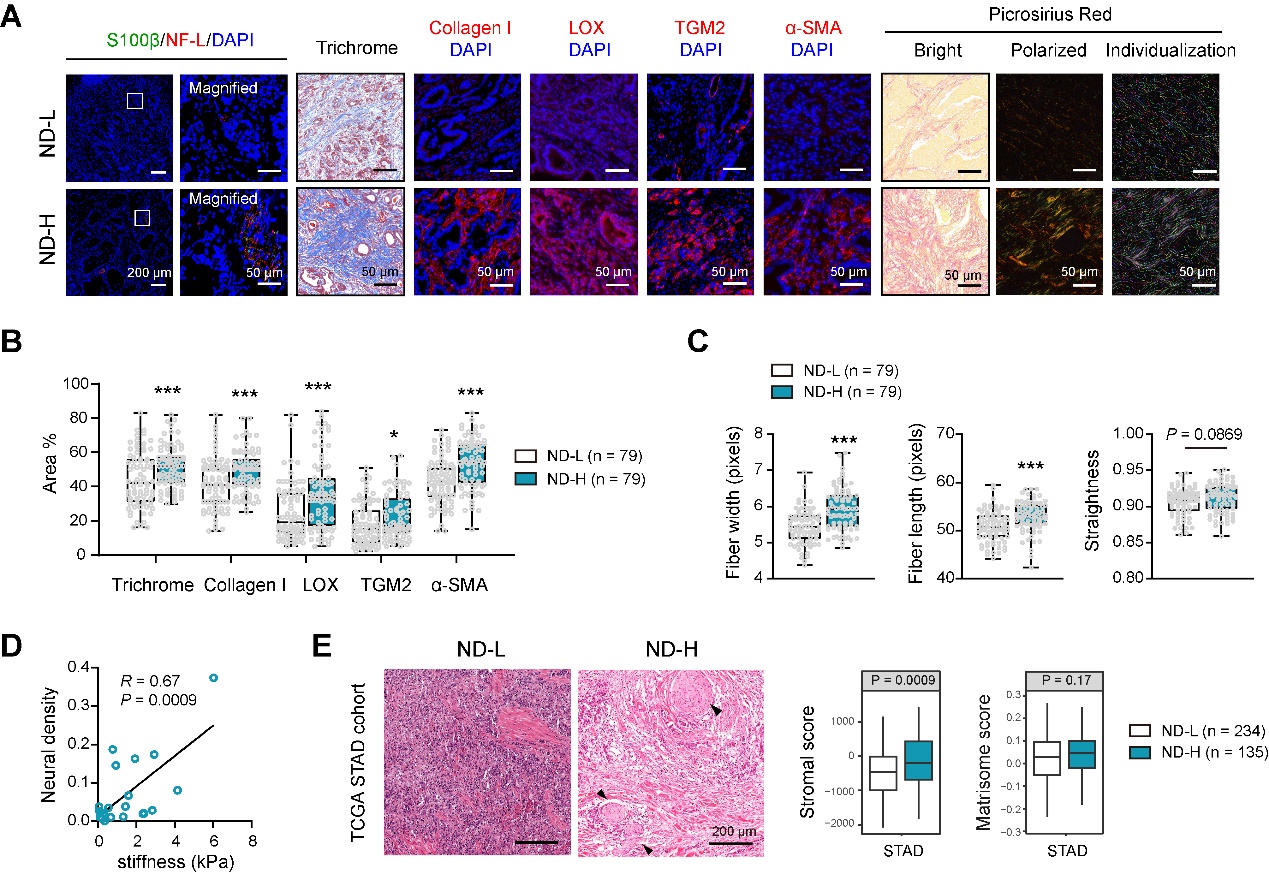

**Figure S8. ECM stiffness is linked to neural density in STAD.** (**A**) Representative histology images of ND-L and ND-H samples, obtained by co-IF analysis of S100β and NF-L, Trichrome stain, collagen I IF stain, LOX IF stain, TGM2 IF stain, α-SMA IF stain, and picrosirius red (including polarized images and collagen fiber individualization analysis) in STAD. Scale bar, 50 and 200 μm as indicated. (**B**) Quantification of Trichrome (collagen content), collagen I area, LOX area, TGM2 area, and α-SMA area between ND-L and ND-H samples (n = 79 per group). (**C**) Collagen fiber width, length, and straightness in ND-L and ND-H samples (n = 79 per group). (**D**) Correlation analysis of tissue stiffness with neural density in a STAD cohort (n = 21). (**E**) Stromal score and matrisome score of STAD cases, stratified by neural density. Data from the TCGA cohort (n = 369). The arrow heads indicate nerves. Scale bar, 200 μm. **P* < 0.05 and ****P* < 0.001. Values were compared by the Student’s t test (**B**, **C**, **E**) and the Spearman's rank correlation methods (**D**).

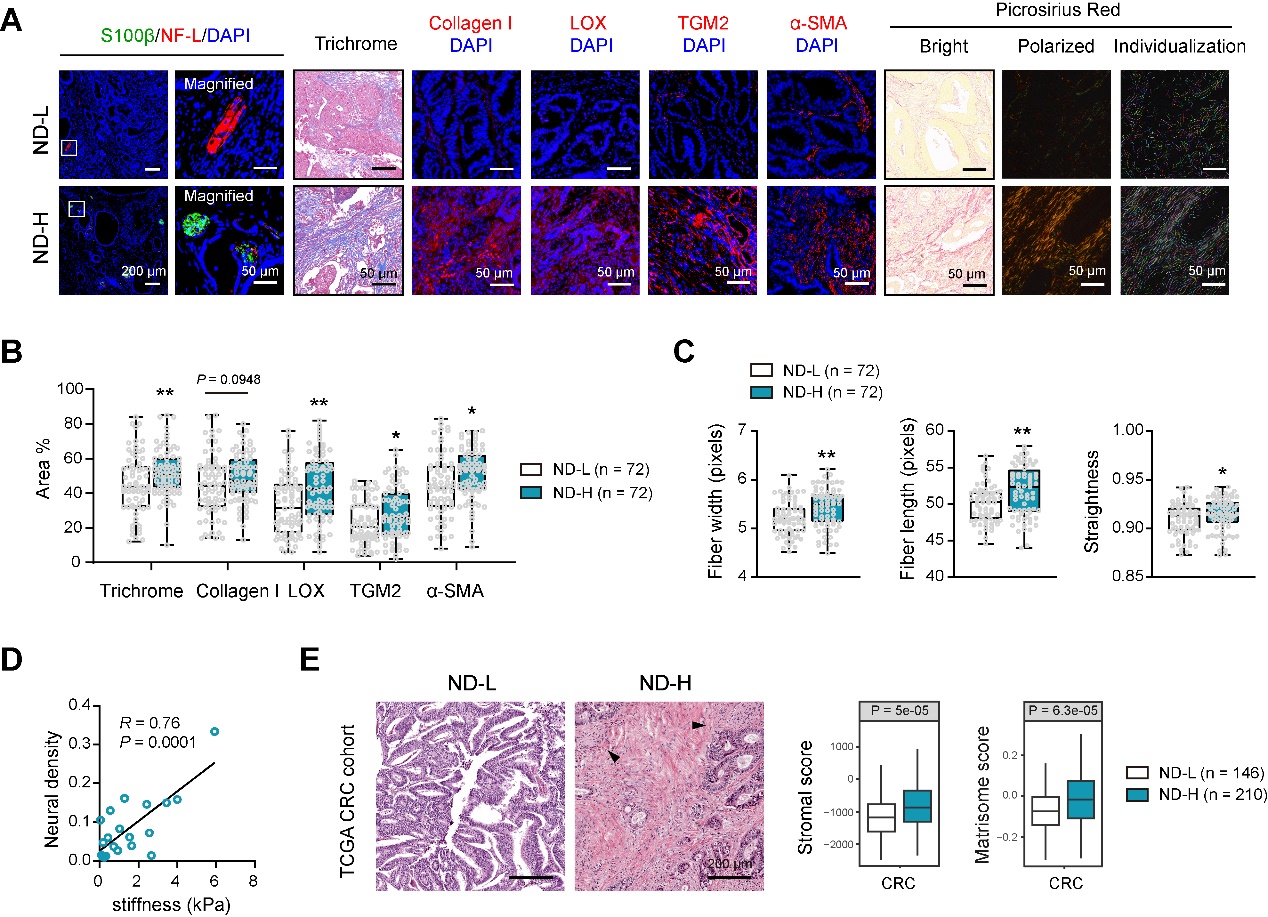

**Figure S9. ECM stiffness is linked to neural density in CRC.** (**A**) Representative histology images of ND-L and ND-H samples, obtained by co-IF analysis of S100β and NF-L, Trichrome stain, collagen I IF stain, LOX IF stain, TGM2 IF stain, α-SMA IF stain, and picrosirius red (including polarized images and collagen fiber individualization analysis) in CRC. Scale bar, 50 and 200 μm as indicated. (**B**) Quantification of Trichrome (collagen content), collagen I area, LOX area, TGM2 area, and α-SMA area between ND-L and ND-H samples (n = 72 per group). (**C**) Collagen fiber width, length, and straightness in ND-L and ND-H samples (n = 72 per group). (**D**) Correlation analysis of tissue stiffness with neural density in a CRC cohort (n = 20). (**E**) Stromal score and matrisome score of CRC cases, stratified by neural density. Data from the TCGA cohort (n = 375). The arrow heads indicate nerves. Scale bar, 200 μm. **P* < 0.05 and ****P* < 0.001. Values were compared by the Student’s t test (**B**, **C**, **E**) and the Spearman's rank correlation methods (**D**).

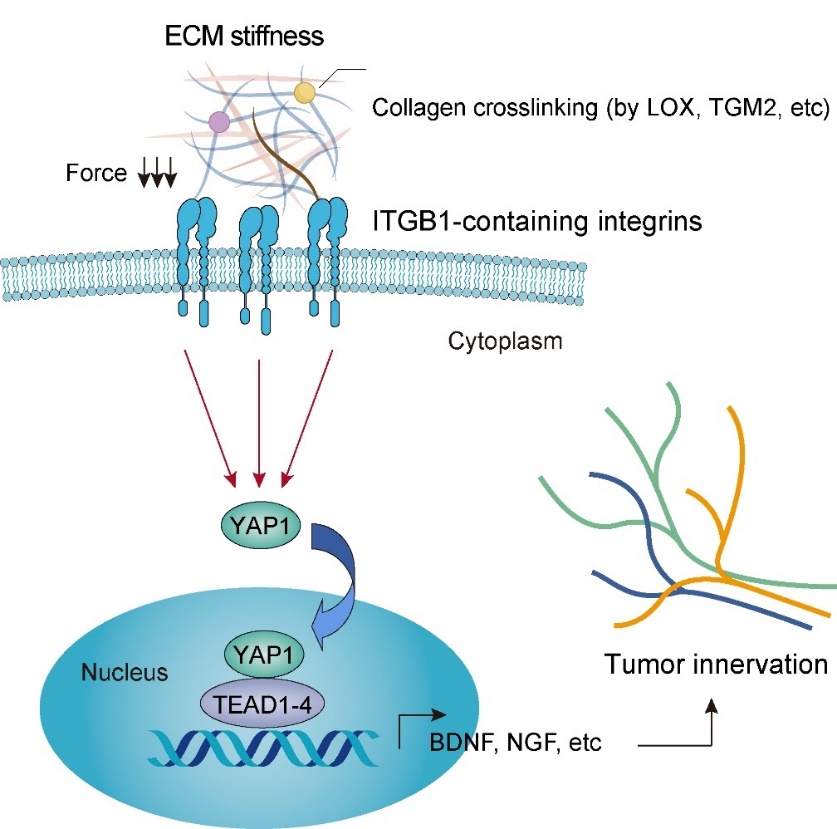

**Figure S10. Proposed mechanism model.** PDAC cells are capable of sensing the ECM stiffness through ITGB1-containing integrins, which activates YAP1-mediated mechanotransduction. YAP1 translocates to the nucleus and interacts with the TEAD transcription factor, leading to the expression of neurotrophic genes, particularly BDNF and NGF, and ultimately promoting the recruitment of nerves into the tumor microenvironment.

**Table S1. Stiffness-related differentially expressed proteins in the nuclear lysates of PDC0034 cells**

| **Gene Symbol** | **Protein name** | **P-value** | **Log_2_FC** |
| --- | --- | --- | --- |
| MPV17L2 | Mpv17-like protein 2 | 0.036247 | 6.036056 |
| ARHGEF10 | Rho guanine nucleotide exchange factor 10 | 0.002584 | 5.370695 |
| MIB1 | E3 ubiquitin-protein ligase MIB1 | 0.020161 | 5.069337 |
| CYB561A3 | Lysosomal membrane ascorbate-dependent ferrireductase CYB561A3 | 0.032468 | 4.669428 |
| RAD51AP1 | RAD51-associated protein 1 | 0.002424 | 4.354534 |
| MKI67 | KI67 Antigen (Fragment) | 0.001554 | 4.216501 |
| PDCL | Phosducin-like protein | 0.000747 | 3.431308 |
| ERCC6L | DNA excision repair protein ERCC-6-like | 0.00057 | 3.385062 |
| MCOLN2 | Mucolipin-2 | 0.002993 | 3.315692 |
| MPP7 | MAGUK p55 subfamily member 7 | 0.02789 | 3.156332 |
| HAUS6 | HAUS augmin-like complex subunit 6 | 0.019229 | 3.131708 |
| ZZZ3 | ZZ-type zinc finger-containing protein 3 | 0.003718 | 2.992061 |
| ASH1L | Histone-lysine N-methyltransferase ASH1L | 0.038287 | 2.991529 |
| KIF14 | Kinesin-like protein KIF14 | 0.00165 | 2.875351 |
| YAP1 | Transcriptional coactivator YAP1 | 0.003131 | 2.873115 |
| KIF20A | Kinesin-like protein KIF20A | 0.005202 | 2.848377 |
| TIMELESS | Protein timeless homolog | 0.006176 | 2.541121 |
| TIMM17A | Mitochondrial import inner membrane translocase subunit Tim17-A | 0.043519 | 2.511207 |
| CHUK | Inhibitor of nuclear factor kappa-B kinase subunit alpha | 0.024035 | 2.371879 |
| TYMS | Thymidylate synthase | 0.018867 | 2.280199 |
| SLC9A6 | Sodium/hydrogen exchanger 6 | 0.025707 | 2.275821 |
| PDE3B | cGMP-inhibited 3',5'-cyclic phosphodiesterase 3B | 0.000103 | 2.257079 |
| TGFBR1 | TGF-beta receptor type-1 | 0.037944 | 2.222823 |
| ZBTB44 | Zinc finger and BTB domain-containing protein 44 | 0.021436 | 2.213526 |
| CKS2 | Cyclin-dependent kinases regulatory subunit 2 | 0.020182 | 2.203427 |
| UHRF1 | E3 ubiquitin-protein ligase UHRF1 | 0.005272 | 2.183199 |
| BRAT1 | BRCA1-associated ATM activator 1 | 0.009005 | 2.165719 |
| ND5 | NADH-ubiquinone oxidoreductase chain 5 (Fragment) | 0.021504 | 2.138639 |
| RACGAP1 | Rac GTPase-activating protein 1 | 0.001895 | 2.134087 |
| RNF138 | E3 ubiquitin-protein ligase RNF138 | 0.027867 | 2.09423 |
| NCS1 | Neuronal calcium sensor 1 | 0.022442 | 2.068508 |
| PSMG4 | Proteasome assembly chaperone 4 | 0.015716 | 2.053139 |
| NUSAP1 | Nucleolar and spindle-associated protein 1 | 0.002105 | 2.033826 |
| HELLS | Lymphoid-specific helicase | 0.001815 | 1.933465 |
| TMEM129 | E3 ubiquitin-protein ligase TM129 | 0.034468 | 1.922337 |
| ZDHHC12 | Palmitoyltransferase ZDHHC12 | 0.029931 | 1.905848 |
| POLE2 | DNA polymerase epsilon subunit 2 | 0.031489 | 1.84953 |
| KDM5B | Lysine-specific demethylase 5B | 0.027921 | 1.791508 |
| CDCA7L | Cell division cycle-associated 7-like protein | 0.001776 | 1.791007 |
| USP20 | Ubiquitin carboxyl-terminal hydrolase 20 | 0.014999 | 1.77907 |
| ZNF687 | Zinc finger protein 687 | 0.004527 | 1.777953 |
| CKS1B | Cyclin-dependent kinases regulatory subunit 1 | 0.024142 | 1.773031 |
| CD320 | CD320 antigen | 0.005724 | 1.677451 |
| ND5 | NADH-ubiquinone oxidoreductase chain 5 | 0.001965 | 1.66899 |
| KPNA2 | Importin subunit alpha-1 | 0.005045 | 1.66075 |
| PRC1 | Protein regulator of cytokinesis 1 | 0.029725 | 1.600234 |
| ZNHIT3 | Zinc finger HIT domain-containing protein 3 | 0.036053 | 1.595453 |
| KIF23 | Kinesin-like protein KIF23 | 0.009321 | 1.593763 |
| CHST14 | Carbohydrate sulfotransferase 14 | 0.017411 | 1.579955 |
| FARP2 | FERM, ARHGEF and pleckstrin domain-containing protein 2 | 0.030092 | 1.564608 |
| KIF22 | Kinesin-like protein KIF22 | 0.033702 | 1.537087 |
| AURKA | Aurora kinase A | 0.032034 | 1.5353 |
| HIC2 | Hypermethylated in cancer 2 protein | 0.007438 | 1.516302 |
| RAB30 | Ras-related protein Rab-30 | 0.026485 | 1.484318 |
| TOP2A | DNA topoisomerase 2-alpha | 0.001816 | 1.452792 |
| DHX37 | Probable ATP-dependent RNA helicase DHX37 | 0.014227 | 1.444543 |
| SRCAP | Helicase SRCAP | 0.005291 | 1.435864 |
| ACTBL2 | Beta-actin-like protein 2 | 0.03652 | 1.433339 |
| LPCAT2 | Lysophosphatidylcholine acyltransferase 2 | 0.017197 | 1.43229 |
|  | cDNA FLJ56662, highly similar to Homo sapiens GTP binding protein 6 (GTPBP6), mRNA | 0.024266 | 1.42579 |
| SPAG5 | Sperm-associated antigen 5 | 0.039147 | 1.411092 |
| FBXO28 | F-box only protein 28 | 0.014724 | 1.40735 |
| DGCR8 | Microprocessor complex subunit DGCR8 | 0.013298 | 1.401729 |
| DEGS1 | Sphingolipid delta(4)-desaturase DES1 | 0.00735 | 1.385626 |
| CCNB1 | G2/mitotic-specific cyclin-B1 | 0.002044 | 1.382123 |
| KRT16 | Keratin, type I cytoskeletal 16 | 0.000499 | 1.377909 |
| SLC6A6 | Sodium- and chloride-dependent taurine transporter | 0.001679 | 1.362726 |
| KIFC1 | Kinesin-like protein KIFC1 | 0.010569 | 1.356785 |
| CENPF | Centromere protein F | 0.042481 | 1.350777 |
| MKI67 | Proliferation marker protein Ki-67 | 0.036042 | 1.348814 |
| ADAM15 | Disintegrin and metalloproteinase domain-containing protein 15 | 0.035572 | 1.335857 |
| KIF20B | Kinesin-like protein KIF20B | 0.032853 | 1.298935 |
| GRSF1 | G-rich sequence factor 1 | 0.001785 | 1.270124 |
| GTPBP8 | GTP-binding protein 8 | 0.029703 | 1.264211 |
| FDFT1 | Squalene synthase | 0.03852 | 1.260001 |
| RAB27A | Ras-related protein Rab-27A | 0.015136 | 1.259238 |
| SHCBP1 | SHC SH2 domain-binding protein 1 | 0.045441 | 1.258927 |
| AXL | Tyrosine-protein kinase receptor UFO | 0.001256 | 1.256509 |
| KRT17 | Keratin, type I cytoskeletal 17 | 0.008035 | 1.256452 |
|  | BPTI/Kunitz inhibitor domain-containing protein (Fragment) | 0.008155 | 1.255595 |
| NCAPD3 | Condensin-2 complex subunit D3 | 0.001189 | 1.255585 |
| SLC5A3 | Sodium/myo-inositol cotransporter | 0.030557 | 1.253007 |
| FANCI | Fanconi anemia group I protein | 0.025173 | 1.245231 |
| KRT6A | Keratin, type II cytoskeletal 6A | 0.003475 | 1.245166 |
| ALAS1 | 5-aminolevulinate synthase, non-specific, mitochondrial | 0.038182 | 1.24149 |
| TIMMDC1 | Complex I assembly factor TIMMDC1, mitochondrial | 0.005621 | 1.225912 |
| SFXN5 | Sideroflexin-5 | 0.012918 | 1.222495 |
| KRT6B | Keratin, type II cytoskeletal 6B | 0.005565 | 1.209716 |
| TBL1XR1 | TBL1X/Y related 1 (Fragment) | 0.044266 | 1.190109 |
| BCL2L12 | BCL2 like 12 | 0.044068 | 1.188948 |
| MGAT1 | Alpha-1,3-mannosyl-glycoprotein 2-beta-N-acetylglucosaminyltransferase | 0.040435 | 1.188402 |
| LDLR | Low-density lipoprotein receptor | 0.011287 | 1.178683 |
| UGT8 | 2-hydroxyacylsphingosine 1-beta-galactosyltransferase | 0.021446 | 1.178644 |
| SERPINA3 | Actin-like protein (Fragment) | 0.01683 | 1.178619 |
| CAV1 | Caveolin-1 | 0.008447 | 1.175029 |
| CIT | Citron Rho-interacting kinase | 0.006757 | 1.172902 |
| CHST7 | Carbohydrate sulfotransferase 7 | 0.046344 | 1.166905 |
| FLG2 | Filaggrin-2 | 0.040823 | 1.163333 |
| ANLN | Anillin | 0.030405 | 1.159938 |
| TPX2 | Targeting protein for Xklp2 | 0.031788 | 1.157994 |
| AMFR | E3 ubiquitin-protein ligase AMFR | 0.016732 | 1.133915 |
| TFB2M | Dimethyladenosine transferase 2, mitochondrial | 0.041748 | 1.131782 |
| NOA1 | Nitric oxide-associated protein 1 | 0.002279 | 1.128254 |
| KIF2C | Kinesin-like protein KIF2C | 0.021863 | 1.127855 |
| TFB1M | Dimethyladenosine transferase 1, mitochondrial | 0.002202 | 1.124099 |
| SLC38A2 | Sodium-coupled neutral amino acid symporter 2 | 0.028984 | 1.120643 |
| PRPSAP2 | Phosphoribosyl pyrophosphate synthase-associated protein 2 | 0.01207 | 1.116425 |
| CLCC1 | Chloride channel CLIC-like protein 1 | 0.00139 | 1.110724 |
| SFT2D2 | Vesicle transport protein SFT2B | 0.038344 | 1.108667 |
| UBE2G2 | Ubiquitin-conjugating enzyme E2 G2 | 0.036369 | 1.103096 |
| ERAL1 | GTPase Era, mitochondrial | 0.008453 | 1.101807 |
| PRIM2 | DNA primase large subunit | 0.00882 | 1.087004 |
| KIF4A | Chromosome-associated kinesin KIF4A | 0.00344 | 1.086266 |
| LMF1 | Lipase maturation factor 1 | 0.00547 | 1.085269 |
| MYO19 | Unconventional myosin-XIX | 0.044508 | 1.085131 |
| TOMM40L | Mitochondrial import receptor subunit TOM40B | 0.045486 | 1.084422 |
| SLC2A1 | Solute carrier family 2, facilitated glucose transporter member 1 | 0.007935 | 1.083855 |
| RAB18 | Ras-related protein Rab-18 | 0.004164 | 1.083551 |
| PKMYT1 | Membrane-associated tyrosine- and threonine-specific cdc2-inhibitory kinase | 0.009735 | 1.082156 |
| POLRMT | DNA-directed RNA polymerase, mitochondrial | 0.021382 | 1.078882 |
| FZD2 | Frizzled-2 | 0.005629 | 1.076449 |
| HSD17B11 | Estradiol 17-beta-dehydrogenase 11 | 0.023049 | 1.076001 |
| DNMT1 | DNA (cytosine-5)-methyltransferase 1 | 1.06E-05 | 1.075454 |
| DNAJA1 | DnaJ homolog subfamily A member 1 | 0.022991 | 1.070849 |
| CD44 | CD44 antigen | 0.014693 | 1.069911 |
| ALCAM | CD166 antigen | 0.019212 | 1.069038 |
| LPCAT1 | Lysophosphatidylcholine acyltransferase 1 | 0.00641 | 1.061162 |
| REEP4 | Receptor expression-enhancing protein 4 | 0.011949 | 1.048583 |
| TFRC | Transferrin receptor protein 1 | 0.001187 | 1.04658 |
| WWOX | WW domain-containing oxidoreductase | 0.012588 | 1.043162 |
| ARMC1 | Armadillo repeat-containing protein 1 | 0.005338 | 1.041794 |
| FAR1 | Fatty acyl-CoA reductase 1 | 0.019527 | 1.038554 |
| ADAM10 | Disintegrin and metalloproteinase domain-containing protein 10 | 0.010026 | 1.022875 |
| KRT14 | Keratin, type I cytoskeletal 14 | 0.000752 | 1.016317 |
| PEX5 | Peroxisomal targeting signal 1 receptor | 0.012232 | 1.01365 |
| TMEM33 | Transmembrane protein 33 | 0.003668 | 1.013614 |
| MRM3 | rRNA methyltransferase 3, mitochondrial | 0.002134 | 1.012393 |
| OAT | Ornithine aminotransferase, mitochondrial | 0.00211 | 1.007949 |
| RCC1L | RCC1-like G exchanging factor-like protein | 0.039837 | 1.006722 |
|  | Glutamyl-tRNA(Gln) amidotransferase subunit C, mitochondrial (Fragment) | 0.046582 | 1.002536 |
| MTG1 | Mitochondrial ribosome-associated GTPase 1 | 0.016038 | 0.991138 |
| ISG20L2 | Interferon-stimulated 20 kDa exonuclease-like 2 | 0.010854 | 0.990804 |
| ATR | Serine/threonine-protein kinase ATR | 0.017749 | 0.989668 |
| PIGK | GPI-anchor transamidase | 0.019539 | 0.989659 |
| RAB3B | Ras-related protein Rab-3B | 0.035615 | 0.987857 |
| CPNE8 | Copine-8 | 0.047139 | 0.986497 |
| CDKAL1 | Threonylcarbamoyladenosine tRNA methylthiotransferase | 0.006933 | 0.977652 |
| OS9 | Protein OS-9 | 0.001742 | 0.975612 |
| TF | Serotransferrin | 0.001141 | 0.968695 |
| AUP1 | Lipid droplet-regulating VLDL assembly factor AUP1 | 0.007903 | 0.966771 |
| NOC4L | Nucleolar complex protein 4 homolog | 0.015737 | 0.966528 |
| MAN2A1 | Alpha-mannosidase 2 | 0.016831 | 0.963912 |
| NDUFAF1 | Complex I intermediate-associated protein 30, mitochondrial | 0.013449 | 0.961834 |
| NDUFA9 | NADH dehydrogenase [ubiquinone] 1 alpha subcomplex subunit 9, mitochondrial | 0.011035 | 0.961769 |
| DHX30 | ATP-dependent RNA helicase DHX30 | 0.002742 | 0.958625 |
| SCAMP2 | Secretory carrier-associated membrane protein (Fragment) | 0.019559 | 0.954045 |
| RETSAT | All-trans-retinol 13,14-reductase | 0.005075 | 0.949389 |
| CLPTM1L | Lipid scramblase CLPTM1L | 0.011959 | 0.947476 |
| GPATCH8 | G patch domain-containing protein 8 | 0.019972 | 0.944925 |
| BPNT2 | Golgi-resident adenosine 3',5'-bisphosphate 3'-phosphatase | 0.037433 | 0.943182 |
| MRPS25 | 28S ribosomal protein S25, mitochondrial | 0.010115 | 0.940462 |
| TMX2 | Thioredoxin-related transmembrane protein 2 | 0.000106 | 0.939814 |
| TCAF1 | TRPM8 channel-associated factor 1 | 0.036753 | 0.937764 |
| OCIAD2 | OCIA domain-containing protein 2 | 0.005974 | 0.933939 |
| UTP20 | Small subunit processome component 20 homolog | 0.004695 | 0.931554 |
| TXNDC15 | Thioredoxin domain-containing protein 15 | 0.009914 | 0.926836 |
| PEF1 | Peflin | 0.028492 | 0.92611 |
| DYSF | Dysferlin | 0.023485 | 0.92191 |
| MRPL32 | 39S ribosomal protein L32, mitochondrial | 0.04003 | 0.920674 |
| CYB5B | Cytochrome b5 type B | 0.044569 | 0.917913 |
| TECR | Very-long-chain enoyl-CoA reductase | 0.009579 | 0.914871 |
| CCDC86 | Coiled-coil domain-containing protein 86 | 0.022723 | 0.913413 |
| YME1L1 | ATP-dependent zinc metalloprotease YME1L1 | 0.004312 | 0.912829 |
| ABCC5 | ATP-binding cassette sub-family C member 5 | 0.034466 | 0.909406 |
| LIMA1 | LIM domain and actin-binding protein 1 | 0.009733 | 0.908556 |
| GLS | Glutaminase kidney isoform, mitochondrial | 0.00106 | 0.907766 |
| ITCH | E3 ubiquitin-protein ligase Itchy homolog | 0.047591 | 0.906647 |
| ARL6IP1 | ADP-ribosylation factor-like protein 6-interacting protein 1 | 0.007296 | 0.906566 |
| ELAC2 | Zinc phosphodiesterase ELAC protein 2 | 0.003579 | 0.906097 |
| MTX2 | Metaxin-2 (Fragment) | 0.045075 | 0.905315 |
| EGFR | Epidermal growth factor receptor | 0.015404 | 0.901477 |
| LRRC8C | Volume-regulated anion channel subunit LRRC8C | 0.036856 | 0.896875 |
| DDX49 | Probable ATP-dependent RNA helicase DDX49 | 0.017838 | 0.89679 |
| GOLIM4 | Golgi integral membrane protein 4 | 0.004582 | 0.896461 |
| SPOUT1 | Putative methyltransferase C9orf114 | 0.044412 | 0.893989 |
| SSR4 | Translocon-associated protein subunit delta | 0.026884 | 0.892566 |
| SMC6 | Structural maintenance of chromosomes 6 | 0.022103 | 0.892133 |
| NECTIN2 | Nectin-2 | 0.032758 | 0.890367 |
| WDHD1 | WD repeat and HMG-box DNA-binding protein 1 | 0.000655 | 0.885639 |
| BAG6 | Large proline-rich protein BAG6 | 0.027374 | 0.884731 |
| MPZL1 | Myelin protein zero-like protein 1 | 0.028125 | 0.882201 |
| C2CD2 | C2 domain-containing protein 2 | 0.045683 | 0.880549 |
| FBXO17 | F-box only protein 17 | 0.010334 | 0.877301 |
| MTERF3 | Transcription termination factor 3, mitochondrial | 0.020458 | 0.874141 |
| DDX28 | Probable ATP-dependent RNA helicase DDX28 | 0.023559 | 0.872828 |
| ALG2 | Alpha-1,3/1,6-mannosyltransferase ALG2 | 0.021237 | 0.872658 |
| SEC11A | Signal peptidase complex catalytic subunit SEC11A | 0.039887 | 0.870155 |
| RAB2A | Ras-related protein Rab-2A | 0.006257 | 0.869043 |
| SCFD2 | Sec1 family domain-containing protein 2 | 0.002838 | 0.863636 |
| ITGB5 | Integrin beta-5 | 0.00988 | 0.863266 |
| MRPS23 | 28S ribosomal protein S23, mitochondrial | 0.00601 | 0.862738 |
| NOP53 | Ribosome biogenesis protein NOP53 | 0.033513 | 0.86021 |
| NDUFB5 | NADH dehydrogenase [ubiquinone] 1 beta subcomplex subunit 5, mitochondrial | 0.048619 | 0.85986 |
| MGAT2 | Alpha-1,6-mannosyl-glycoprotein 2-beta-N-acetylglucosaminyltransferase | 0.029248 | 0.856124 |
| MCM7 | DNA replication licensing factor MCM7 | 0.035786 | 0.855166 |
| EMC1 | ER membrane protein complex subunit 1 | 0.033392 | 0.854554 |
| CYC1 | Cytochrome c1, heme protein, mitochondrial | 0.010693 | 0.849094 |
| SNX14 | Sorting nexin-14 | 0.007608 | 0.847422 |
| PREPL | Prolyl endopeptidase-like | 0.000701 | 0.8461 |
| MAN1A2 | Mannosyl-oligosaccharide 1,2-alpha-mannosidase IB | 0.020827 | 0.845844 |
| APOB | Apolipoprotein B-100 | 0.039165 | 0.84525 |
| DNAJA3 | DnaJ homolog subfamily A member 3, mitochondrial | 0.016605 | 0.845101 |
| MET | Hepatocyte growth factor receptor | 0.021063 | 0.844275 |
| ILVBL | 2-hydroxyacyl-CoA lyase 2 | 0.044188 | 0.844056 |
| MARCHF5 | E3 ubiquitin-protein ligase MARCHF5 | 0.032839 | 0.843414 |
| CWF19L2 | CWF19-like protein 2 | 0.009076 | 0.843209 |
| GHDC | GH3 domain-containing protein | 0.046493 | 0.83834 |
| TBRG4 | FAST kinase domain-containing protein 4 | 0.00422 | 0.836894 |
| MTFP1 | Mitochondrial fission process protein 1 | 0.03447 | 0.83677 |
| DDX54 | ATP-dependent RNA helicase DDX54 | 0.043668 | 0.833714 |
| POLDIP2 | Polymerase delta-interacting protein 2 | 0.002895 | 0.833523 |
| CYP51A1 | Lanosterol 14-alpha demethylase | 0.003694 | 0.831103 |
| MRPL3 | 39S ribosomal protein L3, mitochondrial | 0.036137 | 0.830329 |
| CBX3 | Chromobox protein homolog 3 | 0.008338 | 0.829788 |
| HACD3 | Very-long-chain (3R)-3-hydroxyacyl-CoA dehydratase 3 | 0.030507 | 0.826063 |
| SLC25A5 | ADP/ATP translocase 2 | 0.025671 | 0.825312 |
| HLTF | Helicase-like transcription factor | 0.02046 | 0.824365 |
| CAVIN1 | Caveolae-associated protein 1 | 0.018358 | 0.823867 |
| DERL1 | Derlin-1 | 0.005452 | 0.823758 |
| NOP9 | Nucleolar protein 9 | 0.015975 | 0.823519 |
| TMBIM6 | Bax inhibitor 1 | 0.00401 | 0.820084 |
| GPAA1 | Glycosylphosphatidylinositol anchor attachment 1 protein | 0.029907 | 0.819419 |
| DBN1 | Drebrin | 0.002571 | 0.818976 |
| OXA1L | Oxidase (Cytochrome c) assembly 1-like | 0.011118 | 0.81845 |
| QSOX2 | Sulfhydryl oxidase 2 | 0.009933 | 0.818209 |
| POLA2 | DNA polymerase alpha subunit B | 0.014853 | 0.818071 |
| RHOA | Transforming protein RhoA | 0.011367 | 0.817735 |
| DNAJC11 | DnaJ homolog subfamily C member 11 | 0.036758 | 0.817679 |
| PLD6 | Mitochondrial cardiolipin hydrolase | 0.016651 | 0.817156 |
| DDOST | Dolichyl-diphosphooligosaccharide--protein glycosyltransferase 48 kDa subunit | 0.043421 | 0.812974 |
| NDUFV3 | NADH dehydrogenase [ubiquinone] flavoprotein 3, mitochondrial | 0.00377 | 0.811991 |
| SEC61A1 | Protein transport protein Sec61 subunit alpha isoform 1 | 0.00205 | 0.810558 |
| DDX47 | Probable ATP-dependent RNA helicase DDX47 | 0.032925 | 0.810544 |
| PNPLA6 | Patatin-like phospholipase domain-containing protein 6 | 0.017505 | 0.804575 |
| DNAJC1 | DnaJ homolog subfamily C member 1 | 0.037999 | 0.804382 |
| AQR | RNA helicase aquarius | 0.045276 | 0.802211 |
| ABCC4 | ATP-binding cassette sub-family C member 4 | 0.018155 | 0.801468 |
| ALG9 | Alpha-1,2-mannosyltransferase ALG9 | 0.009583 | 0.800077 |
| CD151 | CD151 antigen | 0.021787 | 0.799966 |
| SLC25A32 | Mitochondrial folate transporter/carrier | 0.032281 | 0.797247 |
| LCLAT1 | Lysocardiolipin acyltransferase 1 | 0.010861 | 0.797139 |
| PPP2R1B | Serine/threonine-protein phosphatase 2A 65 kDa regulatory subunit A beta isoform | 0.000219 | 0.796527 |
| MRPL21 | 39S ribosomal protein L21, mitochondrial | 0.033121 | 0.79601 |
| IPO8 | Importin-8 | 0.006068 | 0.795319 |
| PTDSS1 | Phosphatidylserine synthase 1 | 0.014577 | 0.795178 |
| FASTKD3 | FAST kinase domain-containing protein 3, mitochondrial | 0.001646 | 0.795171 |
| GALNT2 | Polypeptide N-acetylgalactosaminyltransferase 2 | 0.016866 | 0.790477 |
| MRPL28 | 39S ribosomal protein L28, mitochondrial | 0.020036 | 0.790435 |
| ADGRG6 | Adhesion G-protein coupled receptor G6 | 0.018051 | 0.787015 |
| ECSIT | Evolutionarily conserved signaling intermediate in Toll pathway, mitochondrial | 0.011264 | 0.786574 |
| NCBP2AS2 | Protein NCBP2AS2 | 0.025689 | 0.786257 |
|  | cDNA FLJ39696 fis, clone SMINT2011033, highly similar to Sorting and assembly machinery component 50 homolog | 0.023783 | 0.785667 |
| RAP1B | Ras-related protein Rap-1b | 0.025522 | 0.779125 |
| RAB21 | Ras-related protein Rab-21 | 0.00923 | 0.778962 |
| DNAJB12 | DnaJ homolog subfamily B member 12 | 0.026475 | 0.773147 |
| SLC25A4 | ADP/ATP translocase 1 | 0.017998 | 0.770589 |
| MRPL39 | 39S ribosomal protein L39, mitochondrial | 0.003977 | 0.768206 |
| SUPT16H | FACT complex subunit SPT16 | 0.00611 | 0.767653 |
| COMTD1 | Catechol O-methyltransferase domain-containing protein 1 | 0.014883 | 0.765136 |
| PRIM1 | DNA primase small subunit | 0.003037 | 0.764768 |
| MRPL58 | Peptidyl-tRNA hydrolase ICT1, mitochondrial | 0.008982 | 0.760854 |
| TMEM199 | Transmembrane protein 199 | 3.61E-05 | 0.760769 |
| DDX18 | ATP-dependent RNA helicase DDX18 | 0.018586 | 0.760163 |
| SPTLC2 | Serine palmitoyltransferase 2 | 0.030206 | 0.759341 |
| SGPP1 | Sphingosine-1-phosphate phosphatase 1 | 0.007854 | 0.759216 |
| PET100 | Protein PET100 homolog, mitochondrial | 0.001089 | 0.759096 |
| HEATR1 | HEAT repeat-containing protein 1 | 0.04069 | 0.757457 |
| ITGA6 | Integrin alpha-6 | 0.013913 | 0.75684 |
| SSRP1 | FACT complex subunit SSRP1 | 0.006857 | 0.755535 |
| MRPS34 | 28S ribosomal protein S34, mitochondrial | 0.005662 | 0.755134 |
| NSDHL | Sterol-4-alpha-carboxylate 3-dehydrogenase, decarboxylating | 0.009797 | 0.754939 |
| NDUFB10 | NADH dehydrogenase [ubiquinone] 1 beta subcomplex subunit 10 | 0.011275 | 0.754141 |
| RAB32 | Ras-related protein Rab-32 | 0.02377 | 0.752412 |
| MRPL19 | 39S ribosomal protein L19, mitochondrial | 0.007927 | 0.752378 |
| RAB12 | Ras-related protein Rab-12 | 0.004722 | 0.752325 |
| ANO6 | Anoctamin-6 | 0.010134 | 0.751299 |
| MRPL35 | 39S ribosomal protein L35, mitochondrial | 0.036926 | 0.750746 |
| NOP16 | Nucleolar protein 16 | 0.036039 | 0.748259 |
| NT5DC2 | 5'-nucleotidase domain-containing protein 2 | 0.004288 | 0.746796 |
| NDUFS2 | NADH dehydrogenase [ubiquinone] iron-sulfur protein 2, mitochondrial | 0.041492 | 0.746033 |
| GOLT1B | Vesicle transport protein GOT1B | 0.003535 | 0.744905 |
| NDUFS4 | NADH dehydrogenase [ubiquinone] iron-sulfur protein 4, mitochondrial | 0.004417 | 0.744583 |
| DIMT1 | Probable dimethyladenosine transferase | 0.001356 | 0.744504 |
| STT3A | Dolichyl-diphosphooligosaccharide--protein glycosyltransferase subunit STT3A | 0.035589 | 0.744442 |
| COQ10B | Coenzyme Q-binding protein COQ10 homolog B, mitochondrial | 0.045985 | 0.742134 |
| CTCF | Transcriptional repressor CTCF | 0.014419 | 0.738256 |
| EMC7 | ER membrane protein complex subunit 7 | 0.020246 | 0.736991 |
| MCU | Calcium uniporter protein, mitochondrial | 0.015604 | 0.736331 |
| FASTKD5 | FAST kinase domain-containing protein 5, mitochondrial | 0.027714 | 0.734175 |
| MYBBP1A | Myb-binding protein 1A | 0.017266 | 0.733329 |
| KPNA3 | Importin subunit alpha-4 | 0.003972 | 0.732231 |
| NOLC1 | Nucleolar and coiled-body phosphoprotein 1 (Fragment) | 0.017362 | 0.729758 |
| RAB5C | Ras-related protein Rab-5C | 0.018424 | 0.729128 |
| DIPK2A | Divergent protein kinase domain 2A | 0.016361 | 0.728466 |
| MTA1 | Metastasis-associated protein MTA1 | 0.019378 | 0.728324 |
| MRGBP | MRG/MORF4L-binding protein | 0.023856 | 0.727882 |
| CDC42 | Cell division control protein 42 homolog | 0.029441 | 0.727496 |
| TRRAP | Transformation/transcription domain associated protein | 0.035055 | 0.7268 |
| PTPN1 | Tyrosine-protein phosphatase non-receptor type 1 | 0.012609 | 0.724736 |
| ITPR2 | Inositol 1,4,5-trisphosphate receptor type 2 | 0.043215 | 0.724035 |
| DEK | Protein DEK | 0.019385 | 0.72384 |
| MCM6 | DNA replication licensing factor MCM6 | 0.004224 | 0.723737 |
| ZNF593 | Zinc finger protein 593 | 0.007541 | 0.723593 |
| TMEM70 | Transmembrane protein 70, mitochondrial | 0.039476 | 0.721989 |
| CPNE3 | Copine-3 | 0.003133 | 0.721311 |
| CD59 | CD59 glycoprotein | 0.043638 | 0.721074 |
| DAG1 | Dystroglycan 1 | 0.020523 | 0.720054 |
| POR | NADPH--cytochrome P450 reductase | 0.016545 | 0.718588 |
| UQCRQ | Cytochrome b-c1 complex subunit 8 | 0.033337 | 0.718242 |
| DNAJB6 | DnaJ homolog subfamily B member 6 | 0.03866 | 0.718218 |
| AGK | Acylglycerol kinase, mitochondrial | 0.010974 | 0.716883 |
| SLC4A2 | Anion exchange protein 2 | 0.000524 | 0.7164 |
| DHCR7 | 7-dehydrocholesterol reductase | 0.017413 | 0.716388 |
| POLD1 | DNA polymerase delta catalytic subunit | 0.033764 | 0.716363 |
| ATP13A3 | ATPase 13A3 | 0.036344 | 0.71547 |
| RANBP6 | Ran-binding protein 6 | 0.019917 | 0.714692 |
| SMC4 | Structural maintenance of chromosomes protein 4 | 0.005515 | 0.71391 |
| RETREG3 | Reticulophagy regulator 3 | 0.00816 | 0.713685 |
| THEM6 | Protein THEM6 | 0.029873 | 0.713651 |
| CDK1 | Cyclin-dependent kinase 1 | 0.010544 | 0.713105 |
| OMA1 | Metalloendopeptidase OMA1, mitochondrial | 0.00716 | 0.712099 |
| CYP20A1 | Cytochrome P450 20A1 | 0.018755 | 0.710335 |
| OSBPL8 | Oxysterol-binding protein-related protein 8 | 0.006257 | 0.710083 |
| CLPX | ATP-dependent Clp protease ATP-binding subunit clpX-like, mitochondrial | 0.004907 | 0.70998 |
| GEMIN4 | Gem-associated protein 4 | 0.005974 | 0.70899 |
| ATAD1 | Outer mitochondrial transmembrane helix translocase | 0.007032 | 0.708916 |
| DHCR24 | Delta(24)-sterol reductase | 0.039771 | 0.707625 |
| RP2 | Protein XRP2 | 0.02174 | 0.707102 |
| CBX5 | Chromobox protein homolog 5 | 0.029539 | 0.705766 |
| SLC35A4 | SLC35A4 upstream open reading frame protein | 0.040723 | 0.704703 |
| WDR46 | Chromosome 6 open reading frame 11 | 0.041685 | 0.701875 |
| MPHOSPH6 | M-phase phosphoprotein 6 | 0.005024 | 0.699846 |
| SYF2 | Pre-mRNA-splicing factor SYF2 | 0.003011 | 0.699033 |
| COX5A | Cytochrome c oxidase subunit 5A, mitochondrial | 0.006961 | 0.698435 |
| NDUFC2 | NADH dehydrogenase [ubiquinone] 1 subunit C2 | 0.010243 | 0.698274 |
| ABCD3 | ATP-binding cassette sub-family D member 3 | 0.030518 | 0.698267 |
| EXOSC4 | Exosome complex component RRP41 | 0.046525 | 0.696452 |
| NDUFS3 | NADH dehydrogenase [ubiquinone] iron-sulfur protein 3, mitochondrial | 0.004323 | 0.695409 |
| NDUFB6 | NADH dehydrogenase [ubiquinone] 1 beta subcomplex subunit 6 | 0.000804 | 0.694923 |
| SLC12A4 | Solute carrier family 12 member 4 | 0.033926 | 0.694698 |
| DCAF1 | DDB1- and CUL4-associated factor 1 | 0.024958 | 0.693938 |
| MRPL13 | 39S ribosomal protein L13, mitochondrial | 0.018725 | 0.693457 |
| ATL2 | Atlastin-2 | 0.000808 | 0.693443 |
| RAB13 | Ras-related protein Rab-13 | 0.020482 | 0.692064 |
| COX20 | Cytochrome c oxidase assembly protein COX20, mitochondrial | 0.018474 | 0.68962 |
| GPAT4 | Glycerol-3-phosphate acyltransferase 4 | 0.022362 | 0.688711 |
| NAT10 | RNA cytidine acetyltransferase | 0.008139 | 0.688571 |
| COMMD3-BMI1 | COMMD3-BMI1 readthrough | 0.039232 | 0.688469 |
| KIAA0100 | Bridge-like lipid transfer protein family member 2 | 0.026649 | 0.688453 |
| TAP2 | Antigen peptide transporter 2 | 0.045095 | 0.68837 |
| ITGAV | ITGAV protein | 0.011868 | 0.686594 |
| CERS6 | Ceramide synthase 6 | 0.005579 | 0.685109 |
| MLEC | Malectin | 0.049699 | 0.685021 |
| RAB11FIP1 | Rab11 family-interacting protein 1 | 0.027967 | 0.682386 |
| CARD19 | Chromosome 9 open reading frame 89, isoform CRA_b | 0.04312 | 0.682234 |
| STUB1 | E3 ubiquitin-protein ligase CHIP | 0.028562 | 0.681667 |
| PRPS1 | Ribose-phosphate pyrophosphokinase 1 | 0.015546 | 0.680081 |
| GNB1 | Guanine nucleotide-binding protein G(I)/G(S)/G(T) subunit beta-1 | 0.01874 | 0.679225 |
| DHRSX | Dehydrogenase/reductase SDR family member on chromosome X | 0.00358 | 0.677499 |
| ATAD3B | ATPase family AAA domain-containing protein 3B | 0.027957 | 0.676744 |
| PIGS | GPI transamidase component PIG-S | 0.002684 | 0.676335 |
| CPNE1 | Copine-1 | 0.042249 | 0.675881 |
|  | Annexin (Fragment) | 0.014839 | 0.674838 |
| CCDC137 | Coiled-coil domain-containing protein 137 | 0.020798 | 0.673008 |
| MRPL43 | 39S ribosomal protein L43, mitochondrial | 0.015952 | 0.672857 |
| WDR36 | WD repeat-containing protein 36 | 0.030047 | 0.672315 |
| NMD3 | 60S ribosomal export protein NMD3 | 0.030443 | 0.671443 |
| MSH6 | DNA mismatch repair protein Msh6 | 0.014346 | 0.670963 |
| PGAM5 | Serine/threonine-protein phosphatase PGAM5, mitochondrial | 0.019117 | 0.670908 |
| CDK5RAP1 | Mitochondrial tRNA methylthiotransferase CDK5RAP1 | 0.002236 | 0.670874 |
| GRPEL1 | GrpE protein homolog 1, mitochondrial | 0.004627 | 0.670604 |
| OCIAD1 | OCIA domain-containing protein 1 | 0.016909 | 0.666421 |
| SACM1L | Phosphatidylinositol-3-phosphatase SAC1 | 0.001805 | 0.665402 |
| ANAPC4 | Anaphase-promoting complex subunit 4 | 0.001553 | 0.665027 |
| LPCAT3 | Lysophospholipid acyltransferase 5 | 0.003941 | 0.664642 |
| POLA1 | DNA polymerase alpha catalytic subunit | 0.020817 | 0.663754 |
| ESYT1 | Extended synaptotagmin-1 | 0.009274 | 0.663367 |
| BAZ1A | Bromodomain adjacent to zinc finger domain protein 1A | 0.025769 | 0.662536 |
| PIGT | GPI transamidase component PIG-T | 0.022513 | 0.661262 |
| HCCS | Holocytochrome c-type synthase | 0.012173 | 0.66084 |
| INTS15 | Integrator complex subunit 15 | 0.034767 | 0.659558 |
| DDX21 | Nucleolar RNA helicase 2 | 0.01067 | 0.659016 |
| MRPL9 | 39S ribosomal protein L9, mitochondrial | 0.000987 | 0.658971 |
| SLC25A40 | Probable mitochondrial glutathione transporter SLC25A40 | 0.007205 | 0.657247 |
| CYB5R3 | NADH-cytochrome b5 reductase 3 | 0.017413 | 0.656584 |
| MRPS5 | 28S ribosomal protein S5, mitochondrial | 0.04949 | 0.656256 |
| MTPAP | Poly(A) RNA polymerase, mitochondrial | 0.035197 | 0.655991 |
| MCM3 | DNA replication licensing factor MCM3 | 0.007373 | 0.654897 |
| ZMPSTE24 | CAAX prenyl protease 1 homolog | 0.007718 | 0.654492 |
| RHOT2 | Mitochondrial Rho GTPase 2 | 0.004109 | 0.654321 |
| TM9SF3 | Transmembrane 9 superfamily member 3 | 0.026155 | 0.654012 |
| NDUFA7 | NADH dehydrogenase [ubiquinone] 1 alpha subcomplex subunit 7 | 0.046128 | 0.653844 |
| SLIRP | SRA stem-loop-interacting RNA-binding protein, mitochondrial | 0.006046 | 0.653809 |
| ATP1B3 | Sodium/potassium-transporting ATPase subunit beta-3 | 0.023658 | 0.653635 |
| MTARC2 | Mitochondrial amidoxime reducing component 2 | 0.02549 | 0.653055 |
| ITPRID2 | Protein ITPRID2 | 0.049477 | 0.650158 |
| NOMO2 | BOS complex subunit NOMO2 | 0.013326 | 0.649733 |
| MMUT | Methylmalonyl-CoA mutase, mitochondrial | 0.010818 | 0.648855 |
| SRPRA | Signal recognition particle receptor subunit alpha | 0.031815 | 0.648468 |
| MPDU1 | Mannose-P-dolichol utilization defect 1 protein | 0.034773 | 0.648047 |
| SLC25A19 | Mitochondrial thiamine pyrophosphate carrier | 0.008816 | 0.647655 |
| MTCH2 | Mitochondrial carrier homolog 2 | 0.012221 | 0.647384 |
| ATPAF2 | ATP synthase mitochondrial F1 complex assembly factor 2 | 0.022457 | 0.645827 |
| TIMM29 | Mitochondrial import inner membrane translocase subunit Tim29 | 0.034967 | 0.644001 |
| MCUB | Calcium uniporter regulatory subunit MCUb, mitochondrial | 0.033811 | 0.642432 |
| TM9SF2 | Transmembrane 9 superfamily member 2 | 0.025638 | 0.641863 |
| MRPL23 | 39S ribosomal protein L23, mitochondrial | 0.01183 | 0.640858 |
| ALDH18A1 | Delta-1-pyrroline-5-carboxylate synthase | 0.005513 | 0.640396 |
| ALG1 | Chitobiosyldiphosphodolichol beta-mannosyltransferase | 0.015429 | 0.640393 |
| SPTLC1 | Serine palmitoyltransferase 1 | 0.011551 | 0.639369 |
| TOM1L2 | TOM1-like protein 2 | 0.014163 | 0.638069 |
| HMOX2 | Heme oxygenase 2 | 0.018451 | 0.636436 |
| GLG1 | Golgi apparatus protein 1 | 0.003756 | 0.636288 |
| NDUFA8 | NADH dehydrogenase [ubiquinone] 1 alpha subcomplex subunit 8 | 0.009013 | 0.635867 |
| MYO6 | Unconventional myosin-VI | 0.017262 | 0.635284 |
| ACSL3 | Fatty acid CoA ligase Acsl3 | 0.012886 | 0.634787 |
| ITGA1 | Integrin alpha-1 | 0.019038 | 0.634253 |
| SLC25A6 | ADP/ATP translocase 3 | 0.001389 | 0.633342 |
| NT5E | 5'-nucleotidase | 0.012815 | 0.632623 |
| PLCD3 | 1-phosphatidylinositol 4,5-bisphosphate phosphodiesterase delta-3 | 0.017465 | 0.63242 |
| MCM4 | DNA replication licensing factor MCM4 | 0.02643 | 0.630588 |
| MTREX | Exosome RNA helicase MTR4 | 0.019221 | 0.630537 |
| WDR76 | WD repeat-containing protein 76 | 0.021315 | 0.63021 |
| ATL3 | Atlastin-3 | 0.025834 | 0.629988 |
| SLC25A12 | Electrogenic aspartate/glutamate antiporter SLC25A12, mitochondrial | 0.006344 | 0.629987 |
| CLPTM1 | Putative lipid scramblase CLPTM1 | 0.048402 | 0.627875 |
| EXD2 | Exonuclease 3'-5' domain-containing protein 2 | 0.001739 | 0.627592 |
| TAP1 | Antigen peptide transporter 1 | 0.028898 | 0.626853 |
| EMG1 | Ribosomal RNA small subunit methyltransferase NEP1 | 0.047167 | 0.62671 |
| CAMK2G | Calcium/calmodulin-dependent protein kinase type II subunit gamma | 0.01969 | 0.626087 |
| SEC61B | Protein transport protein Sec61 subunit beta | 0.007863 | 0.625276 |
| EPHA2 | Ephrin type-A receptor 2 | 0.02203 | 0.625106 |
| FKBP11 | Peptidyl-prolyl cis-trans isomerase FKBP11 | 0.018148 | 0.623958 |
| NCLN | BOS complex subunit NCLN | 0.015814 | 0.623943 |
| DDX56 | Probable ATP-dependent RNA helicase DDX56 | 0.017164 | 0.623311 |
| DPM1 | Dolichol-phosphate mannosyltransferase subunit 1 | 0.036352 | 0.622812 |
| GPX8 | Probable glutathione peroxidase 8 | 0.031489 | 0.621029 |
| ALG6 | Dolichyl pyrophosphate Man9GlcNAc2 alpha-1,3-glucosyltransferase | 0.006858 | 0.62078 |
| ZNF318 | Zinc finger protein 318 | 0.005088 | 0.620237 |
| RFC3 | Replication factor C subunit 3 | 0.001584 | 0.617624 |
| CTNNB1 | Catenin beta-1 | 0.016192 | 0.617365 |
| CLN6 | Ceroid-lipofuscinosis neuronal protein 6 | 0.018052 | 0.615631 |
| COX7A2 | Cytochrome c oxidase subunit 7A2, mitochondrial | 0.002847 | 0.615075 |
| FBXO2 | F-box only protein 2 | 0.041514 | 0.614639 |
| FAF2 | FAS-associated factor 2 | 0.017368 | 0.613461 |
| F11R | Junctional adhesion molecule A | 0.0467 | 0.613364 |
| RAB11A | Ras-related protein Rab-11A | 0.024156 | 0.611573 |
| DDX5 | Probable ATP-dependent RNA helicase DDX5 | 0.030612 | 0.609761 |
| CERS2 | Ceramide synthase 2 | 0.021872 | 0.60971 |
| SRPRB | Signal recognition particle receptor subunit beta | 0.038646 | 0.609453 |
| HK2 | Hexokinase-2 | 0.018289 | 0.609124 |
| MYOF | Myoferlin | 0.008746 | 0.608576 |
| RAB10 | Ras-related protein Rab-10 | 0.030012 | 0.60738 |
| ATP1A1 | Sodium/potassium-transporting ATPase subunit alpha-1 | 0.021342 | 0.607265 |
| NDUFA10 | NADH dehydrogenase [ubiquinone] 1 alpha subcomplex subunit 10, mitochondrial | 0.010102 | 0.606781 |
| CHMP4B | Charged multivesicular body protein 4b | 0.025393 | 0.606653 |
| RBBP7 | Histone-binding protein RBBP7 | 0.044594 | 0.606257 |
| LRPPRC | Leucine-rich PPR motif-containing protein, mitochondrial | 0.006755 | 0.606248 |
| NT5DC3 | 5'-nucleotidase domain-containing protein 3 | 0.002286 | 0.604295 |
| EXOSC5 | Exosome complex component RRP46 | 0.038376 | 0.604116 |
| C19orf25 | UPF0449 protein C19orf25 | 0.015487 | 0.60367 |
| PGRMC1 | Membrane-associated progesterone receptor component 1 | 0.012627 | 0.603147 |
| UQCRC1 | Cytochrome b-c1 complex subunit 1, mitochondrial | 0.029557 | 0.602001 |
| PES1 | Pescadillo homolog | 0.030072 | 0.601382 |
| UQCRB | Cytochrome b-c1 complex subunit 7 | 0.004359 | 0.600674 |
| SMARCA1 | Probable global transcription activator SNF2L1 | 0.024733 | 0.599637 |
| COX5B | Cytochrome c oxidase subunit 5B, mitochondrial (Fragment) | 0.045058 | 0.597371 |
| TMX3 | Protein disulfide-isomerase TMX3 | 0.019757 | 0.597338 |
| KDM2A | Lysine-specific demethylase 2A | 0.049455 | 0.596699 |
| NDUFA5 | NADH dehydrogenase [ubiquinone] 1 alpha subcomplex subunit 5 | 0.006932 | 0.596383 |
| CDK13 | Cyclin-dependent kinase 13 | 0.014333 | 0.594293 |
| GNA13 | Guanine nucleotide-binding protein subunit alpha-13 | 0.034653 | 0.594165 |
| ATP6AP2 | Renin receptor | 0.047248 | 0.593983 |
| MGME1 | Mitochondrial genome maintenance exonuclease 1 | 0.034416 | 0.593095 |
| CUL2 | Cullin-2 | 0.019773 | 0.590026 |
| BSG | Basigin | 0.025926 | 0.589619 |
| PDE12 | 2',5'-phosphodiesterase 12 | 0.036107 | 0.589584 |
| TMX1 | Thioredoxin-related transmembrane protein 1 | 0.014463 | 0.589573 |
| FZD6 | Frizzled-6 | 0.032144 | 0.58945 |
| SFXN1 | Sideroflexin-1 | 0.020149 | 0.588729 |
| GFM1 | Elongation factor G, mitochondrial | 0.000958 | 0.585886 |
| SUPT6H | Transcription elongation factor SPT6 | 0.049069 | 0.585853 |
| GALNS | N-acetylgalactosamine-6-sulfatase | 0.009153 | -0.58813 |
| PEBP1 | Phosphatidylethanolamine-binding protein 1 | 0.012003 | -0.58875 |
| TFPI | Tissue factor pathway inhibitor | 0.009414 | -0.60529 |
| SNX9 | Sorting nexin-9 | 0.029104 | -0.60601 |
| NUP54 | Nucleoporin p54 | 0.016815 | -0.60976 |
| TP53I3 | Quinone oxidoreductase PIG3 | 0.004208 | -0.60978 |
| SEPTIN2 | Septin-2 | 0.006048 | -0.61196 |
| TRIM25 | E3 ubiquitin/ISG15 ligase TRIM25 | 0.02543 | -0.61546 |
| CTSC | Dipeptidyl peptidase 1 | 0.035738 | -0.62968 |
| HMCES | Abasic site processing protein HMCES | 0.038933 | -0.63526 |
| SEPTIN7 | Septin-7 | 0.02378 | -0.63606 |
| THYN1 | Thymocyte nuclear protein 1 | 0.017285 | -0.63743 |
| AKR1A1 | Aldo-keto reductase family 1 member A1 | 0.011179 | -0.64321 |
| CRTAP | Cartilage-associated protein | 0.028705 | -0.64426 |
| CSTF2T | Cleavage stimulation factor subunit 2 tau variant | 0.030675 | -0.66727 |
| TPI1 | Triosephosphate isomerase | 0.018819 | -0.67118 |
| API5 | cDNA FLJ52148, highly similar to Apoptosis inhibitor 5 | 0.029329 | -0.67388 |
| WARS1 | Tryptophan--tRNA ligase, cytoplasmic | 0.038731 | -0.67636 |
| MSN | Moesin | 0.02749 | -0.67882 |
| HINT1 | Adenosine 5'-monophosphoramidase HINT1 | 0.039075 | -0.68069 |
| RAB33B | Ras-related protein Rab-33B | 0.032305 | -0.68152 |
| PYGB | Glycogen phosphorylase, brain form | 0.008137 | -0.68612 |
| EEF1B2 | Elongation factor 1-beta | 0.004893 | -0.69107 |
| PKN1 | Serine/threonine-protein kinase N1 | 0.026914 | -0.69439 |
| LAMB2 | Laminin subunit beta-2 | 0.040615 | -0.69531 |
| UROD | Uroporphyrinogen decarboxylase | 0.033201 | -0.70392 |
| CPOX | Oxygen-dependent coproporphyrinogen-III oxidase, mitochondrial | 0.049152 | -0.70459 |
| HMGB2 | High mobility group protein B2 | 0.040553 | -0.71643 |
| WDR11 | WD repeat-containing protein 11 | 0.020325 | -0.7166 |
| SERPINB6 | Serpin B6 | 0.000327 | -0.72015 |
| KRT18 | Keratin, type I cytoskeletal 18 | 0.040394 | -0.72399 |
| TKT | Transketolase | 0.044357 | -0.72451 |
| GAA | Lysosomal alpha-glucosidase | 0.018769 | -0.72462 |
| CTSA | Lysosomal protective protein | 0.008753 | -0.72464 |
| S100A6 | Protein S100-A6 | 0.011579 | -0.73098 |
| UGGT1 | UDP-glucose:glycoprotein glucosyltransferase 1 | 0.005255 | -0.73515 |
| WDR13 | WD repeat-containing protein 13 | 0.036249 | -0.74207 |
| ARHGDIA | Rho GDP-dissociation inhibitor 1 | 0.045395 | -0.74455 |
| APOL2 | Apolipoprotein L2 | 0.025062 | -0.74517 |
| OTUB1 | Ubiquitin thioesterase OTUB1 | 0.000506 | -0.74525 |
| GSTP1 | Glutathione S-transferase P | 0.007559 | -0.74625 |
| TCEA1 | Transcription elongation factor A protein 1 | 0.028879 | -0.75037 |
| SUMO2 | Small ubiquitin-related modifier 2 | 0.015779 | -0.75216 |
| CARS1 | Cysteine--tRNA ligase, cytoplasmic | 0.025191 | -0.75259 |
| SOD1 | Superoxide dismutase [Cu-Zn] | 0.038711 | -0.75377 |
| HEXA | Beta-hexosaminidase | 0.005465 | -0.76318 |
| PHACTR4 | Phosphatase and actin regulator 4 | 0.011884 | -0.76322 |
| MVP | Major vault protein | 0.00696 | -0.76482 |
| DNAJC3 | DnaJ homolog subfamily C member 3 | 0.004414 | -0.76739 |
| GPX4 | Phospholipid hydroperoxide glutathione peroxidase | 0.01187 | -0.769 |
| NAGLU | Alpha-N-acetylglucosaminidase | 0.011561 | -0.76955 |
| CCS | Copper chaperone for superoxide dismutase | 0.023725 | -0.77139 |
| TFCP2 | Alpha-globin transcription factor CP2 | 0.014075 | -0.78097 |
| BLVRB | Flavin reductase (NADPH) | 0.032847 | -0.78562 |
| UBLCP1 | Ubiquitin-like domain-containing CTD phosphatase 1 | 0.018824 | -0.80391 |
| GMFB | Glia maturation factor beta | 0.004369 | -0.81092 |
| TPP1 | Tripeptidyl-peptidase 1 | 0.005075 | -0.81383 |
| UCHL3 | Ubiquitin carboxyl-terminal hydrolase isozyme L3 | 0.034757 | -0.81587 |
| HEXB | Beta-hexosaminidase | 0.005763 | -0.81633 |
| AKR1E2 | 1,5-anhydro-D-fructose reductase | 0.025581 | -0.82159 |
|  | cDNA FLJ76436 | 0.021272 | -0.82771 |
| CTSB | Cathepsin B | 0.003472 | -0.83718 |
| PKP3 | Plakophilin-3 | 0.024523 | -0.8458 |
| PGLS | 6-phosphogluconolactonase | 0.021799 | -0.84871 |
| SEC23A | Protein transport protein Sec23A | 0.039729 | -0.85516 |
| PLAU | Urokinase-type plasminogen activator | 0.008343 | -0.86975 |
| P3H1 | Prolyl 3-hydroxylase 1 | 0.007267 | -0.87563 |
| ANP32A | Acidic leucine-rich nuclear phosphoprotein 32 family member A | 0.008569 | -0.87606 |
| HSPA5 | Endoplasmic reticulum chaperone BiP | 0.000357 | -0.87668 |
| SERPINE1 | Plasminogen activator inhibitor 1 | 0.002345 | -0.8767 |
| TXLNA | Alpha-taxilin | 0.006159 | -0.87831 |
| BPNT1 | 3'(2'),5'-bisphosphate nucleotidase 1 | 0.00586 | -0.88063 |
| PSAP | Prosaposin | 0.018483 | -0.8867 |
| GLMP | Glycosylated lysosomal membrane protein | 0.014372 | -0.88987 |
| PLBD2 | Putative phospholipase B-like 2 | 0.035003 | -0.89124 |
| DNM2 | Dynamin-2 | 0.009703 | -0.89348 |
| TRIP11 | Thyroid receptor-interacting protein 11 | 0.028491 | -0.9015 |
| CYCS | Cytochrome c | 0.001345 | -0.91018 |
| PGK1 | Phosphoglycerate kinase 1 | 0.01414 | -0.91982 |
| PIGG | GPI ethanolamine phosphate transferase 2 | 0.003186 | -0.92589 |
| SH3BGRL | Adapter SH3BGRL | 0.034492 | -0.9343 |
| COLGALT1 | Procollagen galactosyltransferase 1 | 0.005135 | -0.94098 |
| RMDN1 | Regulator of microtubule dynamics protein 1 | 0.017853 | -0.94714 |
| CTSD | Cathepsin D | 0.010284 | -0.95614 |
| KRT8 | Keratin, type II cytoskeletal 8 | 0.033508 | -0.95731 |
| PAFAH1B2 | Platelet-activating factor acetylhydrolase IB subunit alpha2 | 0.046417 | -0.97055 |
| APEX1 | DNA-(apurinic or apyrimidinic site) endonuclease | 0.013117 | -0.97059 |
| DCK | Deoxycytidine kinase | 0.040892 | -0.98764 |
| DNAJB11 | DnaJ homolog subfamily B member 11 | 0.024819 | -0.99453 |
| ND3 | NADH-ubiquinone oxidoreductase chain 3 | 0.01906 | -1.0063 |
| UFC1 | Ubiquitin-fold modifier-conjugating enzyme 1 | 0.036342 | -1.01926 |
| VIM | Vimentin | 0.015868 | -1.03827 |
| ITSN2 | Intersectin-2 | 0.046434 | -1.04014 |
| ADD3 | Gamma-adducin | 0.033848 | -1.04158 |
| PLOD2 | procollagen-lysine 5-dioxygenase | 0.007309 | -1.04278 |
| PDIA3 | Protein disulfide-isomerase A3 | 0.001646 | -1.04477 |
| PRPF39 | Pre-mRNA-processing factor 39 | 0.034606 | -1.0698 |
| TTC19 | Tetratricopeptide repeat protein 19, mitochondrial | 0.028854 | -1.07069 |
|  | deoxyribonuclease II (Fragment) | 0.001926 | -1.08631 |
| PSPH | Phosphoserine phosphatase | 0.040924 | -1.08782 |
| SERPINB9 | Serpin B9 | 0.012388 | -1.10137 |
| RNASET2 | Ribonuclease T2 | 0.033095 | -1.12256 |
| DDX58 | Antiviral innate immune response receptor RIG-I | 0.000629 | -1.14392 |
| AK2 | Adenylate kinase 2, mitochondrial | 0.000399 | -1.14693 |
| WDR44 | WD repeat-containing protein 44 | 0.008586 | -1.15925 |
| PRDX4 | Peroxiredoxin-4 | 0.000682 | -1.16255 |
| CBR3 | Carbonyl reductase [NADPH] 3 | 0.027855 | -1.18588 |
| RHBDF2 | Inactive rhomboid protein 2 | 0.04853 | -1.21398 |
| SIL1 | Nucleotide exchange factor SIL1 | 0.017449 | -1.21622 |
| COMMD5 | COMM domain-containing protein 5 | 0.036058 | -1.26027 |
| RCN2 | Reticulocalbin-2 | 0.045965 | -1.26405 |
| GANAB | cDNA FLJ59643, highly similar to Neutral alpha-glucosidase AB | 0.024165 | -1.27208 |
| ARSA | Arylsulfatase A | 0.045861 | -1.27917 |
| HEBP1 | Heme-binding protein 1 | 0.032706 | -1.28607 |
| CALU | Calumenin | 0.003961 | -1.30231 |
| SLC22A18 | Solute carrier family 22 member 18 | 0.020079 | -1.31201 |
| UHRF2 | E3 ubiquitin-protein ligase UHRF2 | 0.039123 | -1.31336 |
| GANAB | Neutral alpha-glucosidase AB | 0.006193 | -1.3617 |
| HSP90B1 | Endoplasmin | 0.006558 | -1.36522 |
| IFNGR1 | Interferon gamma receptor 1 | 0.040973 | -1.37745 |
| CHID1 | Chitinase domain-containing protein 1 | 0.018361 | -1.39886 |
| EEA1 | Early endosome antigen 1 | 0.018827 | -1.45009 |
| FLYWCH2 | FLYWCH family member 2 | 0.000555 | -1.45204 |
| P4HA1 | Prolyl 4-hydroxylase subunit alpha-1 | 0.001758 | -1.45974 |
| PDIA6 | Protein disulfide-isomerase A6 | 0.007683 | -1.49886 |
| MESD | LRP chaperone MESD | 0.023575 | -1.50073 |
| COA7 | Cytochrome c oxidase assembly factor 7 | 0.004026 | -1.50948 |
| PRKCSH | Glucosidase 2 subunit beta | 0.004937 | -1.52669 |
| HLA-F | HLA class I histocompatibility antigen, alpha chain F | 0.028246 | -1.53575 |
| NPC2 | NPC intracellular cholesterol transporter 2 | 0.004567 | -1.53785 |
| HYOU1 | Hypoxia up-regulated protein 1 | 0.000514 | -1.53994 |
| CALR | Calreticulin | 0.003903 | -1.56166 |
| KIF21A | Kinesin-like protein KIF21A | 0.049267 | -1.56229 |
| SIAE | Sialate O-acetylesterase | 0.004738 | -1.56808 |
| RALGAPB | Ral GTPase-activating protein subunit beta | 0.044414 | -1.59015 |
| ABHD14B | Putative protein-lysine deacylase ABHD14B | 0.033678 | -1.5905 |
| MANF | Mesencephalic astrocyte-derived neurotrophic factor | 0.005921 | -1.59094 |
| B3GLCT | Beta-1,3-glucosyltransferase | 0.002905 | -1.61251 |
| BAX | Apoptosis regulator BAX | 0.003655 | -1.62844 |
| HTRA2 | Serine protease HTRA2, mitochondrial | 0.011129 | -1.64366 |
| EOGT | EGF domain-specific O-linked N-acetylglucosamine transferase | 0.007862 | -1.66076 |
| APIP | Methylthioribulose-1-phosphate dehydratase | 0.014043 | -1.67302 |
| RCN1 | Reticulocalbin-1 | 0.003764 | -1.67471 |
| PDIA5 | Protein disulfide-isomerase A5 | 0.00111 | -1.68766 |
| TXNDC5 | Thioredoxin domain-containing protein 5 | 0.000464 | -1.69091 |
| ISG15 | Ubiquitin-like protein ISG15 | 0.013061 | -1.69849 |
| P4HB | Protein disulfide-isomerase | 0.00151 | -1.71395 |
| AP5B1 | AP-5 complex subunit beta-1 | 0.026804 | -1.73778 |
| DTX3L | E3 ubiquitin-protein ligase DTX3L | 0.045461 | -1.75011 |
| POGLUT3 | Protein O-glucosyltransferase 3 | 0.001736 | -1.75093 |
| CNPY3 | Protein canopy homolog 3 | 0.003666 | -1.76387 |
| KIAA1191 | Putative monooxygenase p33MONOX | 0.037031 | -1.79935 |
| POGLUT2 | Protein O-glucosyltransferase 2 | 0.021235 | -1.80126 |
| ERAP2 | Endoplasmic reticulum aminopeptidase 2 | 0.028656 | -1.80151 |
| B4GALT5 | Beta-1,4-galactosyltransferase 5 | 0.047535 | -1.80835 |
| SERPINH1 | Serpin H1 | 0.000238 | -1.83661 |
| P4HA2 | Prolyl 4-hydroxylase subunit alpha-2 | 0.02314 | -1.85142 |
| MSRB3 | Peptide-methionine (R)-S-oxide reductase (Fragment) | 0.003837 | -1.87663 |
| COL6A1 | Collagen alpha-1(VI) chain | 0.009883 | -1.88257 |
| UBL7 | Ubiquitin-like protein 7 | 0.02477 | -1.9047 |
| PDIA4 | Protein disulfide-isomerase A4 | 0.000636 | -1.92073 |
| ERAP1 | Endoplasmic reticulum aminopeptidase 1 | 0.000475 | -1.92721 |
| CHCHD5 | Coiled-coil-helix-coiled-coil-helix domain-containing protein 5 | 0.038169 | -1.94828 |
| SARNP | SAP domain-containing ribonucleoprotein | 0.024703 | -1.95128 |
| TP53RK | EKC/KEOPS complex subunit TP53RK | 0.006128 | -1.9793 |
| POFUT1 | GDP-fucose protein O-fucosyltransferase 1 | 0.004483 | -1.98745 |
| POFUT2 | GDP-fucose protein O-fucosyltransferase 2 | 0.016255 | -2.00868 |
| OARD1 | ADP-ribose glycohydrolase OARD1 | 0.043931 | -2.02519 |
| ERP29 | Endoplasmic reticulum resident protein 29 | 0.001449 | -2.04398 |
| BTD | Biotinidase | 0.010539 | -2.05414 |
| ANKRD12 | Ankyrin repeat domain-containing protein 12 | 0.001776 | -2.0586 |
| SUMF2 | Inactive C-alpha-formylglycine-generating enzyme 2 | 0.001141 | -2.07479 |
| SUMF1 | Formylglycine-generating enzyme | 0.034134 | -2.07712 |
| IDUA | Alpha-L-iduronidase | 0.00938 | -2.08254 |
| RIPK2 | Receptor-interacting serine/threonine-protein kinase 2 | 0.039333 | -2.08388 |
| FKBP9 | Peptidyl-prolyl cis-trans isomerase FKBP9 | 0.00457 | -2.08842 |
| TXNDC12 | Thioredoxin domain-containing protein 12 | 0.01417 | -2.09274 |
| FAM114A1 | Protein NOXP20 | 0.018235 | -2.12034 |
| ERO1A | ERO1-like protein alpha | 0.000655 | -2.17112 |
| H6PD | GDH/6PGL endoplasmic bifunctional protein | 0.019735 | -2.21876 |
| DIABLO | Diablo IAP-binding mitochondrial protein | 0.016339 | -2.24284 |
| DES | Desmin | 0.009517 | -2.24763 |
| ERO1B | ERO1-like protein beta | 0.049318 | -2.26996 |
| MYDGF | Myeloid-derived growth factor | 0.00038 | -2.2958 |
| KRT13 | Keratin, type I cytoskeletal 13 | 0.003279 | -2.34824 |
| CNPY2 | Protein canopy homolog 2 | 0.002509 | -2.40322 |
| IFI35 | Interferon-induced 35 kDa protein | 0.019077 | -2.41805 |
| FKBP10 | Peptidyl-prolyl cis-trans isomerase FKBP10 | 0.011936 | -2.42461 |
| TRIM2 | Tripartite motif-containing protein 2 | 0.018573 | -2.45192 |
| FKBP2 | Peptidyl-prolyl cis-trans isomerase FKBP2 | 0.003629 | -2.47448 |
| YTHDF1 | YTH domain-containing family protein 1 | 0.012913 | -2.49952 |
| P4HTM | Transmembrane prolyl 4-hydroxylase | 0.018182 | -2.59585 |
| CRK | Adapter molecule crk | 0.013527 | -2.5979 |
| GGCT | Gamma-glutamylcyclotransferase | 0.001765 | -2.63158 |
| MACO1 | Macoilin | 0.004336 | -2.69016 |
| CCDC88A | Girdin | 0.035487 | -2.71469 |
| FAM136A | Protein FAM136A | 0.021839 | -2.75768 |
| SUOX | Sulfite oxidase, mitochondrial | 0.022718 | -2.76727 |
| PPIC | Peptidyl-prolyl cis-trans isomerase C | 0.006375 | -2.88949 |
| IFIT1 | Interferon-induced protein with tetratricopeptide repeats 1 | 0.00683 | -2.91872 |
| CRELD2 | Protein disulfide isomerase CRELD2 | 0.007552 | -3.01519 |
| FHL3 | Four and a half LIM domains protein 3 | 0.007777 | -3.03153 |
| GRIA1 | Glutamate receptor 1 | 0.003254 | -3.09533 |
| CALU | Calumenin, isoform CRA_a | 0.01418 | -3.13661 |
| ENOPH1 | Enolase-phosphatase E1 | 0.004774 | -3.23129 |
| MIX23 | Protein MIX23 | 0.005537 | -3.30314 |
| TXNDC16 | Thioredoxin domain-containing protein 16 | 0.013377 | -3.36055 |
|  | cDNA FLJ56447, highly similar to Dynamin-2 | 0.018938 | -3.58 |
| MANBA | Beta-mannosidase | 0.007203 | -3.65432 |
| P4HA1 | procollagen-proline 4-dioxygenase (Fragment) | 0.033344 | -3.79713 |
| IRF9 | Interferon regulatory factor 9 | 0.049991 | -4.10625 |
| ATF3 | Cyclic AMP-dependent transcription factor ATF-3 | 0.013061 | -4.80024 |
| REPS2 | RalBP1-associated Eps domain-containing protein 2 | 0.000428 | -5.25089 |
| RNF25 | E3 ubiquitin-protein ligase RNF25 | 0.017319 | -5.62688 |
| IFIT3 | Interferon-induced protein with tetratricopeptide repeats 3 | 9.04E-07 | -5.67401 |
| TRIAP1 | TP53-regulated inhibitor of apoptosis 1 | 0.02492 | -5.69279 |
| OAS1 | 2'-5'-oligoadenylate synthase 1 | 0.045746 | -5.90219 |
| ASS1 | Argininosuccinate synthase | 0.001598 | -6.10895 |
| LGALSL | Galectin-related protein | 0.025414 | -6.62655 |
| ITIH1 | Inter-alpha-trypsin inhibitor heavy chain H1 | 0.01213 | -6.93277 |
| WDR45B | WD repeat domain phosphoinositide-interacting protein 3 | 0.016785 | -8.06515 |

**Table S2. Sequences for siRNA in this study**

| **Oligonucleotides** | **Sequences** |
| --- | --- |
| si*YAP1*-#1 | CCACCAAGCUAGAUAAAGATT |
| si*YAP1*-#2 | GGUCAGAGAUACUUCUUAATT |
| si*YAP1*-#3 | UCAGAGUGCUCCAGUGACCTT |
| si*p130Cas* | AGAAGGAGCUGCUGGAAAATT |
| si*ITGB1* | CCACAGACAUUUACAUUAATT |
| si*DDR1* | GGAGAUGGAGUUUGAGUUTT |
| si*PIEZO1* | GGAACAGGCAGGACAGCUATT |
| si*CD44* | CCACAAUGGCCCAGAUGGTT |
| *Control siRNAs* | UUCUCCGAACGUGUCACGUTT |

**Table S3. The sequences for primers used in this study**

| **For real-time qPCR** | | |  | |
| --- | --- | --- | --- | --- |
| **Gene** | **Forward primer (5' -> 3')** | | **Reverse primer (5' -> 3')** | |
| *NGF* | TGTGGGTTGGGGATAAGACCA | | GCTGTCAACGGGATTTGGGT | |
| *BDNF* | CTACGAGACCAAGTGCAATCC | | AATCGCCAGCCAATTCTCTTT | |
| *NTF3* | CGTGGTGGCGAACAGAACAT | | GGCCGATGACTTGTCGGTC | |
| *NTF4* | CTGTGTGCGATGCAGTCAGT | | TGCAGCGGGTTTCAAAGAAGT | |
| *SEMA3A* | AGACTCACTTGTACGCCTGTG | | CCCAAGAGTTCGGAAGATAGCAA | |
| *SEMA3B* | ACGTCCAAGTCTCCGAACAGA | | GCCGTCTCACGAAAGAAGAAGT | |
| *SEMA3C* | TAACCAAGAGGAATGCGGTCA | | TGCTCCTGTTATTGTCAGTCAGT | |
| *SEMA3D* | GCAAAGGAACGGGTGGAATTA | | TCTGCCAGACTCCAAATTATGTG | |
| *SEMA3E* | GTTTGCTGGACTCTACAGTGAC | | CTTTCAACAGACGCTCATCGT | |
| *SEMA3F* | CCACAGCGCATCGAGGAAT | | CATGGGGTTGTAGGCACCTG | |
| *SEMA3G* | CAGAGGATGGGACCTACGATG | | GTTGGCACCTTAAACACCTGG | |
| *SLIT1* | GCCTGGAACTCAATGGCAAC | | CTGGTTTCGGTTCAGTCGCA | |
| *SLIT2* | GCGAAGCTATACAGGCTTGAT | | TGCAGTCGAAAAGTCCTAAGTTT | |
| *SLIT3* | CGGCATCACCGATGTGAAGAA | | AGGCGCAGAGTTCGGATCT | |
| *NTN1* | ACAACCCGCACAACCTGAC | | GGGACAGTGTGAGCGTGAC | |
| *NTN3* | GCCGTCGTCCCTTACTCCTA | | TGAGGCATGTCCATTGCACTT | |
| *NTN4* | GTACTTTGCGACTAACTGCTCC | | TCCAGTGCATGGAAAAGGACT | |
| *NTN5* | TGCCGGTTCAACTCTGAGC | | CTGTTGCCCCAATAGGGTGG | |
| *EFNA1* | TCAGGCCCATGACAATCCAC | | GTGACCGATGCTATGTAGAACC | |
| *EFNA2* | TACGCCGTCTACTGGAACC | | GAGCCTCGTACAGGGTCTC | |
| *EFNA3* | CATGCGGTGTACTGGAACAG | | AGATAGTCGTTCACGTTCACCT | |
| *EFNA4* | CTCCGCCACGTAGTCTACTG | | TACAAAGCAAACGTCTCGGGG | |
| *EFNA5* | CTACATGGTGAACTTTGATGGCT | | GAGGCCGGTTACATTCCCA | |
| *EFNB1* | TGGAGCCCGTATCCTGGAG | | TTGGGGTCGAGAACTGTGCTA | |
| *EFNB2* | TATGCAGAACTGCGATTTCCAA | | TGGGTATAGTACCAGTCCTTGTC | |
| *EFNB3* | CGCTCGCACCACGATTACTA | | GCCTCTGGTTAGGCACACA | |
| *CTGF* | AAAAGTGCATCCGTACTCCCA | | CCGTCGGTACATACTCCACAG | |
| *CYR61* | GGTCAAAGTTACCGGGCAGT | | GGAGGCATCGAATCCCAGC | |
| *ITGB1* | CCTACTTCTGCACGATGTGATG | | CCTTTGCTACGGTTGGTTACATT | |
| *DDR1* | CCGACTGGTTCGCTTCTACC | | CGGTGTAAGACAGGAGTCCATC | |
| *PIEZO1* | GGACTCTCGCTGGTCTACCT | | GGGCACAATATGCAGGCAGA | |
| *CD44* | CTGCCGCTTTGCAGGTGTA | | CATTGTGGGCAAGGTGCTATT | |
| *ACTB* | CATGTACGTTGCTATCCAGGC | | CTCCTTAATGTCACGCACGAT | |
| *Ngf* | CCAGTGAAATTAGGCTCCCTG | | CCTTGGCAAAACCTTTATTGGG | |
| *Ctgf* | AAAAGTGCATCCGTACTCCCA | | CCGTCGGTACATACTCCACAG | |
| *Cyr61* | CTGCGCTAAACAACTCAACGA | | GCAGATCCCTTTCAGAGCGG | |
| *Ankrd1* | CGTGGAGGAAACCTGGATGTT | | GTGCTGAGCAACTTATCTCGG | |
| *Actb* | GGCTGTATTCCCCTCCATCG | | CCAGTTGGTAACAATGCCATGT | |
| **For ChIP-qPCR** | | | |  |
| **Gene** | | **Forward primer (5' -> 3')** | | **Reverse primer (5' -> 3')** |
| *BDNF* | GCACACACACACACACACAC | | GCACGCCCACCTGATAGTAT | |
| *CTGF* | TGTGCCAGCTTTTTCAGACG | | TGAGCTGAATGGAGTCCTACACA | |
| *HBB* | GCTTCTGACACAACTGTGTTCACTAGC | | CACCAACTTCATCCACGTTCACC | |
